# A candidate causal variant underlying both enhanced cognitive performance and increased risk of bipolar disorder

**DOI:** 10.1101/580258

**Authors:** Susan Q. Shen, Jeong Sook Kim-Han, Lin Cheng, Duo Xu, Omer Gokcumen, Andrew E.O. Hughes, Connie A. Myers, Joseph C. Corbo

## Abstract

Bipolar disorder is a highly heritable mental illness, but the relevant genetic variants and molecular mechanisms are largely unknown. Recent GWASs have identified an intergenic region associated with both cognitive performance and bipolar disorder. This region contains dozens of putative fetal brain-specific enhancers and is located ∼0.7 Mb upstream of the neuronal transcription factor *POU3F2*. We identified a candidate causal variant, rs77910749, that falls within a highly conserved putative enhancer, LC1. This human-specific variant is a single-base deletion in a PAX6 binding site and is predicted to be functional. We hypothesized that rs77910749 alters LC1 activity and hence *POU3F2* expression during neurodevelopment. Indeed, transgenic reporter mice demonstrated LC1 activity in the developing cerebral cortex and amygdala. Furthermore, *ex vivo* reporter assays in embryonic mouse brain and human iPSC-derived cerebral organoids revealed increased enhancer activity conferred by the variant. To probe the *in vivo* function of LC1, we deleted the orthologous mouse region, which resulted in amygdala-specific changes in *Pou3f2* expression. Lastly, ‘humanized’ rs77910749 knock-in mice displayed behavioral defects in sensory gating, an amygdala-dependent endophenotype seen in patients with bipolar disorder. Our study suggests a molecular mechanism underlying the long-speculated link between enhanced cognitive performance and neuropsychiatric disease.

## INTRODUCTION

Genome-wide association studies (GWAS’s) have identified thousands of disease-associated non-coding regions, but pinpointing the underlying ‘causal variants’ is challenging^1^. For neuropsychiatric diseases, this is particularly challenging due to the multiple layers of biological organization between the variant and behavioral phenotype, and the lack of appropriate model systems. Bipolar disorder (BD) is a neuropsychiatric illness characterized by altered mood, classically with episodes of mania and depression^2^. It affects ∼1% of the world population and has high morbidity and mortality^3^. While BD is highly (∼80%) heritable, the relevant genes and pathways are largely unknown, although the amygdala and prefrontal cortex are strongly implicated^2,4,5^. Fascinatingly, BD is associated with heightened creativity, highlighting the long-speculated link between ‘madness’ and ‘genius’^6,7^.

Recently, three GWAS’s of BD implicated an intergenic region at the *MIR2113/POU3F2* locus in 6q16.1^8-10^. In parallel, multiple GWAS’s of educational attainment and cognitive performance pinpointed the same locus^11-13^. The ‘lead SNPs’ (i.e., with the lowest p-values) in the studies of BD and in the studies of cognitive performance are in strong linkage disequilibrium (LD), suggesting a common underlying causal variant. Since the nearest protein-coding gene, *POU3F2*, is located ∼0.7 Mb away, we hypothesized that the underlying causal variant affects the activity of a non-coding *cis*-regulatory element (CRE, i.e., enhancer or silencer) that regulates *POU3F2*.

*POU3F2* (*BRN-2*) is a transcription factor (TF) that is widely expressed in the developing brain. *POU3F2* and *POU3F3* (*BRN-1*) jointly regulate the neurogenesis, maturation, and migration of upper-layer cortical neurons^14-16^. The transcriptional targets of *POU3F2* likely include *FOXP2*, a TF involved in speech and vocalization^17^. Furthermore, overexpression of *POU3F2* facilitates direct reprogramming of fibroblasts and astrocytes into neurons^18,19^. In mice, both increased and decreased levels of *Pou3f2* are associated with altered neuronal fate, and a specific mutation in *Pou3f2* affects cognitive function^15,20,21^. In humans, deletions encompassing *POU3F2*, and a missense mutation in *POU3F2*, are associated with intellectual disability^22,23^. Thus, *cis*-regulatory changes that alter *POU3F2* expression levels could similarly perturb brain development, affecting cognition and other neuropsychiatric traits.

Here, we identify and investigate a candidate causal variant, rs77910749, which falls within LC1, a putative brain enhancer located upstream of *POU3F2* in the intergenic region implicated by GWAS’s of BD and cognitive performance. We create transgenic reporter mice to interrogate enhancer activity in neurodevelopment. We also implement a multiplex reporter assay, CRE-seq, in developing mouse brain and human induced pluripotent stem cell (iPSC)-derived cerebral organoids to quantify the effect of rs77910749 on enhancer activity. Finally, we use CRISPR-Cas9 to generate LC1 knockout mice and ‘humanized’ rs77910749 knock-in mice, thereby establishing models for behavioral assays. We demonstrate evidence for a BD-related behavioral phenotype in rs77910749 knock-in mice. Together, our studies provide molecular evidence for a mechanistic link between cognitive performance and BD.

## RESULTS

### The *MIR2113/POU3F2* locus harbors non-coding variants associated with both enhanced cognitive performance and elevated risk of BD

To assess whether genetic markers associated with cognitive performance and BD might have a shared biological origin, we first compared lead SNPs across studies. Two GWAS’s of educational attainment in Caucasians identified a genome-wide significant signal at the *MIR2113/POU3F2* intergenic region (lead SNP rs9320913)^11,24^. Educational attainment was later shown to be a proxy phenotype for cognitive performance^25,26^. In two GWAS meta-analyses for cognitive ability, rs9320913 was not directly genotyped, but the proxy variant rs1906252 (r^2^ = 0.96 with rs9320913) was associated with increased general cognitive ability^12,27^. Similarly, another GWAS meta-analysis found that the proxy variant rs10457441 (r^2^ = 0.91 with rs9320913) is associated with greater general cognitive ability^13^. Lastly, a study of 1.1 million individuals identified this locus as one of the top hits for educational attainment, cognitive performance, and highest math class completed^28^. Thus, multiple studies demonstrated an association between cognition and variants at this locus, pointing to a single causal haplotype (Table S1).

Around the same time, a GWAS of 9,747 Caucasian BD patients and 14,278 controls identified a novel risk locus at the same intergenic region^8^. The lead SNP, rs12202969, was associated with ∼10-20% increased risk for BD. Another GWAS study of BD confirmed this signal^10^. A third GWAS of 9,784 Caucasian BD patients and 30,471 controls pinpointed the proxy variant rs1487441 (r^2^ = 0.98 with rs12202969)^9^. We observed that the two BD GWAS lead SNPs (rs12202969 and rs1487441) were in high LD with the lead SNPs in the GWAS’s of educational attainment and cognition (rs9320913, rs1906252, and rs104757441) (pairwise r^2^ = 0.92-0.99), suggesting a shared genetic basis for cognitive performance and BD (Table S1). Intriguingly, the variants associated with enhanced cognitive performance were associated with *increased* BD risk, consistent with the finding that children with higher IQs are at higher risk for developing manic features^29,30^.

### The candidate causal variant rs77910749 is a human-specific non-coding variant

To identify candidate causal variants, we surveyed the epigenomic landscape of the 0.5 Mb region (Chr6:98,300,000-98,800,000 Mb in hg19) identified by the GWAS’s (Fig. 1A, yellow box). This LD block contains dozens of human fetal brain-specific DNase-seq peaks, which are open chromatin regions that demarcate putative CREs^31^. We then focused on the ∼60 kb region of highest LD, which contains all five lead SNPs, SNPs: rs9320913, rs1906252, rs10457441, rs12202969, and rs1487441 (Fig. 1A, purple box). Within this region, we identified six fetal brain-specific DNase I hypersensitive sites (DHSs), termed LC0 through LC5 (the ‘local cluster’) (Fig. 1B). While none of the lead SNPs fell within fetal brain DHSs, four variants in LD with rs9320913 (r^2^ > 0.2 based on HaploReg v4.1 using 1000 Genomes Phase 1 for Europeans^32,33^) fell within fetal brain DHSs in the local cluster (Fig. 1C top panel, blue font): rs77910749 in LC1, rs13208578 in LC2, rs12204181 in LC4, and rs17814604 in LC5.

**Figure 1.**
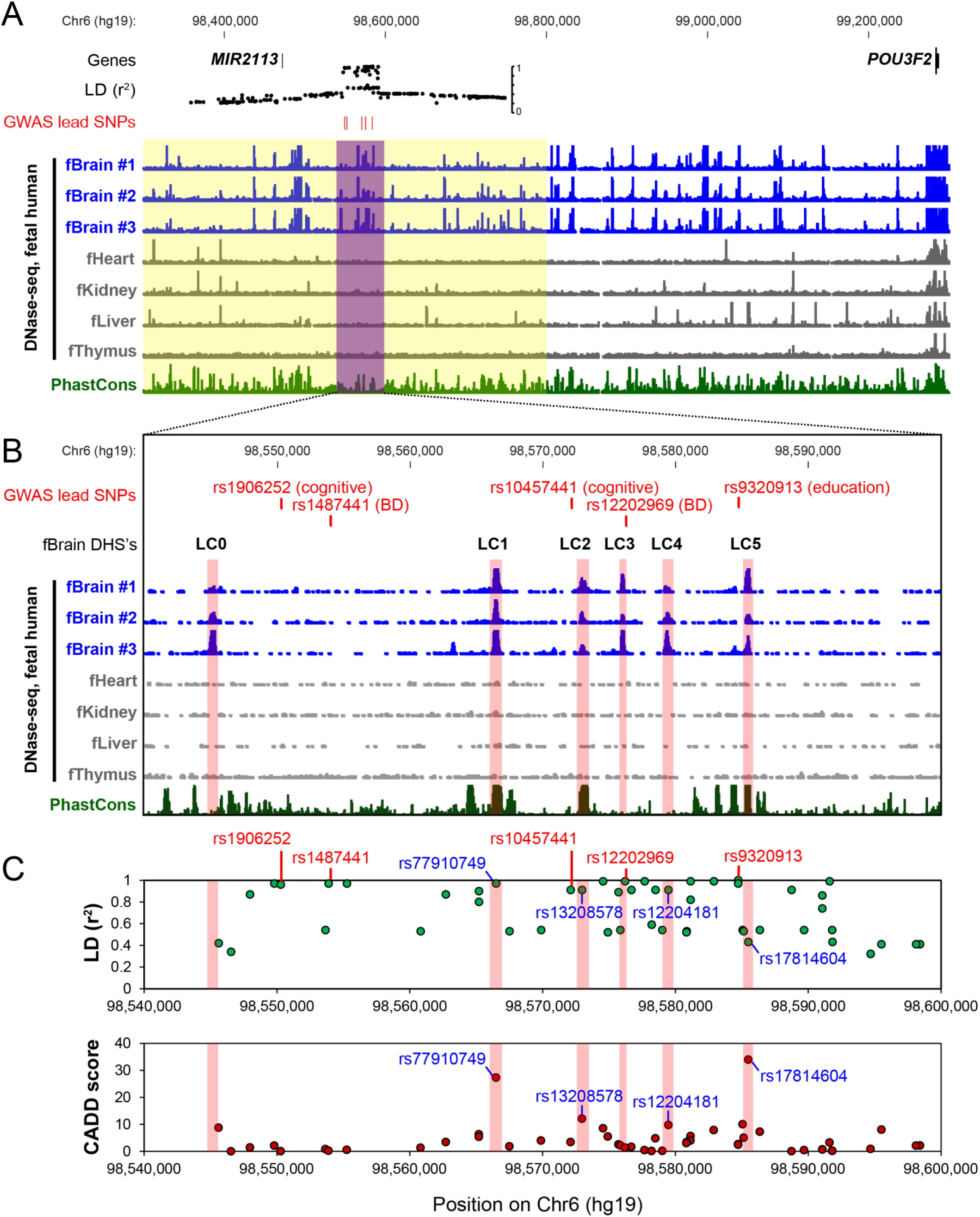
Prioritization of candidate variants at 6q16.1 associated with higher educational attainment, enhanced cognitive performance, and elevated risk for bipolar disorder. (A) Genomic context (hg19, 1 Mb window) of the intergenic locus implicated in GWAS’s of educational attainment, cognition, and BD. The 0.5 Mb region identified by these studies (yellow box) contains a ∼60 kb ‘local cluster’ region (purple box) with the highest LD. All variants in LD with rs9320913 (r^2^ > 0.2) are shown. The nearest protein-coding gene, *POU3F2*, is ∼0.7 Mb away. DNase-seq data from three human fetal brains and four other human fetal tissues are shown^31^. PhastCons depict 100-way vertebrate conservation^73^. The UCSC Genome Browser was used for visualization^70^. (B) Enlarged view the 60 kb ‘local cluster’. Note the fetal brain (fBrain) DHSs (LC0 to LC5, pink box). Lead SNPs (red font): rs9320913 for educational attainment^11,24^, rs1906252 for cognitive performance^12^, rs10457441 for cognitive performance^13^, rs12202969 for BD^8^, and rs1487441 for BD^9^. (C) Variants within the local cluster that are in LD with rs9320913 (as defined by r^2^ > 0.2). Note the five lead SNPs (red font) and four variants that fall within LC1-5 (blue font). The r^2^ values (green dots) and Phred-scaled CADD scores are shown (red dots)^35^.

Since phylogenetic conservation often reflects functionality, we hypothesized that the causal variant fell within a phylogenetically conserved region. As LC4 exhibits low conservation, rs12204181 was deemed a less likely candidate. LC2 is highly conserved, but the derived allele corresponding to rs13208578 is present in multiple vertebrate species, including primates, suggesting that it is well-tolerated (Fig. S1A). Furthermore, LC2 did not exhibit enhancer activity at E11.5 in a transgenic reporter mouse^34^. Thus, rs77910749 and rs17814604 were the top candidates, a finding corroborated by CADD, a bioinformatic tool that predicts variant deleteriousness. CADD ranked rs77910749 and rs17814604 respectively in the top 0.2% and 0.04% of all possible human variants for predicted deleteriousness (Fig. 1C, bottom)^35^.

Next, we examined rs77910749 and rs17814604. Phylogenetic analyses demonstrated that rs17814604 is a newer and rarer allele that emerged from a haplotype already containing rs77910749 (Fig. S3A and SI). In particular, the allele frequency of rs17814604 is only 0.2% in East Asians (1000 Genomes Phase 3)^36^. A study of 342 Han Chinese found a significant association between rs12202969 (r^2^ = 0.96 with rs9320913 in Han Chinese) and math ability^37^. Since rs17814604 is nearly absent in Han Chinese, it is unlikely that rs17814604 is the underlying causal variant. By contrast, rs77910749 is relatively common worldwide (Fig. S3B and SI), with an allele frequency of 51% in Europeans (1000 Genomes Phase 3)^36^.

Inspection of rs77910749 revealed that it is a single base pair deletion of a ‘T’ in a stretch of ∼100 bases that are nearly perfectly conserved among vertebrates down to coelacanth (Fig. S1B). Furthermore, although we did not find evidence of a traditional selective sweep, rs77910749 appears to be a human-specific variant (Fig. S4, Fig. S5, and SI). Thus, rs77910749 is a common, human-specific variant at an evolutionarily conserved nucleotide, which is hypothesized to be associated with both enhanced cognitive performance and increased risk of BD.

### rs77910749 falls within a putative developmental brain enhancer

Since LC1 is highly conserved, we examined the orthologous region in other vertebrate genomes. We found that LC1 is located between *Mir2113* and *Pou3f2* in multiple vertebrate genomes, suggesting that LC1 is part of a genomic regulatory block whose conserved synteny reflects functionality^38^. This was corroborated by analysis of available Hi-C data (Fig. S6 and SI).

Next, we examined the epigenomic landscape of LC1. Published DNase-seq data across multiple mouse tissues^39^ demonstrated that LC1 is a region of open chromatin in the developing mouse brain, with a strong signal at E14.5 and greatly diminished signal by E18.5 and adulthood (Fig. 2A). ChIP-seq for two enhancer marks, p300 and H3K27ac^40,41^, suggested that LC1 is an active brain enhancer at E14.5 (Fig. 2A). Moreover, DNase-seq of mouse retina showed that LC1 is open in the early postnatal period but subsequently closes, suggesting that LC1 has a role in neurogenesis in both brain and retina (Fig. 2A)^42^. Human methylation data support the notion that LC1 is active in neural progenitors (Fig. S7). Interestingly, rs77910749 creates a novel CpG site in LC1 with potential for methylation (Fig. S8).

**Figure 2.**
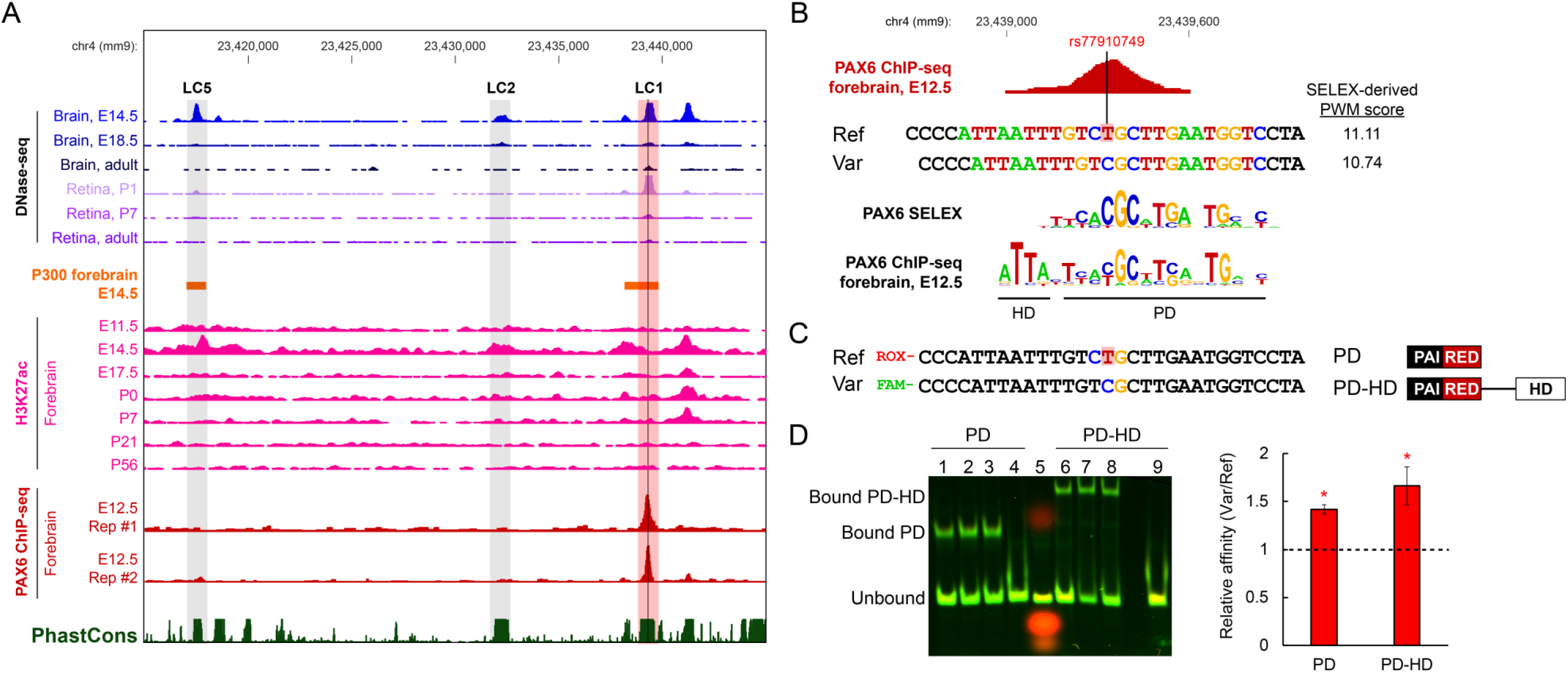
The candidate causal variant, rs77910749, affects PAX6 binding. (A) The 30 kb ‘local cluster’ in the mouse genome (mm9). Mouse LC1 overlaps with E14.5 brain and P1 retina DNase-seq^39^, E14.5 forebrain p300 ChIP-seq (orange)^40^, E14.5 forebrain H3K27ac ChIP-seq (pink)^41^, and E12.5 forebrain PAX6 ChIP-seq (dark red; two replicates are shown)^47^. The orthologous position of human-specific rs77910749 (black vertical line in LC1) falls within the PAX6 ChIP-seq peak. (B) Comparison of the reference sequence (‘Ref’), sequence with rs77910749 (‘Var’), and PAX6 consensus motifs. The position of rs77910749 is indicated (red highlighted ‘T’). The reference sequence is conserved between mouse and human, and the minus strand is shown. Motifs were scored using SELEX-derived position weight matrices (PWMs) for PAX6 protein with PD-HD domains^48^. The logo was generated in enoLOGOS^74^. The E12.5 PAX6 ChIP-seq motif was derived from^47^. (C) Quantitative EMSA assay. Reference and variant probes of equal lengths were fluorescently labeled. A second probe set (not shown) in which fluorescent labels were reversed yielded similar results. PAI and RED domains form the PAX6 paired domain (PD), which is separated by the homeodomain (HD) with a linker. (D) Left, representative EMSA gel. Lanes 1-3, PD binding reaction. Lane 4, cold competition reaction with PD. Lane 5, probes and marker dye only (no protein). Lanes 6-8, PD-HD binding reaction. Lane 9, cold competition reaction with PD-HD. For cold competition reactions, 500-fold molar excess of unlabeled vs. labeled probe was used. Right, quantification of EMSA results. Bound and unbound fractions were quantified, and relative binding affinity was calculated^49^. Error bars indicate SEM across six binding reactions (three each from the two probe sets with reversed fluorophores). Black dotted horizontal line: null hypothesis that rs77910749 has no effect on affinity. P < 0.05 for PD and PD-HD (95% confidence interval).

### rs77910749 increases the affinity of a PAX6 binding site within LC1

Because *cis*-regulatory variants can alter enhancer activity by disrupting TF binding, we searched for predicted TF motifs within LC1 using FIMO (see SI)^43^. We found that rs77910749 falls within a predicted binding site for PAX6 (Fig. 2A). PAX6 is a TF with multiple critical roles in brain development and likely directly regulates *Pou3f2*^15,44-46^. Published PAX6 ChIP-seq data from E12.5 mouse forebrain revealed that LC1 is strongly bound by PAX6 *in vivo* (80^th^ ranked peak out of 3,536 peaks) and the only prominent peak in the region (Fig. 2A and 2B)^47^.

Based on *in vitro* binding preferences from SELEX^48^, rs77910749 is predicted to slightly (∼3%) decrease PAX6 binding affinity (Fig. 2B). To directly measure the effect of rs77910749 on binding affinity, we expressed and purified the paired domain (PD) of PAX6 and conducted quantitative electrophoretic mobility shift assays (EMSAs) with fluorescently labeled DNA probes^49^ (Fig. 2C). Since PAX6 has a homeodomain (HD) that can interact with PD, we also expressed PD with HD (‘PD-HD’ protein). We found that both PD alone and PD-HD can bind both the wild-type sequence (‘Ref’) and the rs77910749-containing sequence (‘Var’), as demonstrated by specific gel shifts. However, PD5a (an isoform of canonical PAX6) cannot bind to either the reference or variant sequence (Fig. S9 and SI).

Upon quantification of probe binding, we found that rs77910749 confers ∼40% increased binding affinity for PD (95% confidence interval [CI]: 1.32-1.52 fold higher affinity) and ∼60% increased binding affinity for PD-HD (95% CI: 1.28-2.05 fold higher affinity) (Fig. 2D). Thus, contrary to *in silico* predictions, rs77910749 confers a significant increase in PAX6 binding affinity.

### Transgenic reporter mice show evidence of LC1 enhancer activity in the developing central nervous system (CNS)

To test whether LC1 is a *bona fide* enhancer and to investigate its spatiotemporal activity pattern, we created transgenic reporter mice carrying human LC1 (∼1 kb fragment) cloned upstream of the minimal *Hsp68* promoter and LacZ (Fig. 3A and SI)^50^. Since the mouse DNase-seq signal for LC1 is strongest at E14.5, we screened ‘transient’ transgenic embryos at E14.5 (i.e., embryos were F0’s and represented independent transgenesis events). Among the seven embryos that were genotypically positive for LacZ, five showed LacZ expression (Fig. 3B): in cerebral cortex (lines #1, 4, 5), amygdala (lines #1, 2, and 3), and skin (line #5).

**Figure 3.**
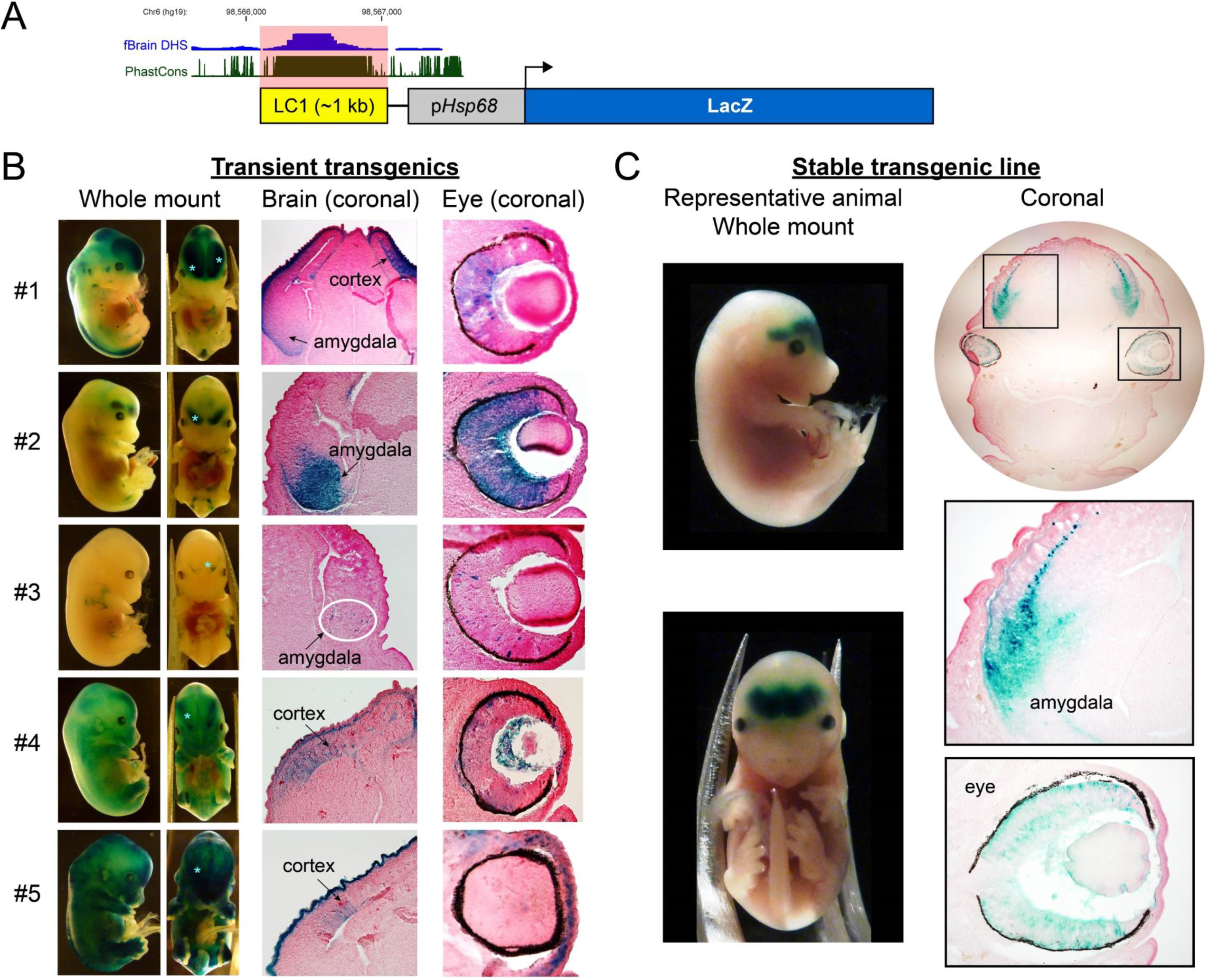
Transgenic reporter mice show evidence of LC1 activity in the developing CNS. Mice were generated that carried a reporter construct for wild-type human LC1 (951 bp fragment) on the *Hsp68* promoter, driving the expression of LacZ, which stains blue with X-gal^50^. (A) Schematic of the reporter construct (drawn to scale). (B) Transient transgenic embryos. Of seven genotypically positive embryos, five (#1-5 shown here) exhibited LacZ staining. Each mouse represents an independent integration event. Whole mount images of lateral and frontal views; light blue asterisks in the frontal views denote the approximate location of annotated regions in the brain coronal sections. The entire head was embedded and cryosectioned. For the brain coronal image of embryo #3, the white oval encircles sparse LacZ-expressing cells. Magnified images of the eye are also shown. (C) Representative embryo from a stable transgenic line. Of three genotypically positive stable transgenic lines, only this line exhibited LacZ staining. Multiple embryos from this stable line had essentially identical LacZ staining patterns, as expected. Lateral and frontal views are shown. Coronal section of head and corresponding enlarged images of the amygdala and eye are shown. Sections were counterstained with Nuclear Fast Red.

We also created three independent stable lines (in which F0 transgenics were outcrossed to generate F1’s). Two stable lines showed essentially no enhancer activity in multiple genotypically positive E14.5 embryos. The third stable line showed consistent LacZ expression in the developing amygdala (Fig. 3C). Thus, overall, 6/10 transgenic lines showed LacZ expression in the developing brain, with 4/6 in the developing amygdala and 3/6 in the developing cortex. Additionally, 5/6 expressed LacZ in the developing retina (Fig. 3), consistent with retinal DNase-seq data (Fig. 2A). Together, these data indicate that LC1 is transcriptionally active in the developing amygdala, cerebral cortex, and retina. PAX6 has known roles in the development of all three regions^51-53^, whereas POU3F2 has known roles in the cerebral cortex and retina, and a suggested role in the amygdala^14,16,54^. Some variability in expression was seen among the reporter lines, possibly due to insertion site effects^55^.

### rs77910749 increases LC1 enhancer activity in mouse cerebral cortex and human cerebral organoids

To quantitatively assess whether rs77910749 alters the enhancer activity of LC1, we utilized a multiplexed plasmid reporter assay, CRE-seq^56^. In CRE-seq, a library of barcoded reporter constructs is introduced into cells, and the resulting expressed transcripts are quantified by RNA-seq. We previously used CRE-seq to assay thousands of CREs in postnatal mouse retina and adult cerebral cortex^56,57^. Here, we assayed a smaller pool of constructs with greater coverage and depth. We created three types of constructs: wild-type LC1 (‘Ref’), LC1 with rs77910749 (‘Var’), and a promoter-only control. To increase the sensitivity of our assay, the enhancers were synthesized as multimers (Fig. 4A and SI). For each of the three construct types, twenty barcoded constructs were created, for a total of sixty barcoded constructs in the GFP reporter library.

**Figure 4.**
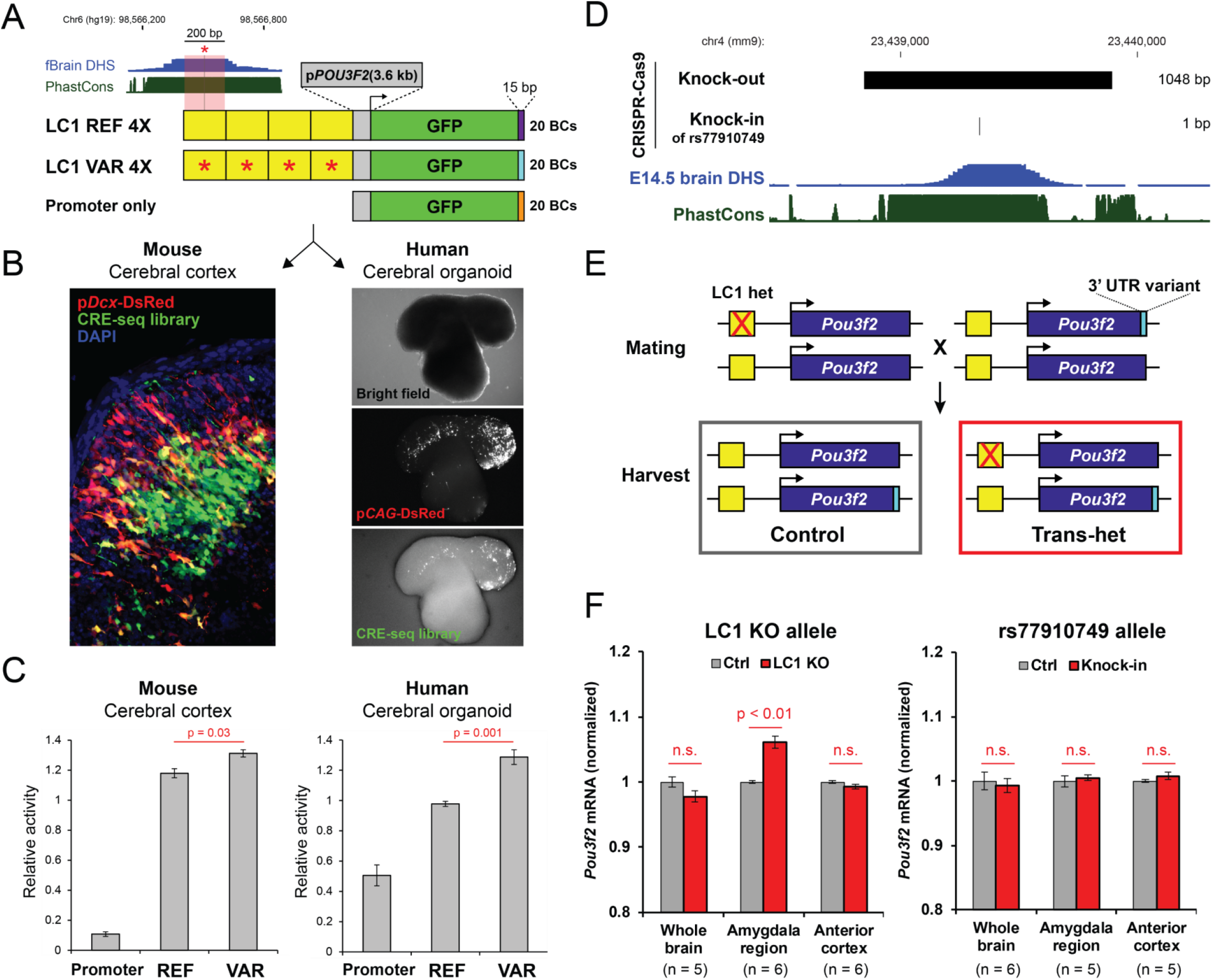
The variant rs77910749 increases enhancer activity in *ex vivo* mouse brain and human iPSC-derived cerebral organoids, and knockout of LC1 alters *Pou3f2* expression in the developing amygdala *in vivo*. (A) Schematic of the CRE-seq experiment. Multimers (4X) of the central 200 bp of human LC1 were cloned upstream of a 3.6 kb *POU3F2* (human) promoter fragment and GFP with unique 15 bp barcodes (BCs) in the 3’ UTR. ‘REF’ indicates wild-type sequence and ‘VAR’ indicates the presence of rs77910749 (red asterisk), whose position is shown by the black vertical line. Twenty barcoded constructs were generated for each of REF, VAR, and promoter-only. (B) Library delivery. Left: E12.5 mouse cerebral cortex was electroporated and harvested after 2 days in culture. A vibratome section (100 μm thickness) shows library GFP expression in the deeper layers of the cerebral cortex. The co-electroporated control construct, p*Dcx*-DsRed, is expressed in post-mitotic migrating neurons^59^. DAPI is a nuclear counterstain. Right: Human iPSC-derived cerebral organoids were electroporated and harvested after 7 days in culture. A representative live image of an electroporated organoid shows library GFP expression. The co-electroporated control construct, p*CAG*-DsRed, marks electroporated cells. (C) Quantification of *cis*-regulatory activity by CRE-seq. P-values were calculated with two-tailed Student’s t-test. (D) CRISPR-Cas9 mutants. Sizes of deletions are indicated. Note that rs77910749 ‘knock-in’ introduces a 1 bp deletion. (E) Schematic of the ASE experiment (not to scale). Mice heterozygous for an LC1 mutation were mated to mice with a variant in the 3’ UTR of *Pou3f2*, which served as a molecular transcript barcode (light blue rectangle). Resulting ‘trans-het’ mice (heterozygous for both the LC1 mutation and the 3’ UTR variant) were analyzed for allele-specific *Pou3f2* expression at E14.5. The LC1 mutation is in *cis* to the wild-type 3’ UTR. To account for any effects of the 3’ UTR variant alone, control animals (wild-type for LC1 and heterozygous for the 3’ UTR variant) were included. (F) E14.5 whole brain, microdissected amygdala region, and microdissected anterior cortex were analyzed for allele-specific *Pou3f2* expression in control and trans-het LC1 KO animals (left panel), and in control and trans-het rs77910749 knock-in animals (right panel). Expression is normalized to controls. For trans-het LC1 KO whole brain, data were pooled across two lines with nearly identical deletions (see SI). P-values were calculated with two-tailed Student’s t-test. Error bars indicate SEM between biological replicates. Each biological replicate consisted of tissue from one embryo. Sample size per condition is indicated (amygdala and anterior cortex samples were collected from the same embryos). Non-significant, n.s.

We introduced this library into developing mouse cerebral cortex by *ex vivo* electroporation at E12.5, followed by two days of explant culture^58^. Histologic examination revealed GFP expression in the deeper cortical layers (Fig. 4B). By contrast, p*Dcx*-DsRed (a co-electroporated control construct) was expressed in the upper cortical layers, as expected^59^. *Dcx* encodes doublecortin, which is expressed in post-mitotic, migrating cortical neurons^60^. There was little colocalization of DsRed and GFP, suggesting that the CRE-seq library was not active in migrating neurons, but rather in progenitors and/or a subset of developing neurons in the cerebral cortex.

In parallel, we introduced the CRE-seq library into human iPSC-derived cerebral organoids (Fig. S10)^61,62^. Seven days after electroporation, live imaging showed electroporated cells expressing the p*CAG-*DsRed (a co-electroporated control construct with ubiquitous activity) (Fig. 4B). A subset of DsRed-expressing cells also expressed GFP, indicating CRE-seq library activity.

We then quantified the *cis*-regulatory activity of the constructs by barcode sequencing (Fig. 4C and SI). For both mouse cerebral cortex and human cerebral organoids, we observed enhancer activity of LC1 multimers (both ‘Ref’ and ‘Var’) relative to the promoter-only control. In the mouse cerebral cortex, the ‘Var’ multimer had ∼11% higher activity than ‘Ref’, while in the human cerebral organoids, the ‘Var’ multimer had ∼32% higher activity than ‘Ref’. Thus, rs77910749 confers significantly higher LC1 enhancer activity in two orthogonal assay systems.

### *In vivo* deletion of LC1 confers region-specific changes in *Pou3f2* expression

To directly address whether LC1 regulates *Pou3f2* expression and whether rs77910749 affects *Pou3f2* expression, we used CRISPR-Cas9 to delete the mouse LC1 region (∼1 kb) (‘LC1 KO’ mice). We also used CRISPR-Cas9 to knock-in rs77910749 into the orthologous position of the mouse genome (‘humanized’ KI mice) (Fig. 4D). A global survey of gene expression with E14.5 whole-brain RNA-seq of homozygous LC1 KO mice and homozygous rs77910749 KI mice (and corresponding controls) revealed minimal changes (Table S2). This suggested that LC1 may act in a cell type- and/or region-specific manner not detectable in whole-brain assays^63^.

As a more focused approach, we developed an allele-specific expression assay. First, we used CRISPR-Cas9 to generate mice with a small deletion (4 bp) in the 3’ UTR of *Pou3f2*, which serves as a barcode. Mice heterozygous for the LC1 deletion (‘LC1 het’) were crossed to mice with the 3’ UTR variant (Fig. 4E). By measuring allele-specific *Pou3f2* transcripts, we quantified changes in expression due to the LC1 KO allele relative to the LC1 wild-type allele.

Examination of the whole brain revealed no difference in allele-specific *Pou3f2* expression (Fig. 4F). Since the LacZ transgenic reporter assays suggested that LC1 is active in the amygdala and cerebral cortex (Fig. 3), we then analyzed the amygdala and cerebral cortex separately. No difference in *Pou3f2* expression was observed in the microdissected cortex. However, in the microdissected amygdala, the LC1 KO allele was associated with ∼8% higher *Pou3f2* expression (Fig. 4F). This suggests that LC1 acts as a silencer in a subset of cells in the amygdala at E14.5.

To test the effect of rs77910749 on *Pou3f2* expression, we crossed rs77910749 KI mice to *Pou3f2* 3’ UTR variant mice and conducted an analogous series of experiments. We did not observe any allele-specific changes in *Pou3f2* expression associated with rs77910749 in the whole brain, amygdala, or cerebral cortex at E14.5 (Fig. 4F). Altogether, these data suggest that LC1 has a role in regulating *Pou3f2*, but rs77910749 alone does not significantly alter *Pou3f2* expression at this level of tissue resolution in the developing mouse brain.

### LC1 knockout mice have normal behavior, but humanized rs77910749 knock-in mice have defective sensory gating

Next, we asked whether deletion of LC1 alters behavior. We subjected adult homozygous LC1 KO mice and wild-type siblings to a locomotion assay and sensorimotor battery, which did not detect any gross abnormalities. We then assayed the animals for the following: spatial learning and memory (Morris water maze), conditioned fear, sensorimotor reactivity and sensory gating (acoustic startle and prepulse inhibition), and anxiety (elevated plus maze and open field test) (File S1 and SI). The LC1 KO animals appeared normal as measured by these standard behavioral assays.

We then asked whether rs77910749 modifies mouse behavior. In homozygous rs77910749 KI mice compared to wild-type siblings, no abnormalities in locomotion, sensorimotor battery, Morris water maze, conditioned fear, or elevated plus maze were seen. However, when subjected to acoustic startle/prepulse inhibition (PPI) testing, the homozygous KI mice had a significant (p = 0.039, ANOVA) defect in PPI, with 22% less PPI compared to WT (Fig. 5; File S2 and SI). PPI is a measure of sensory gating and correlates strongly with altered cognition (thought disturbances and psychosis) in humans, and defective PPI is associated with BD, especially acute mania^64^. Furthermore, intact amygdala function is required for normal PPI^65^. Thus, rs77910749 KI mice have a specific defect in sensory gating, an amygdala-dependent BD endophenotype. No deficits in a social approach test or tail suspension test (measuring depressive behavior) were seen, further demonstrating the specificity of this behavioral deficit (File S2 and SI)^66,67^.

**Figure 5.**
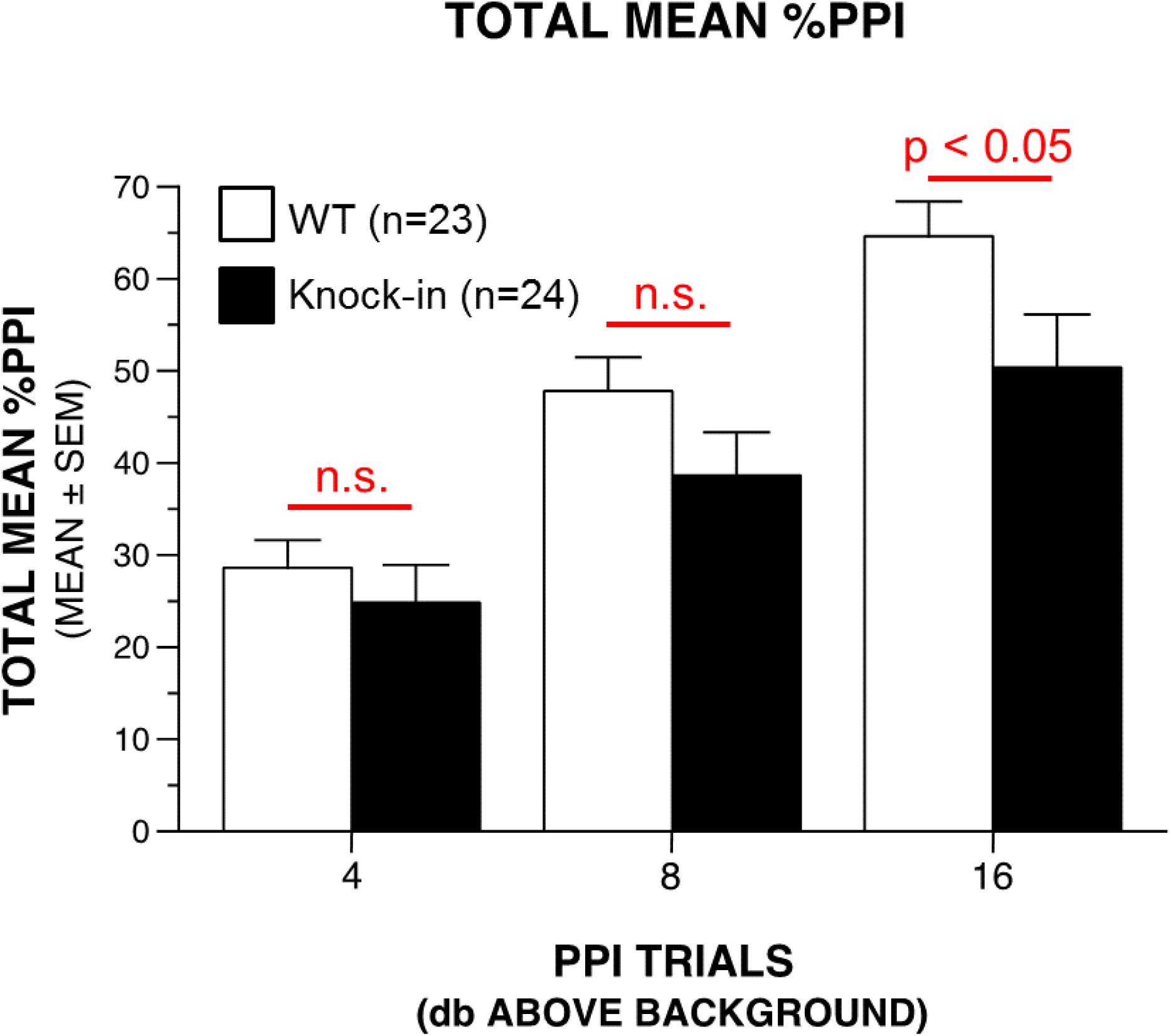
Prepulse inhibition (PPI) is defective in ‘humanized’ rs779710749 knock-in mice. Adult mice homozygous for the rs77910749 knock-in allele and wild-type (WT) siblings (age- and sex-matched) underwent acoustic startle testing with prepulse inhibition (PPI) assays. The knock-in (KI) animals showed defective prepulse inhibition that was statistically significant for the highest decibel (db) tested (p = 0.039, ANOVA). Mean %PPI ± SEM are shown (WT: 64.6 ± 3.8, KI: 50.4 ± 5.7). Single block %PPI analysis yielded similar results (File S2). One WT animal did not have a startle response at baseline and was excluded from the analysis. PPI measurements were normalized to baseline startle responses. Of note, baseline startle response magnitudes were lower in KI than WT animals (p = 0.018). Non-significant (n.s.) comparisons are indicated.

## DISCUSSION

Here, we sought to identify the ‘causal variant’ underlying GWAS signals at the *MIR2113/POU3F2* locus, which is associated with both enhanced cognitive performance and higher risk of BD. Our experiments suggest a molecular mechanism that links the human-specific non-coding variant rs77910749 to a BD-associated phenotype.

We found that rs77910749 falls within a PAX6 binding site and increases the binding affinity of PAX6. We then showed that rs77910749 falls within an active enhancer, LC1, and increases enhancer activity as assayed in developing mouse cerebral cortex and human cerebral organoids. We found that LC1 is active in the developing cerebral cortex, amygdala, and retina. CRISPR-Cas9 deletion of mouse LC1 altered *Pou3f2* expression in the amygdala. Remarkably, CRISPR-Cas9 knock-in of rs77910749 (‘humanized’ mice) resulted in defective sensory gating, an amygdala-dependent endophenotype seen in humans with BD.

While the amygdala had been strongly implicated in BD previously^5^, here we provide molecular evidence of a transcriptional program affecting the amygdala, with downstream effects on neuropsychiatric phenotypes. Future studies with greater spatiotemporal resolution may reveal the relevant neuronal subpopulations, while environmental or pharmacological perturbations of the humanized mice may elicit additional relevant phenotypes.

Here, we evaluated rs77910749 as a candidate causal variant. However, we cannot rule out the possibility that multiple tightly linked variants act together in a local ‘haplotype block’ for full phenotypic effect. The *MIR2113/POU3F2* intergenic region contains dozens of fetal brain-specific DHSs, which may act together or in a functionally redundant manner^68,69^. Additionally, *Pou3f2* and *Pou3f3* have largely overlapping expression patterns in the CNS and considerable functional redundancy in the mouse cerebral cortex^14,16^. These layers of redundancy reduce the likelihood that any single variant will profoundly alter neurodevelopment. Nonetheless, our study highlights a potential mechanism whereby a common variant gives rise to molecular and behavioral changes. Our studies underscore the notion that ostensibly positive traits, such as enhanced cognitive performance, may also confer susceptibility to neuropsychiatric disease.

## METHODS

### Reference genomes

Unless otherwise indicated, genomic coordinates are in hg19 (human) and mm9 (mouse).

### Custom materials and antibodies

Oligos, primers, and adapters are listed in Table S3. Buffers and media are listed in Table S4. Antibody information is provided in SI.

### Animals

Mice were kept on a 12 hour light/dark cycle at ∼20-22 °C with free access to food and water. Pregnant dams were euthanized with CO2 anesthesia and cervical dislocation. For timed pregnancies, mating occurred overnight and the next day was considered E0.5. All experiments were conducted in accordance with the Guide for the Care and Use of Laboratory Animals of the National Institutes of Health and approved by the Washington University Institutional Animal Care and Use Committee. Behavioral assays are described in SI.

### DNase-seq data

Human fetal DNase-seq data from Roadmap Epigenomics and mouse (C57BL/6) DNase-seq data from ENCODE were visualized in the UCSC Genome Browser (see SI)^31,39,70^.

### Calculation of linkage disequilibrium (LD)

Unless otherwise indicated, LD measures (r^2^ and D’) are based on EUR 1000G Phase 1, as calculated by HaploReg v4.1^32^.

### Electrophoretic mobility shift assays (EMSAs)

Quantitative EMSAs were conducted essentially as described (see also SI)^49^. Binding reactions were conducted light-protected at 4 °C for 1 hr. Protein-DNA complexes were separated on 10% TBE gels (Invitrogen), followed by imaging and quantification of band intensities.

### Generation of transgenic reporter mice

The LC1-*Hsp68*-LacZ construct was synthesized by cloning a 951 bp fragment of LC1 (chr6:98,566,099-98,567,049 in hg19, initially obtained by PCR of human gDNA) into the HindIII and PstI sites of *Hsp68-*LacZ Gateway vector^50^. The Sanger sequencing-confirmed construct was linearized with HindIII, gel-purified, and diluted with Microinjection Buffer. DNA was microinjected by the Washington University Mouse Genetics Core into fertilized eggs of C57BL/6 x CBA hybrid mice and implanted into pseudopregnant dams^71^.

### Mouse cerebral cortex electroporations

*Ex vivo* cerebral cortex electroporation of E12.5 CD-1 mouse embryos was conducted essentially as described (see also SI)^58^. After two days, electroporated regions were microdissected under a fluorescent microscope (Leica MZ16 F) in cold HBSS with calcium and magnesium and stored in TRIzol (Invitrogen) at -80 °C. Each biological replicate consisted of tissue from five to eight cortices.

### Human cerebral organoid electroporations

Cerebral organoids were cultured from human iPSCs (see SI). The CRE-seq library (1 μg/uL) was co-electroporated with p*CAG*-DsRed (1 μg/uL)^72^ into Day 88-109 organoids. After 7 days, organoids were rinsed with HBSS with calcium and magnesium and stored in TRIzol (Invitrogen) at -80 °C. Each biological replicate consisted of eight electroporated organoids.

### CRISPR-Cas9 mice generation

For CRISPR-Cas9 design, oligos, and genotyping, see Table S3 and SI. Microinjections were conducted in a C57BL/6J background by the Washington University Mouse Genetics Core and the Micro-injection Core (see SI).

### Allele-specific expression (ASE) analysis

E14.5 embryo brains were microdissected and processed with TRIzol (Invitrogen) for RNA and DNA extraction. The cDNA (from reverse transcription) and DNA underwent PCR to amplify the 3’UTR of *Pou3f2* for subsequent amplicon-seq (see SI). The allelic counts of variant and reference 3’ UTR sequences were tabulated to calculate normalized allele-specific expression.

### Data availability

RNA-seq data are available at Gene Expression Omnibus (GEO), accession GSE117877.

## ACKNOWLEDGEMENTS

We thank the Washington University Animal Behavior Core (David Wozniak), Center for Genome Sciences and Systems Biology (Jessica Hoisington-Lopez), Micro-injection Core (J. Michael White), Mouse Genetics Core (Mia Wallace), Protein and Nucleic Acid Chemistry Laboratory (Misty Veschak), Genome Engineering and iPSC Center (GEiC, Shondra Miller), and the Genome Technology Access Center for help with genomic analysis. GEiC is supported in part by NCI grant P30 CA091842, Eberlein, PI. We thank Qiang Lu for the p*Dcx*-DsRed construct and Matthew Toomey, Henry Lather, and Allison Loynd for assistance with experiments. We are grateful to Shuyi Ma, Daniel Murphy, Matthew Toomey, and Leo Volkov for critical reading of the manuscript. This work was supported by the NSF grant 1714867 to OG, NIH grant 5T32EY013360 to SQS, 5T32EY013360 to AH, and EY024958, EY025196, and EY026672 to JCC, and the Washington University Intellectual and Developmental Disability Research Center.

## AUTHOR CONTRIBUTIONS

SQS and JCC designed the experiments. SQS, JSK, LC, and CAM conducted experiments. SQS, AEH, DX, and OG conducted bioinformatic analyses. SQS and JCC wrote the manuscript.

