## Supplemental Information for "A candidate causal variant underlying both enhanced cognitive performance and increased risk of bipolar disorder"

Table of Contents

I.	Phylogenetic analysis of rs77910749 and rs17814604…………………………………….2

II.	Hi-C data……………………………………………………………………………….….2

III.	PD5a EMSA……………………………………………………………………………….2

IV.	Supplemental Methods…………………………………………………………………….3

V.	Supplemental Figure Legends…………………………………………………………......9

VI.	Supplemental Table Legends…………………………………………………………….12

VII.	Supplemental File Descriptions……………………………………………………..........13

VIII.	Supplemental References………………………………………………………………...14

**I. PHYLOGENETIC ANALYSIS OF RS77910749 AND RS17814604**

We noticed that the r^2^ value for rs17814604 (r^2^ = 0.43 with rs9320913) was relatively low despite a high D’ (0.99), suggesting a difference in allelic frequencies between rs17814604 and rs9320913. Indeed, rs17814604 was rarer than rs9320913, suggesting that individuals with rs17814604 are a subset of those with rs9320913.

Construction of a phylogenetic tree (Fig. S2) revealed that a ‘derived haplotype’ emerged containing rs77910749, rs13208578 in LC2, rs12204181 in LC4, and the lead SNPs rs10457441 (cognition), rs12202969 (BD), and rs9320913 (education). This derived haplotype (defined by rs77910749) is found worldwide but is not shared with Neanderthal and Denisovan genomes. As such, we surmised that the derived haplotype emerged after the Human-Neanderthal split, but before modern humans migrated out of Africa, placing the haplotype’s origin at least ~60,000 years ago^1^. The more recent haplotype (defined by rs17814604) is mostly confined to Europe and Southeast Asia, and hence likely evolved in the last 50,000 years (Fig. S3A).

Despite the high allele frequency of rs77910749, we found no evidence for a traditional selective sweep, suggesting that this variant does not dramatically alter fitness (Fig. S4). However, we cannot rule out more complex scenarios where selection acts in a time- or geography-dependent manner.

Lastly, we examined vertebrate genomes, including all 79 non-human primate individuals (representing five species) sequenced in the Great Ape Genome Project^2^ (Fig. S5). This analysis demonstrated that rs77910749 appears to be a human-specific variant, absent from other known primate and vertebrate genomes.

**II. HI-C DATA**

A survey of published Hi-C data revealed that LC1 falls within a topologically-associated domain (TAD) in multiple mouse and human cell types, suggesting that this is an evolutionarily conserved and cell-type invariant TAD^3-7^ (Fig. S6).

**III. PD5A EMSA**

PD5a is a brain-expressed splice isoform of PAX6 that contains a 14-amino acid insertion in the PAI domain of PD. PD5a has a known distinct DNA binding preference compared to canonical PAX6^8,9^. Neither PD5a nor PD5a-HD bound to either the reference or variant sequence under conditions in which PD and PD-HD bound the probes robustly (Fig. S9).

**IV. SUPPLEMENTAL METHODS**

**DNase-seq data sources**

The following human fetal DNase-seq data from Roadmap Epigenomics were visualized in the UCSC Genome Browser^10,11^. Donor name, age, sex, and GEO accession are listed: fBrain #1 (donor H-23284, 96 day female, GSM595928), fBrain #2 (donor H-22911, 117 day female, GSM595920), fBrain #3 (donor H-22510, 122 day male, GSM530651), fHeart (donor H-23604, 110 day female, GSM665830), fKidney (donor H-22676, 122 day sex unknown, GSM530655), fLung (donor H-22727, 101 day sex unknown, GSM530662), and fThymus (donor H-23964, 98 day female, GSM701537). The following mouse (C57BL/6) DNase-seq data from ENCODE were visualized^12^. Mice were 8 weeks old unless otherwise indicated: E14.5 brain (GSM1014197), E18.5 brain (GSM1014184), adult brain (GSM1014151), P1 retina (GSM1014188), P7 retina (GSM1014198), adult retina (GSM1014175), adult heart (GSM1014166), adult kidney (GSM1014193), adult lung (GSM1014194), E14.5 liver (GSM1014183), adult liver (GSM1014195), and adult thymus (GSM1014185).

**Construction of phylogenetic tree, haplotype analysis, and measures of a classical sweep**

We used phased haplotypes from 1000 Genomes Project Phase 3^13^ to construct a phylogenetic tree using VCFtoTree^14^ and visualized it in Dendroscope^15^. To check for signatures for any sweeps in the region, we used the 1000 Genomes Selection Browser^16^.

**Analysis of primate genomes**

Variant calls (SNPs and indels) for primate genomes^2^ were downloaded as VCF files in hg18 from <https://eichlerlab.gs.washington.edu/greatape/data/VCFs/>. VCFtools v0.1.10^2^ was used to obtain variants in the interval Chr6:98,673,000-98,674,000 in hg18, which is equivalent to Chr6:98,566,279-98,567,279 in hg19. Variants in this 1 kb window were manually examined for rs77910749, which is at chr6:98,673,228 in hg18 or chr6:98,566,507 in hg 19. The LiftOver tool on the UCSC Genome Browser was used to convert between hg18 and hg19^11^.

**Allele-specific methylation analysis**

Sample preparation and analysis were conducted essentially as previously described^17^. DNA was extracted from brain tissue with the DNeasy kit (Qiagen). For each biological replicate, ‘whole brain’ was dissected as described below for ASE, and the right half of the brain was used for bisulfite analysis. About 1 μg DNA was bisulfite-converted with EpiTect Bisulfite Kit (Qiagen) and subjected to PCR with LC1_bis_F and LC1_bis_R primers (Table S3). The resulting products were cloned into the pCR2.1 TOPO vector (Invitrogen) and Sanger sequenced with universal M13 reverse primer. Sequence data were analyzed and visualized with BISMA using default parameters with removal of PCR duplicates^18^.

**Motif analysis**

For ‘SELEX PWM’ scores (Fig. 2B), FIMO in MEME v4.9.1^19,20^ was used with the default p-value threshold (0.0001) to scan for occurrences of TF motifs using input TF motifs from^21^. The PAX6 motif was the only one identified that overlapped with rs77910749. The logo for the SELEX motif was generated in enoLOGOS^22^ using default parameters with *M. musculus* %GC. The logo for the ChIP-seq motif is based on^23^.

**Pax6 protein expression and EMSA conditions**

PAX6 is perfectly conserved at the amino acid level between mouse and human^24^. PD, PD-HD, PD5a, and PD5a-HD sequences were ordered as gene blocks from Integrated DNA Technologies with *E. coli* codon optimization and cloned as NdeI/NotI fragments into the pET-28a(+) vector. Constructs were confirmed by Sanger sequencing.

BL21 cells were transformed and induced with IPTG overnight at 16 °C. His-tagged proteins were purified with HisPur Ni-NTA Resin (Thermo Scientific), dialyzed with phosphate buffered saline (PBS), and concentrated with Amicon Ultra-4 10K MWCO (Millipore). Proteins were quantified with the Pierce BCA Assay Kit (Thermo Scientific) and stored in 50% glycerol at -80 °C.

EMSA binding reactions were conducted light-protected at 4 °C for 1 hr in EMSA Binding Buffer 1. For PD-HD and PD5a-HD reactions in Fig. S9, the binding reactions were conducted in EMSA Binding Buffer 2. Concentration of labeled probes was ~10 nM (for cold competition, 500-fold molar excess of unlabeled probe was used). Protein concentration was ~1 μM.

Protein-DNA complexes were separated on 10% TBE gels (Invitrogen) at 100 V for 90 min, light-protected at room temperature. Gels were imaged with variable-mode Typhoon scanners with excitation laser at 532 nm, emission filter at 526 nm for FAM, and emission filter at 610 nm for ROX. Band intensities were quantified with ImageQuant TL 8 (GE Healthcare Life Sciences).

**LacZ staining and histology**

E14.5 embryos were dissected in cold PBS. The tail (plus yolk sac for transient transgenics) was saved for PCR genotyping with LacZ primers. Embryos were rinsed with PBS with 0.1% Tween-20 and then fixed on ice for 90 min LacZ Fixing Buffer. After rinsing three times with LacZ Wash Buffer, embryos were incubated with X-gal Staining Buffer. Incubation was conducted at 37 ⁰C overnight (up to several days, with fresh X-gal Staining Buffer added every ~12 hr). Embryos were post-fixed with 4% paraformaldehyde in PBS and stored at 4 ⁰C until whole-mount imaging. For cryosections, embryos were equilibrated in 30% sucrose/PBS and decapitated. The head was embedded in Tissue-Tek OCT (Sakura) and cryosectioned at 20 μm. Sections were rinsed with PBS and counterstained with Nuclear Fast Red (Sigma).

**CRE-seq library construction**

To create the LC1 multimer constructs, individual 200 bp sequences (centered on the position of rs77910749) were obtained by PCR using template DNA with or without rs77910749, with primers to add restriction enzyme sites. These ‘monomers’ were ligated pairwise in two rounds to create the 4X multimer with or without the variant (NotI-LC1-XbaI-LC1-XhoI-LC1-XmaI-LC1-FseI). The LC1 monomer with rs77910749 includes an additional ‘T’ base at the 3’ end, such that the length and base content is the same as the LC1 monomer without rs77910749. Multimers were confirmed by Sanger sequencing. Each multimer was then cloned into the NotI/FseI sites of the previously described CRE-seq vector, which has random 15 bp barcodes in the 3’ UTR^25^. The 3.6 kb *POU3F2* promoter encompassing chr6:99,279,024-99,282,671 (hg19) was obtained by PCR of human gDNA. The basal *rho* promoter-GFP cassette of the CRE-seq vector was replaced with a 3.6 kb *POU3F2* promoter-GFP cassette between the FseI and AscI sites. The LC1 multimer was cloned into the NotI/FseI site, and individual colonies were picked for Sanger sequencing with Barcode_seq_R to determine barcode sequences. For the promoter-only control constructs, there was no insert upstream of the 3.6 kb *POU3F2* promoter. Twenty barcoded constructs were obtained for each of the LC1 REF multimer, the LC1 VAR multimer, and the promoter-only control. Each pool of twenty constructs was maxiprepped (Invitrogen) and then pooled together for electroporations.

**Electroporation conditions for CRE-seq in *ex vivo* mouse cerebral cortex**

The CRE-seq library (2.5 μg/uL) was pooled with p*Dcx*-DsRed (1 μg/uL) for a total of 3.5 μg/uL DNA. To visualize the injection, Fast Green dye (~0.02%) was added. DNA was injected with a pulled glass pipette and Hamilton syringe. Electroporation was conducted with BTX ECM830 (Harvard Apparatus): 33 V, 50 ms pulse duration separated by 950 ms intervals, for 5 pulses. After electroporation, the head was transected just superior to the level of the eye and transferred on ice to Explant Media. Up to three heads were arranged on a 25 mm circular Whatman Nucleopore 0.2 μm filter (cut surface against the shiny side of the filter), which floated in one well of a 6-well dish containing Explant Media with 1X B27 (Gibco) and 1X G5 (Gibco). Explants were incubated at 37 °C with 5% CO_2_.

**Culture and electroporation conditions for CRE-seq in cerebral organoids**

Human iPS(IMR90)-4 (WiCell) were cultured in mTeSR1 (STEMCELL Technologies) and passaged every 3-4 days at 1:10 with ReleSR (STEMCELL Technologies) on 6-well plates with Matrigel (Corning)^26,27^. Cells were maintained at 37 °C with 5% CO_2_. Cells (passage 64) were differentiated following a protocol similar to that of^28^. On the day of passage (Day 0), cell aggregates were released with ReLeSRTM and floated freely in a 100 mm low-bind Petri dish in Neural Induction Media. On Day 18, media was changed to Cerebral Growth Media similar to that of^29^. Media was changed every 3-4 days.

For electroporation, the same equipment and similar protocol as for *ex vivo* retinal electroporations were used (see SI)^30^. Four organoids were loaded into an electroporation chamber and allowed to float freely. Electroporation settings were: 35 V, 50 ms pulse duration separated by 950 ms intervals, for five pulses. Organoids were placed back into conditioned Cerebral Growth Media and allowed to float freely.

**Histology, imaging, and antibodies**

For Fig. 4B, two days after electroporation, mouse cerebral cortex tissue was fixed in 4% paraformaldehyde/PBS, embedded in 4% agarose, and vibratome sectioned at 100-200 μm. Sections were mounted with Vectashield (Vectorlabs), coverslipped, and subjected to laser confocal imaging (Zeiss LSM700) with ZEN 2009 software (Zeiss).

Prior to harvest, human iPSC-derived cerebral organoids were live-imaged (Fig. 4B) with an inverted fluorescent microscope (Nikon Eclipse TE300). For Fig. S10, organoids were fixed in 4% paraformaldehyde/PBS for 45 min, equilibrated in 30% sucrose/PBS and embedded in Tissue-Tek OCT (Sakura) for cryosections (12-14 μm). Confocal images were collected on a BX61 WI microscope (Olympus) with a DSU spinning disk and ORCA-ER CCD camera (Hamamatsu) and processed with MetaMorph (Molecular Devices).

The following antibodies were used: anti-PAX6 (PRB-278P at 1:300), anti-POU3F2 (sc-6029 at 1:80), and anti-KI67 (BD Pharmigen 550609 at 1:100). Note that the anti-POU3F2 antibody recognizes both POU3F2 and POU3F3^31^.

**CRE-seq tissue processing and data analysis**

RNA and DNA were isolated with TRIzol (Invitrogen), treated with TURBO DNase (Ambion), and purified with RNeasy Mini (Qiagen) as described^25^. RNA (~0.5-1 μg) was reverse-transcribed with SuperScript IV (Invitrogen), and the cDNA was treated with RNaseH. The barcode region of the cDNA and DNA was amplified by PCR with Nano_initial_PCR primers using Phusion (New England BioLabs) as follows: 98 °C for 30 sec, X cycles of 98 °C for 10 sec, 64 °C for 30 sec, 72 °C for 30 sec, and finally 72 °C for 5 min. The number of PCR cycles (X) was: 16 for DNA, 20 for mouse cortex cDNA, and 22 for organoid cDNA. Samples were then prepared for amplicon-seq (see below).

Barcode sequences were bioinformatically extracted as a perfect match to one of the sixty known 15 bp barcodes plus 6 bp of flanking sequence on either side (27 total bp) in either the forward or reverse direction (since sequencing adapters were ligated non-directionally). The RNA read count was normalized to the DNA read count for each barcode and then averaged across the twenty barcodes to yield the overall activity of a construct type (‘Ref’, ‘Var’, or promoter-only) in a given biological replicate.

**CRISPR-Cas9 mouse generation**

CRISPR/Cas9 reagents^32,33^ were generated at the Genome Engineering and iPSC Center at Washington University. For the ‘LC1 KO’ allele, a pair of guides flanking LC1 was used to delete the intervening sequence. Multiple lines were generated with nearly identical deletions, but Line 2674 (deletion of chr4:23,438,846-23,439,893, depicted in Fig. 4D) was used for all ‘LC1 KO’ experiments, with the exception of Fig. 4F LC1 knock-out ‘whole brain,’ which has pooled data from Line 2674 and Line 2677 (deletion of chr4:23,438,848-23,439,894). For knock-in of rs77910749, a single-stranded donor oligo centered on the variant (with ~60 bp of homology on either side) was injected with a central LC1 guide for homologous recombination^34^. For the ‘*Pou3f2* 3’UTR variant’, a single guide in the 3’ UTR was used to generate a 4 bp deletion (chr4:22,412,587-22,412,590). For the ‘LC1 Small Indel’ (Fig. S8), the central LC1 guide was used alongside a central LC5 guide, such that this strain carries a 14 bp deletion (chr4:23,439,327-23,439,340) plus an insertion of ‘C’ at the site of the deletion, as well as a 103 bp deletion in LC5 (chr4:23,417,446-23,417,548).

Hormone-primed females were mated to generate embryos (E0.5), which underwent pronuclear micro-injection of 2.5 ng/μl guide RNA and 5 ng/μl of Cas9 mRNA. Embryos were transferred to pseudo-pregnant recipient females. Founders (F0’s) were outbred to C57BL/6J. For genotyping, alleles were confirmed by Sanger sequencing of PCR products. After the F1 generation, LC1 KO animals were genotyped by PCR only.

**RNA-seq**

E14.5 embryos were harvested in cold HBSS with calcium and magnesium, and whole brains were rapidly dissected and stored in TRIzol (Invitrogen) at -80 °C. Each brain represented a single biological replicate. Tail tissue was saved from each embryo for genotyping. Sex-matched, litter-matched wild-type and mutant embryos were used. RNA was extracted with TRIzol, treated with DNase I, and purified with RNeasy Mini (Qiagen). Agilent 2100 Bioanalyzer was used to verify RNA integrity. Library preparation for mRNA-seq was essentially as previously described ^35^. The Illumina oligo-dT kit was used according to the manufacturer’s protocol. Twelve samples (three biological replicates each of LC1 knockout homozygotes vs. corresponding wild-type controls, and rs77910749 knock-in homozygotes vs. corresponding wild-type controls) were indexed and pooled for sequencing on a lane of HiSeq 3000 with single 50 bp reads.

Reads were aligned to the Ensembl release 76 assembly with STAR v2.0.4b^36^, gene counts were obtained with Subread:featureCount v1.4.5, and transcript counts were generated with Sailfish v0.6.3^37^. Sequencing performance was assessed with RSeQC v2.3^38^. Counts were imported and TMM normalization size factors were calculated in EdgeR^39^. Limma and weighted likelihoods based on the observed mean-variance relationship of every gene/transcript were generated with Voom^40^. Generalized linear models were used to test for differential expression.

**Allele-specific expression analysis**

E14.5 embryos were harvested in cold HBSS with calcium and magnesium, and brain tissue was rapidly dissected and stored in TRIzol (Invitrogen) at -80 °C. For ‘whole brain’ dissection, the olfactory lobes were left intact, and the brain was transected coronally at the posterior edge of the cortex (i.e., through the midbrain). For ‘amygdala region’ and ‘anterior cortex’ microdissection, the brain was also transected coronally at the anterior-posterior level of the middle cerebral artery in the circle of Willis. The anterior tissue (with olfactory lobes removed) was harvested as ‘anterior cortex’. The inferior and lateral portions of the posterior tissue were harvested as ‘amygdala region’. For rs77910749 knock-in ‘whole brain’, only the left half was used for ASE (the right half was used for methylation studies). Tail tissue was saved for genotyping.

RNA was extracted with TRIzol (Invitrogen), treated with TURBO DNase (Invitrogen), and purified with RNeasy Mini (Qiagen). RNA (~1-2 μg) was then reverse-transcribed with SuperScript IV (Invitrogen) and treated with RNase. The 3’ UTR of *Pou3f2* was amplified with Pou3f2_3UTR_F and Pou3f2_3UTR_R primers using Phusion (New England BioLabs) as follows: 98 °C for 30 sec, X cycles of 98 °C for 10 sec, 64 °C for 30 sec, 72 °C for 30 sec, and finally 72 °C for 5 min. The number of PCR cycles (X) was: 20 for whole brain, 21 for half brain, and 22 for microdissected regions.

After amplicon-seq, reads containing ‘CGTATATATATGGG’ (wild-type 3’ UTR) or ‘TGCGTATATGGGAT’ (variant 3’ UTR) were tabulated, and the ratio of reads (i.e., allelic bias) was calculated. The 3’ UTR variant itself causes ~10% increased *Pou3f2* mRNA levels compared to the wild-type 3’ UTR, as seen in the controls (which were WT for LC1 but heterozygous for the 3’ UTR variant). In Fig. 4F, the ratio of reads in the LC1 trans-het animals were normalized to the ratio of reads in the controls.

**Amplicon-seq**

Qubit dsDNA HS Assay (Invitrogen) was used to quantify samples. About 200 ng of cDNA or DNA was end repaired, 3’ adenylated, and ligated to MiSeq adapters according to standard protocols ^41^. The product was then amplified with a universal Illumina PCR primer and an indexed primer with Phusion as follows: 98 °C for 30 sec, 18 cycles (for ASE) or 20 (for CRE-seq) cycles of 98 °C for 10 sec, 57 °C for 30 sec, 72 °C for 30 sec, and finally 72 °C for 5 min. Products were gel-purified and verified on an Agilent Bioanalyzer. For a given sequencing run, up to six indexed samples were pooled. Samples were loaded at 7-8 pM onto MiSeq for 2x250 bp sequencing, representing ~10% of reads on a full lane, yielding ~1-2 million reads per pool of samples. Reads were demultiplexed and checked with FastQC^42^.

**Behavioral assays**

Animals were habituated in the Washington University Animal Behavior Core facility for at least 1 week before testing^43^. A total of 10 homozygous LC1 knockout and 10 age-matched, sex-matched wild-type siblings were subjected to the following tests at age 10-15 weeks: 1-hour locomotor activity, sensorimotor battery, Morris water maze, conditioned fear, acoustic startle and PPI, elevated plus maze, and open field test.

A total of 12 homozygous rs77910749 knock-in and 12 age-matched, sex-matched wild-type siblings were subjected to the same tests at age 10-15 weeks. For the acoustic startle and PPI assays in the rs77910749 knock-in animals vs. wild-type animals, two independent cohorts were tested and data from these two cohorts were pooled for a total of 24 homozygous rs77910749 knock-in and 24 wild-type control animals. For tail suspension and social approach tests, a total of 10 homozygous rs77910749 knock-in and 10 age-matched, sex-matched wild-type siblings were tested at age 12-18 weeks. All behavioral data are provided in Files S1 and File S2.

**V. SUPPLEMENTAL FIGURE LEGENDS**

**Figure S1. Phylogenetic conservation of rs13208578 and rs77910749.** Multiz alignments of vertebrates representing seven phylogenetic groups from the UCSC Genome Browser in hg19 ^11,44^. A 100 bp window is shown centered on each variant (position of variant is highlighted in red): (A) rs13208578 (a substitution of ‘C’ to ‘T’) with non-reference bases in red font, and (B) rs77910749 (a 1 bp deletion of a ‘T’).

**Figure S2. Identification of a derived haplotype through construction of a human phylogenetic tree.** (A) Phylogenetic tree of phased human haplotypes. Note that rs77910749 is one of the indicator variants for the derived haplotype (pink circle), and that rs17814604 emerged from this haplotype (green circle). (B) Haplotype analysis. The derived haplotype contains GWAS variants (red font) and linked variants (blue font) that fall within LC1 to LC5. The analysis was anchored on rs10457441. See also Figure 1.

**Figure S3. Global distribution of rs17814604 and rs77910749 frequencies.** Allele frequencies based on Phase 3 of the 1000 Genomes Project^13^ are shown for the major populations (large pie charts) as well as for subpopulations (small pie charts), with black indicating the reference allele: (A) rs17814604 (green allele) and (B) rs77910749 (purple allele). Abbreviations: AFR, African; AMR, American; BEB, Bengali in Bangladesh; CDX, Chinese Dai in Xishuangbanna, China; CHB, Han Chinese in Beijing, China; CHS, Southern Han Chinese, China; CLM, Colombian in Medellin, Colombia; EAS, East Asian; ESN, Esan in Nigeria; EUR, European; FIN, Finnish in Finland; GBR, British in England and Scotland; IBS, Iberian populations in Spain; JPT, Japanese in Tokyo, Japan; KHV, Kinh in Ho Chi Minh City, Vietnam; LWK, Luhya in Webuye, Kenya; MAG, Mandinka in The Gambia; MSL, Mende in Sierra Leone; PEL, Peruvian in Lima, Peru; PJL, Punjabi in Lahore, Pakistan; PUR, Puerto Rican in Puerto Rico; SAS, South Asian; TSI, Toscani in Italy; YRI, Yoruba in Ibadan, Nigeria.

**Figure S4. No evidence of a selective sweep in the genomic region around rs77910749.** The 1000 Genomes Selection Browser was used to search for signatures of selection^16^. (A) Comparison of Pi and Tajima's D values for the: i) target region of interest (chr6:98,565,000-98,585,000) defined by the 20 kb region encompassing rs77910749; ii) flanking region of 2 Mb around rs77910749 (chr6:97,500,000-99,500,000±1,000 kb); and iii) neutral regions consisting of randomly selected 25-35 kb putatively neutral regions across the genome (from the Neutral Region Explorer^45^). Pi is a measure of haplotype diversity. Even though it is highly sensitive to mutation rate, lower Pi values are expected in regions under negative selection. Tajima’s D measures deviation from the expected allele frequency spectrum^46^. Negative values indicate an excess of rare alleles and are consistent with negative selection or complete selective sweeps. Positive values (an abundance of intermediate frequency alleles) are consistent with balancing selection. A Tajima's D near zero is consistent with no evidence of selection. Here, both the target and flanking regions show lower values for Pi and Tajima's D compared to the corresponding values for neutral regions. The most parsimonious explanation is that this locus is under negative selection. (B) Comparison of the Fst values in the target region (harboring rs77910749) vs. the flanking region did not reveal a clear difference. Fst is a measure of among-population variation as compared to within-population variation. Higher values indicate higher levels of population differentiation, which is often regarded as a signature of population-specific adaptive forces^47^. GLO, global Fst (expected heterozygosity). CEU, Utah residents with Northern and Western European Ancestry. CHB, Han Chinese in Beijing, China. YRI, Yoruba in Ibadan, Nigeria. (C) Comparison of Fst values for different allele frequency clusters (in target vs. flanking regions) did not identify clear outliers.

**Figure S5. Absence of rs77910749 from non-human primate genomes.** The genomes of 79 individuals from five non-human primate species (Sumatran orangutan, Bornean orangutan, gorilla, chimp, and bonobo) were examined for rs77910749, and none were found to contain this variant. Number of individuals for each species is indicated. Sequences and estimates of divergence times (in millions of years ago, MYA) are from Prado-Martinez et al., 2013^2^. Minor allele frequency (MAF) in humans is based on aggregate 1000 Genomes Phase 3 data^13^.

**Figure S6. LC1 falls within a conserved topologically associating domain (TAD).** Published Hi-C data were visualized with default heat map scaling on the 3D Genome Browser (<http://www.3dgenome.org>). Darker red indicates higher frequency of interactions. Data are shown at 40 kb resolution, except for CH12 (25 kb resolution). The TAD containing LC1 is highlighted in yellow, and LC1 (labeled) is highlighted in pink. Note the positions of *Mir2113/MIR2113* and *Pou3f2/POU3F2* (red font) within the TADs. In the mouse genome, the region is inverted such that the relative orientation (LC1 upstream of *Pou3f2*) is preserved. Mouse *Mir2113* is a non-RefSeq gene identified by homology with the human sequence. (A) Human Hi-C data for left ventricle of heart, liver, and H1-derived neural progenitor cells (NPC)^3,6^. (B) Mouse Hi-C data for CH12 (a B cell lymphoma cell line), embryonic stem cells (ESC), and cerebral cortex^4,5^.

**Figure S7. The methylation landscape of LC1 to LC5 in human primary tissues and cultured cells.** Active enhancers are typically unmethylated^48^. LC1 is highlighted (pink), and LC2 to LC5 are also shown (gray). LC1 is unmethylated in fetal brain and neural progenitors, and methylated in adult brain and other tissues. MRE enriches for unmethylated regions, while MeDIP enriches for methylated regions^49^. The height of MethylC-seq and bisulfite (BS)-seq signals reflects the degree of methylation at single CpG resolution. The lack of data for LC3 and LC5 is due to the paucity of CpG sites. Data are from Roadmap Epigenomics^10^ except for MethylC-seq^50,51^, with the following GEO accessions: MRE (GSM669604 and GSM707015) and the corresponding MeDIP (GSM669614 and GSM707019) of fetal brain (fBrain), MethylC-seq (GSE47966) of fetal frontal cortex (fFC) and middle frontal gyrus (MFG, part of the cortex) at the indicated ages (d = day, yo = years old), H1-derived neuronal progenitor cells (HDNP) (GSM675546), brain germinal matrix (BGM) (GSM941747), NCD neurosphere culture (cortex-derived) (GSM1127118), left ventricle (LV) of heart (GSM1010978), lung (GSM983647), liver (GSM916049), fetal thymus (GSM1172595), thymus (GSM1010979), and fetal muscle (fMuscle) from leg (GSM1172596).

**Figure S8. Allele-specific methylation analysis of mouse LC1.** (A) Region within LC1 analyzed by bisulfite sequencing. ‘Knock-in’ of rs77910749 introduces a single bp deletion of ‘T’ (on the minus strand for mm9), creating a novel CpG site (site #6). (B) Bisulfite sequencing of E14.5 brain from mice that were heterozygous for rs77910749 knock-in (KI) allele (n = 4, left panels), or the LC1 Small Indel allele (n = 3, right panels). Each row represents a clone, and each column represents a CpG site. Note the two clones (pink arrows) of the KI allele, in which site #6 is methylated. The LC1 Small Indel allele also has a 103 bp deletion within LC5 (see Methods). Red = methylated, blue = unmethylated, white = no data. Note that CpG site #6 is absent in the WT allele and LC1 Small Indel allele. (C) Quantification of methylation at each CpG site. Top: rs77910749 knock-in heterozygotes. Bottom: LC1 Small Indel heterozygotes. No data (N.D.) for site #6 in the WT allele or indel allele given the absence of site #6. Error bars indicate SEM. P-values were calculated with two-tailed Fisher’s exact test (combining reads across replicates).

**Figure S9. EMSA analysis of PAX6 PD5a binding.** The PD5a splice isoform of PAX6 has a 14 amino acid insertion (yellow segment) within PAI. Under conditions in which PAX6 PD (left panels, lanes 1-3) and PD-HD (right panels, lanes 1-3) robustly bound to both reference (top channel) and variant (bottom channel) probes, PD5a (left panels, lanes 6-8) and PD5a-HD (right panels, lanes 6-8) failed to bind, reflecting differences in DNA binding preferences. Cold competition reactions: lanes 4 and 9. Probe and marker dye only (no protein): lanes 5 and 10. PD, paired domain. HD, homeodomain.

**Figure S10. Antibody staining of cerebral organoids.** Human iPSC-derived cerebral organoids expressing PAX6 and POU3F2/POU3F3. (The anti-POU3F antibody recognizes both POU3F2 and POU3F3^31^.) Human iPSCs were differentiated into cerebral organoids and grown in culture for 53 days (left panels) or 70-79 days (right panels) prior to harvest for immunohistochemistry. Cryosections were labeled with anti-POU3F antibody (green, all panels) and anti-PAX6 antibody (red, top panels) or anti-KI67 antibody (red, bottom panels). KI67 is a marker of proliferation^52^. Blue, DAPI counterstain.

**VI. SUPPLEMENTAL TABLE LEGENDS**

**Table S1. Measures of LD among lead SNPs in GWAS’s of educational attainment, cognitive ability, and BD.** Pairwise r^2^ and D’ values among the following five lead GWAS are shown: rs9320913 for educational attainment^53,54^, rs1906252 for cognitive ability^55,56^, rs10457441 for cognitive ability^57^, rs12202969 for BD^58^, and rs1487441 for BD^59^. Values were retrieved from HaploReg V4.1^60^ for European populations based on 1000 Genomes Phase 1^61^.

**Table S2. RNA-seq differential expression analysis.** Whole brains were harvested at E14.5, and the experiments were conducted in triplicate with each replicate consisting of a single brain. Comparisons were made between LC1 knockout vs. sex-matched wild-type littermate (first sheet), and between rs77910749 knock-in (RKI) homozygotes vs. sex-matched wild-type littermates (second sheet). For either comparison, no genes were significantly differentially expressed at an FDR-adjusted p-value of less than 0.05. Raw and processed data are available in Gene Expression Omnibus (GEO) under accession GSE117877.

**Table S3. Oligonucleotides used in this study.**

**Table S4. Buffers and media used in this study.**

**VII. SUPPLEMENTAL FILE DESCRIPTIONS**

**File S1. Additional behavioral data for homozygous LC1 knockout mice and wild-type siblings.** Animals were age- and sex-matched. Data are from the following assays: 1-hour locomotor activity, sensorimotor battery, Morris water maze, conditioned fear, acoustic startle/PPI, elevated plus maze, and open-field test. Sample sizes and p-values (ANOVA) are indicated. Wild-type (WT), white circles and bars. Mutant (MT), black squares and bars.

**File S2. Additional behavioral data for homozygous rs77910749 knock-in mice and wild-type siblings.** Animals were age- and sex-matched. Data are from the following assays: 1-hour locomotor activity, sensorimotor battery, Morris water maze, conditioned fear, acoustic startle/PPI, elevated plus maze, social approach test, and tail suspension test. Sample sizes and p-values (ANOVA) are indicated. Wild-type (WT), white circles and bars. Mutant (MT), black squares and bars.
