## Supplemental Figures for "A candidate causal variant underlying both enhanced cognitive performance and increased risk of bipolar disorder"

Supplemental Figure S1

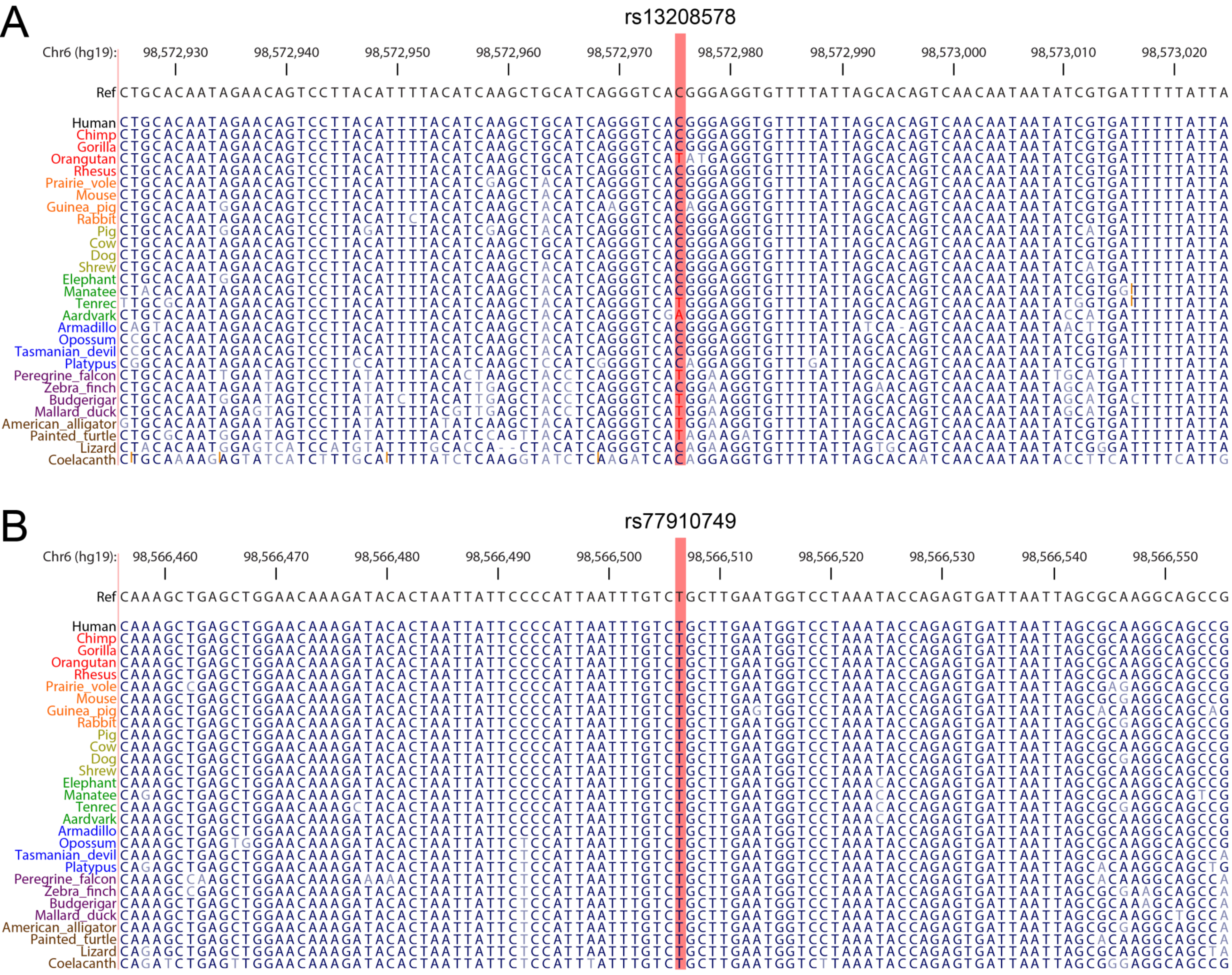

Supplemental Figure S2

A

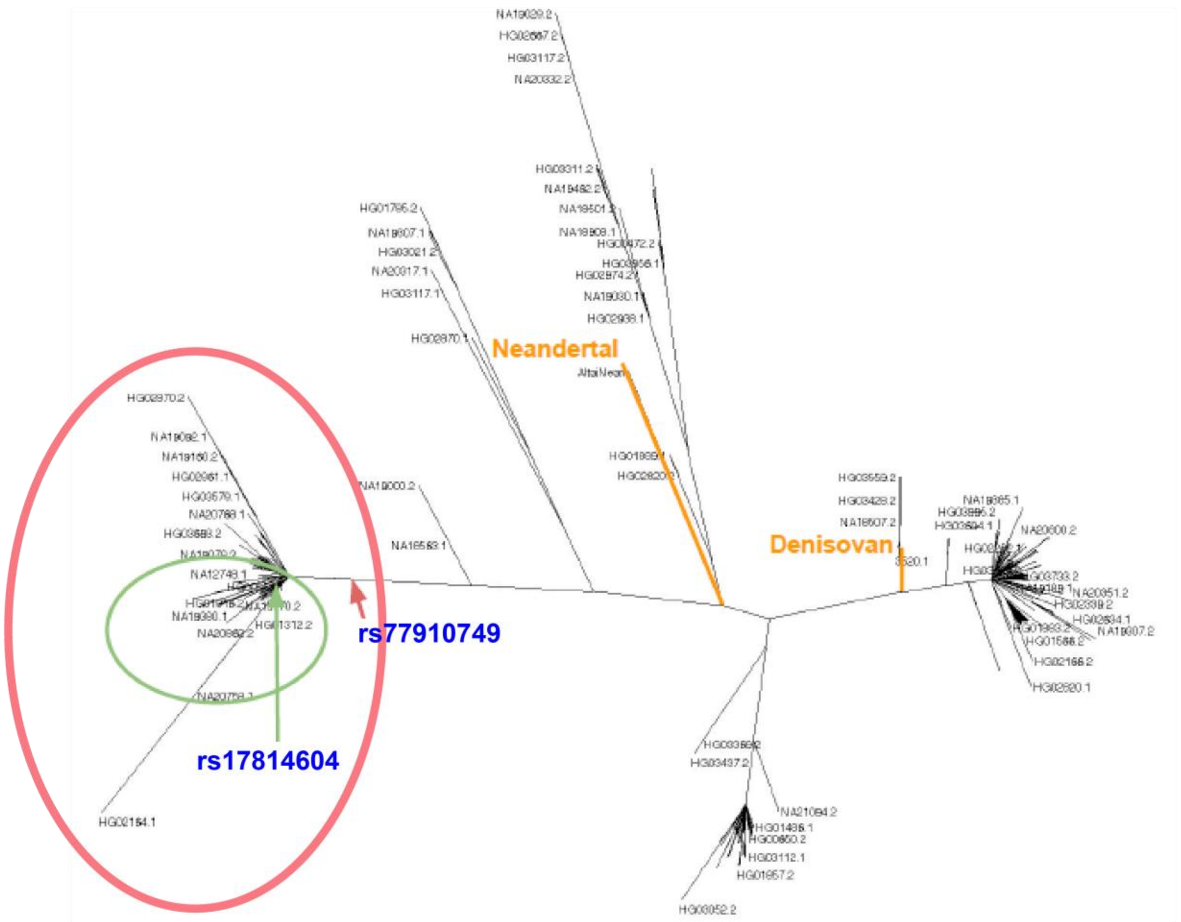


B

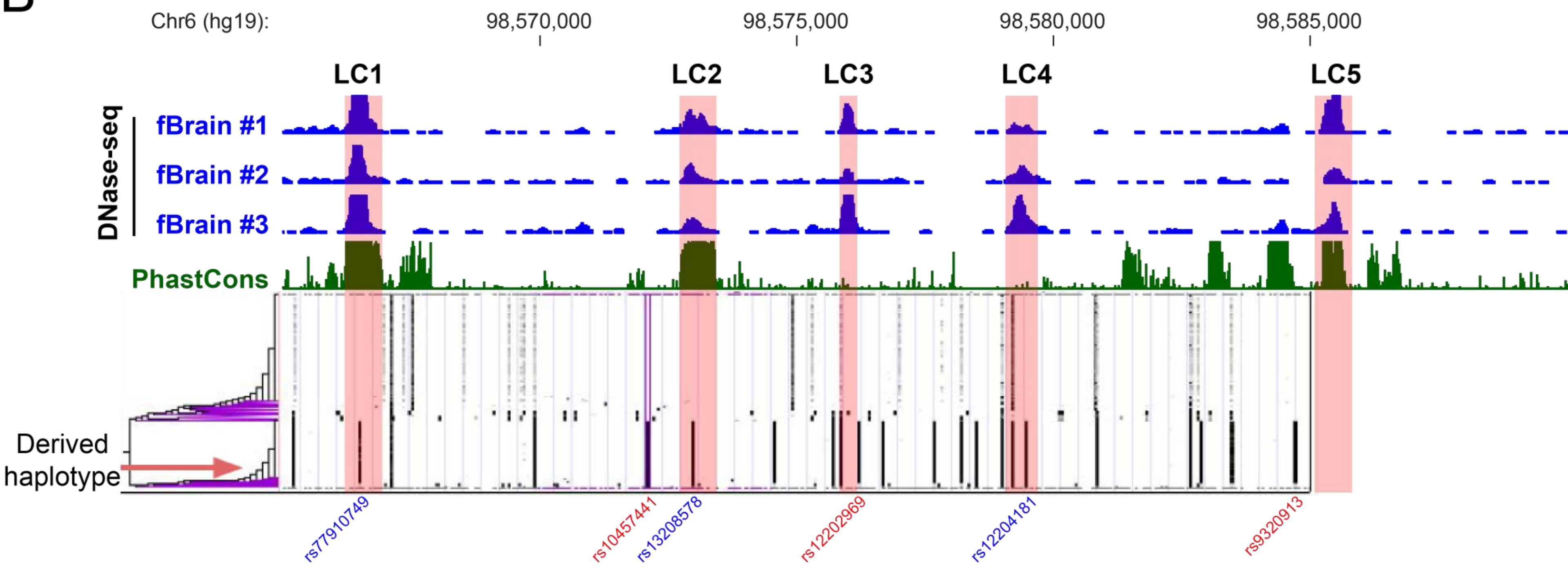

Supplemental Figure S3

A

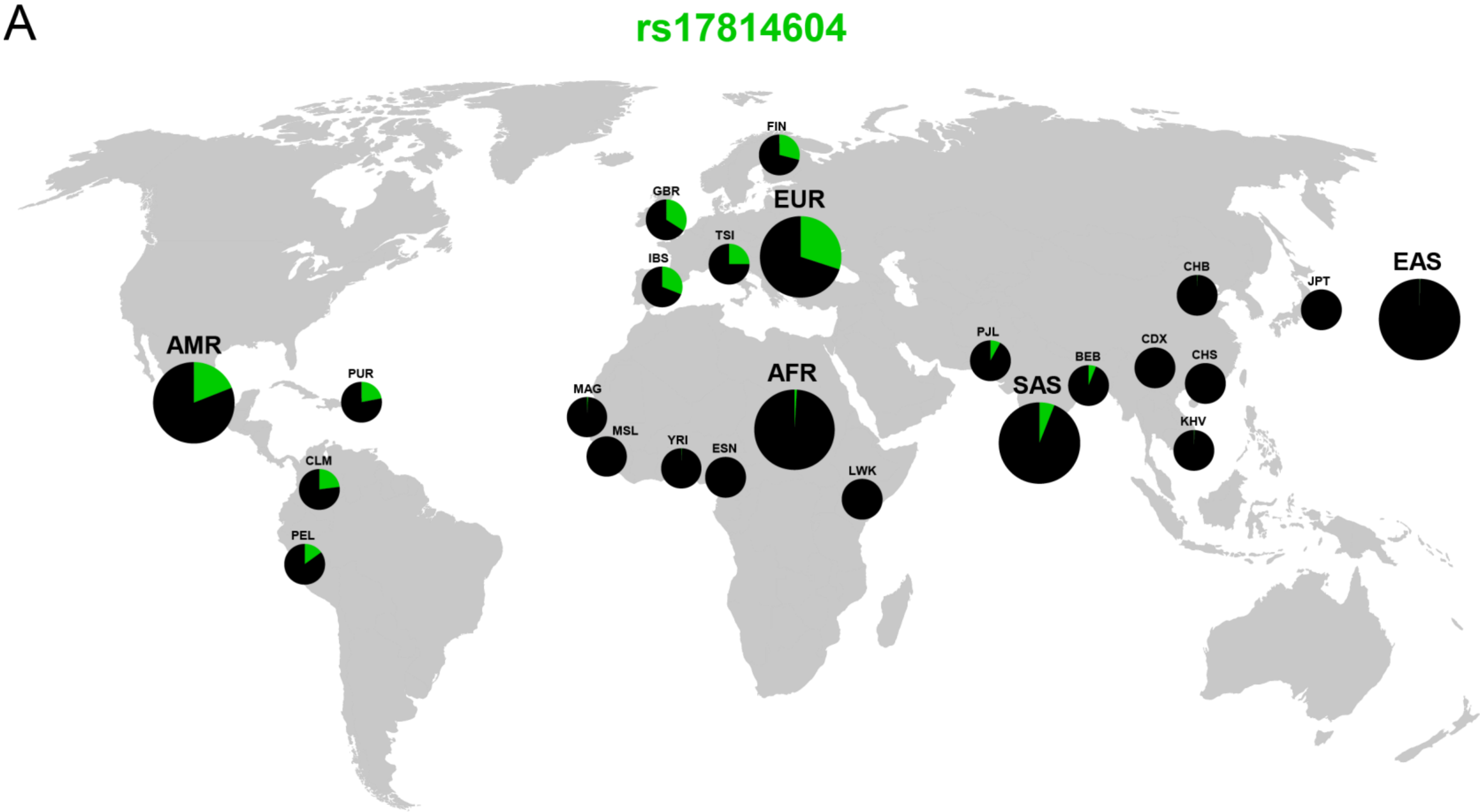


B

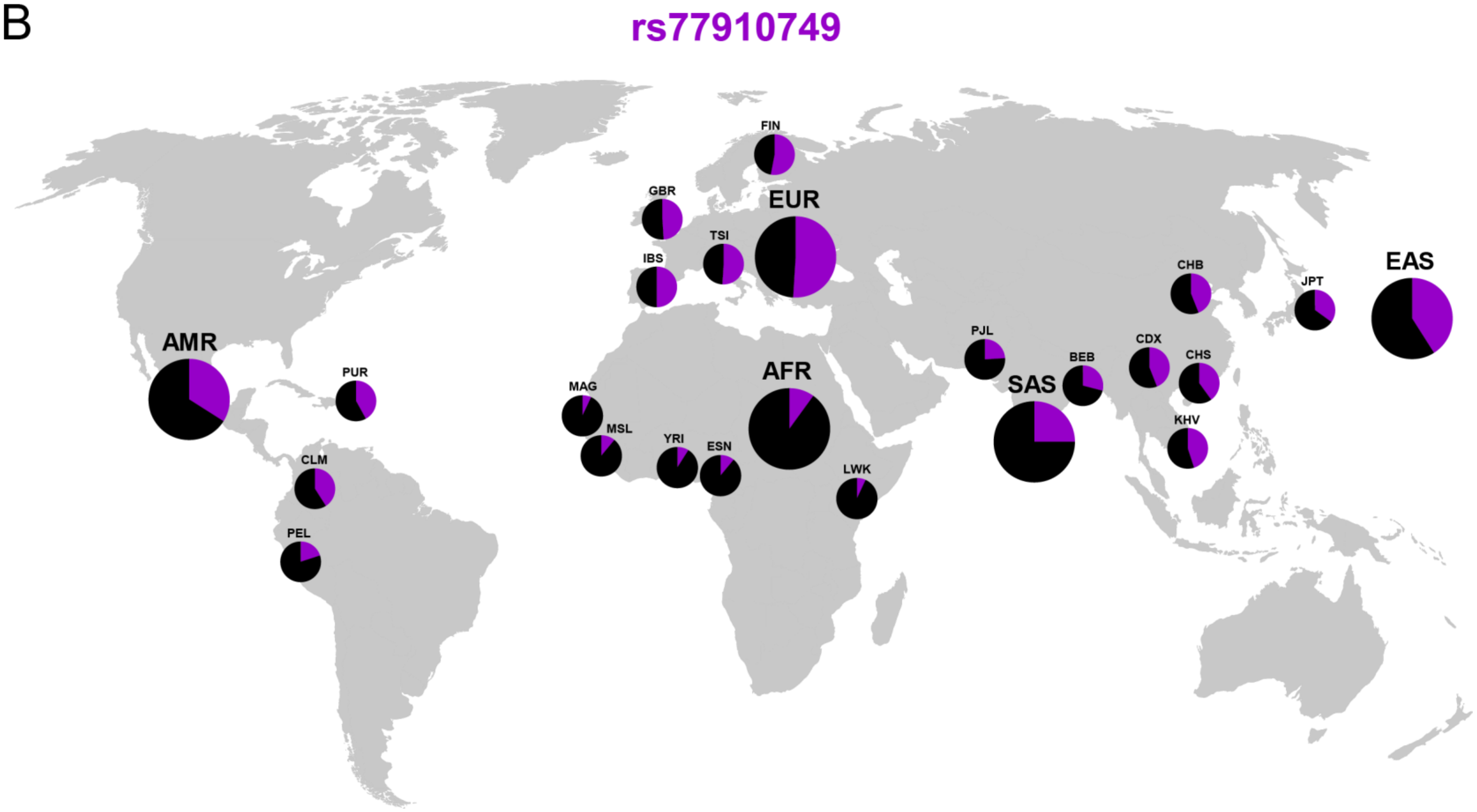

Supplemental Figure S4

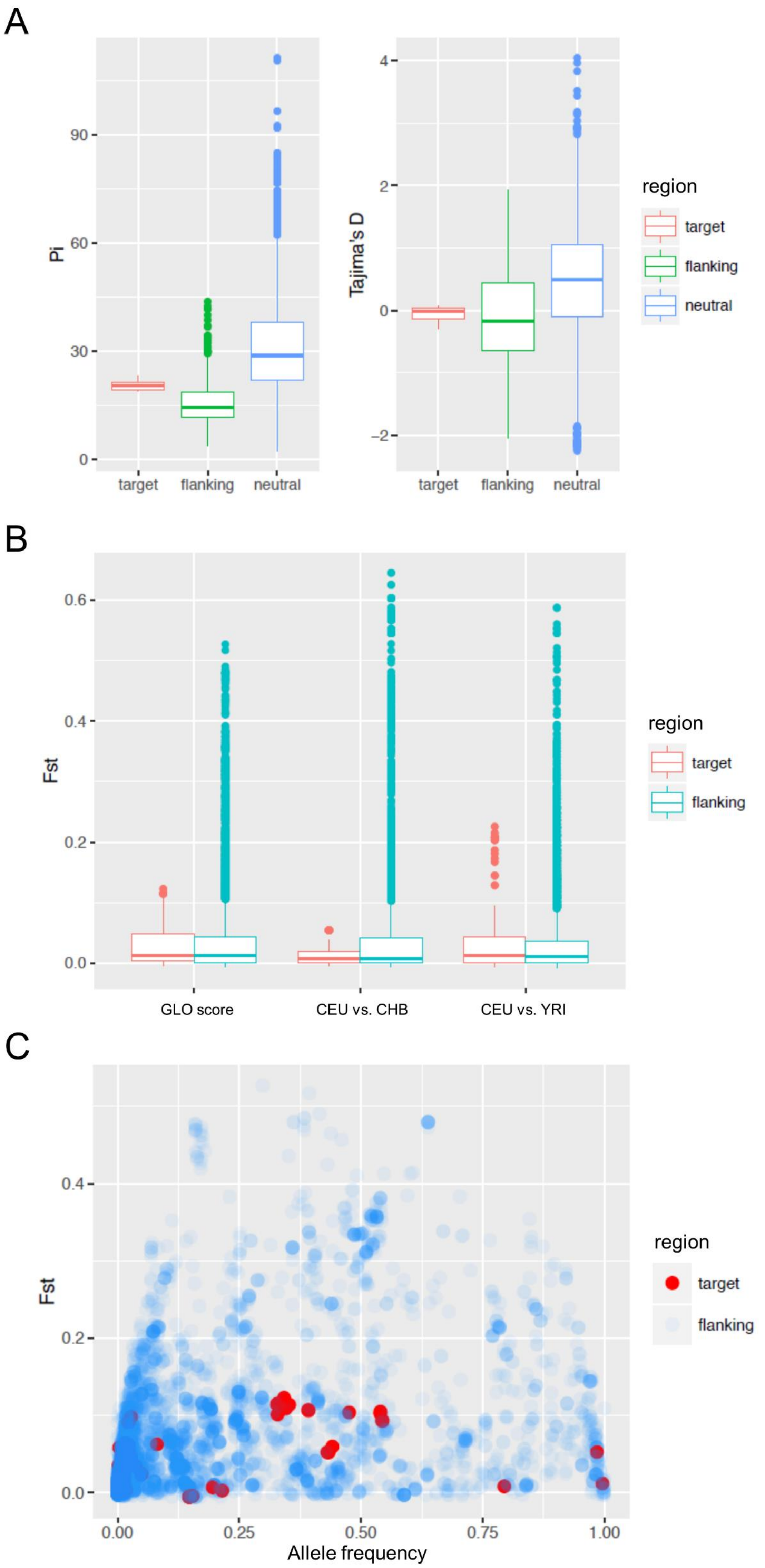

Supplemental Figure S5

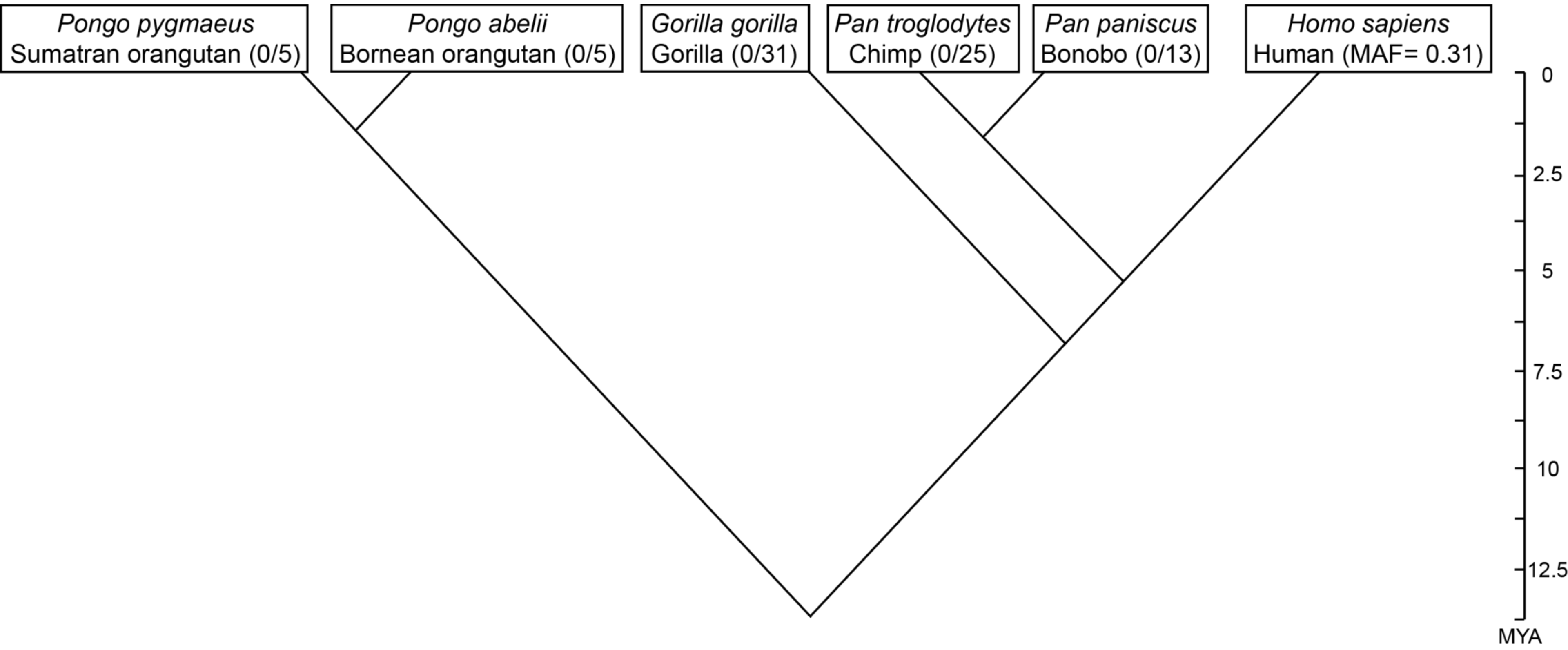

Supplemental Figure S6

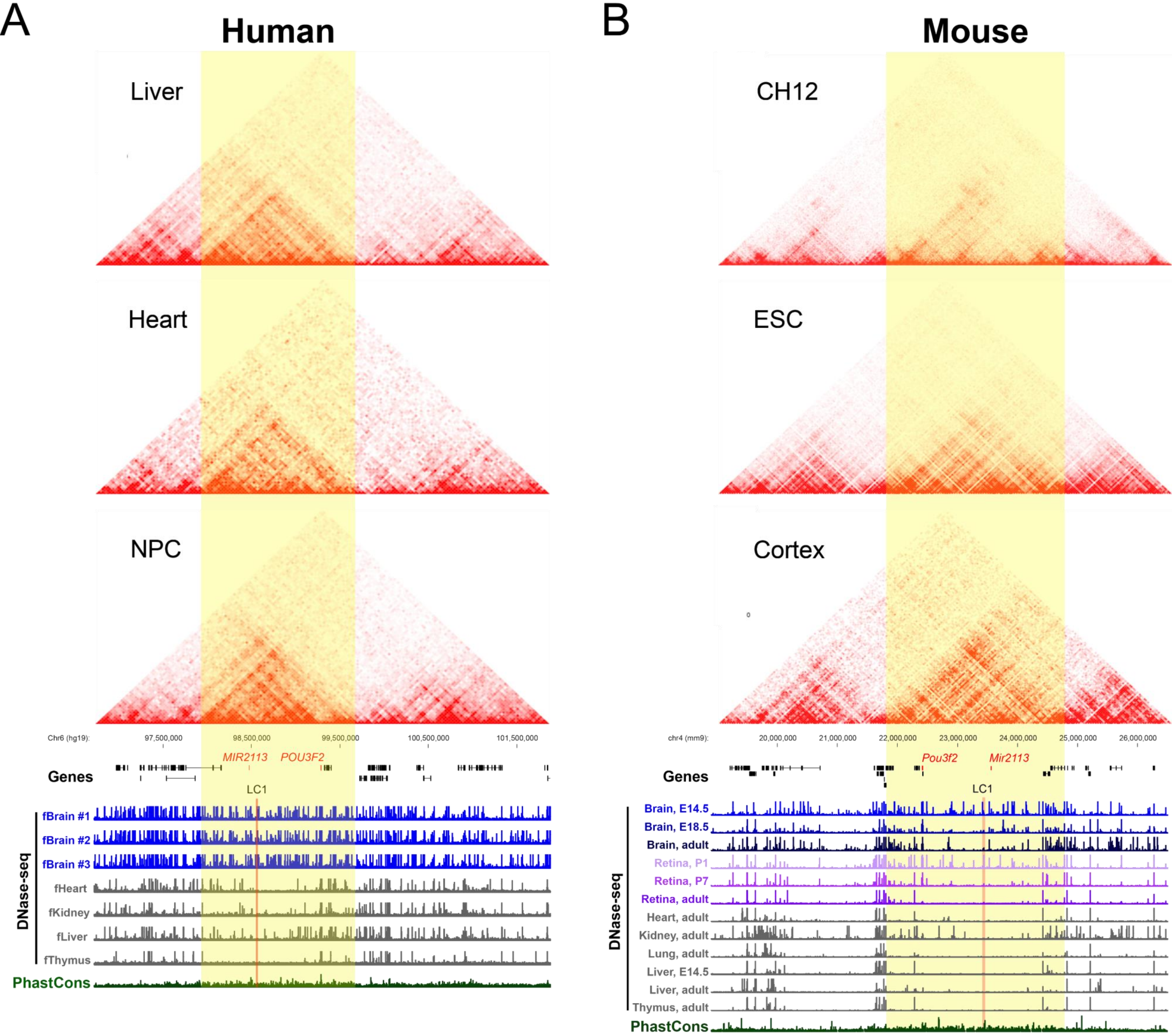

Supplemental Figure S7

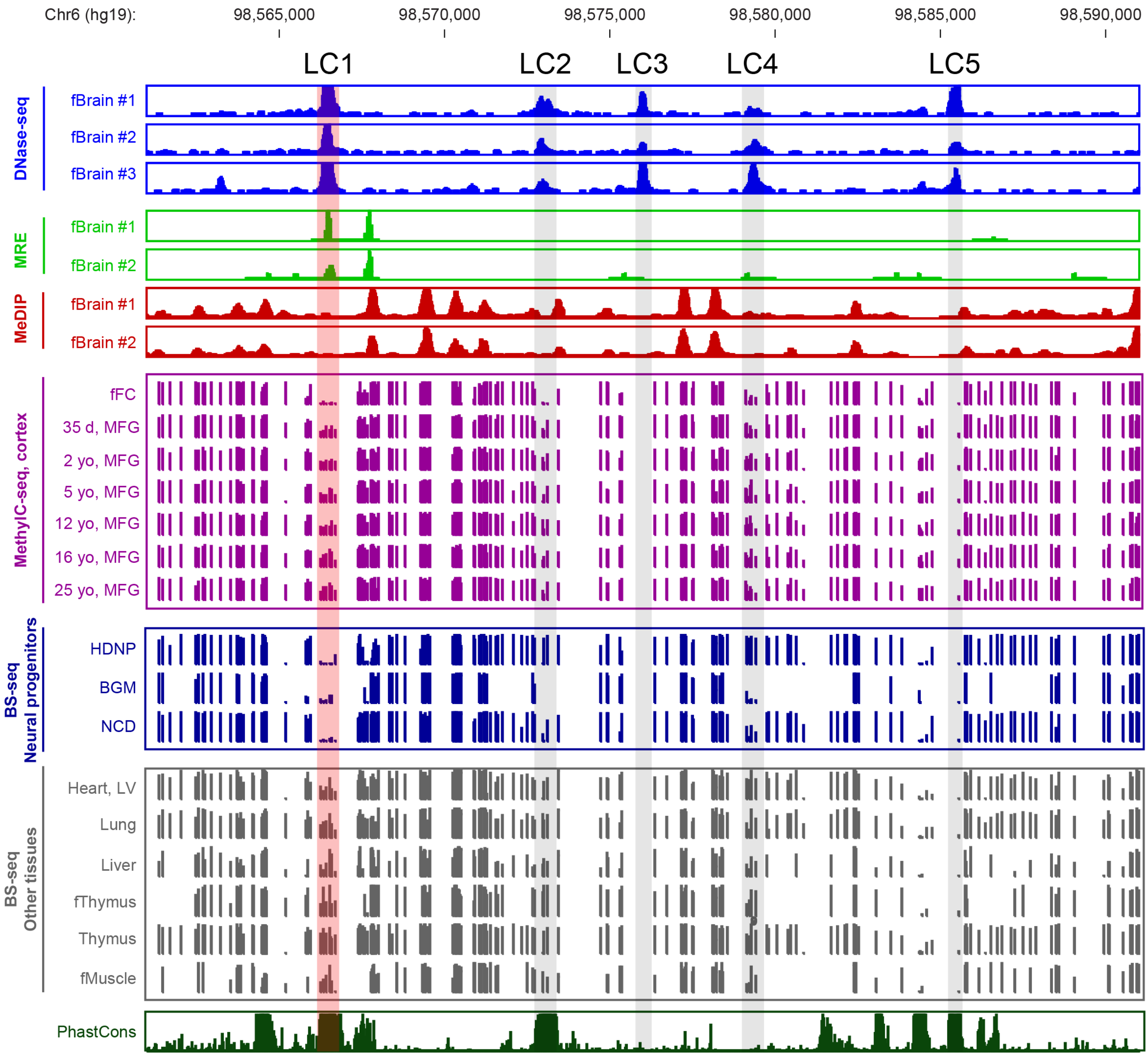

Supplemental Figure S8

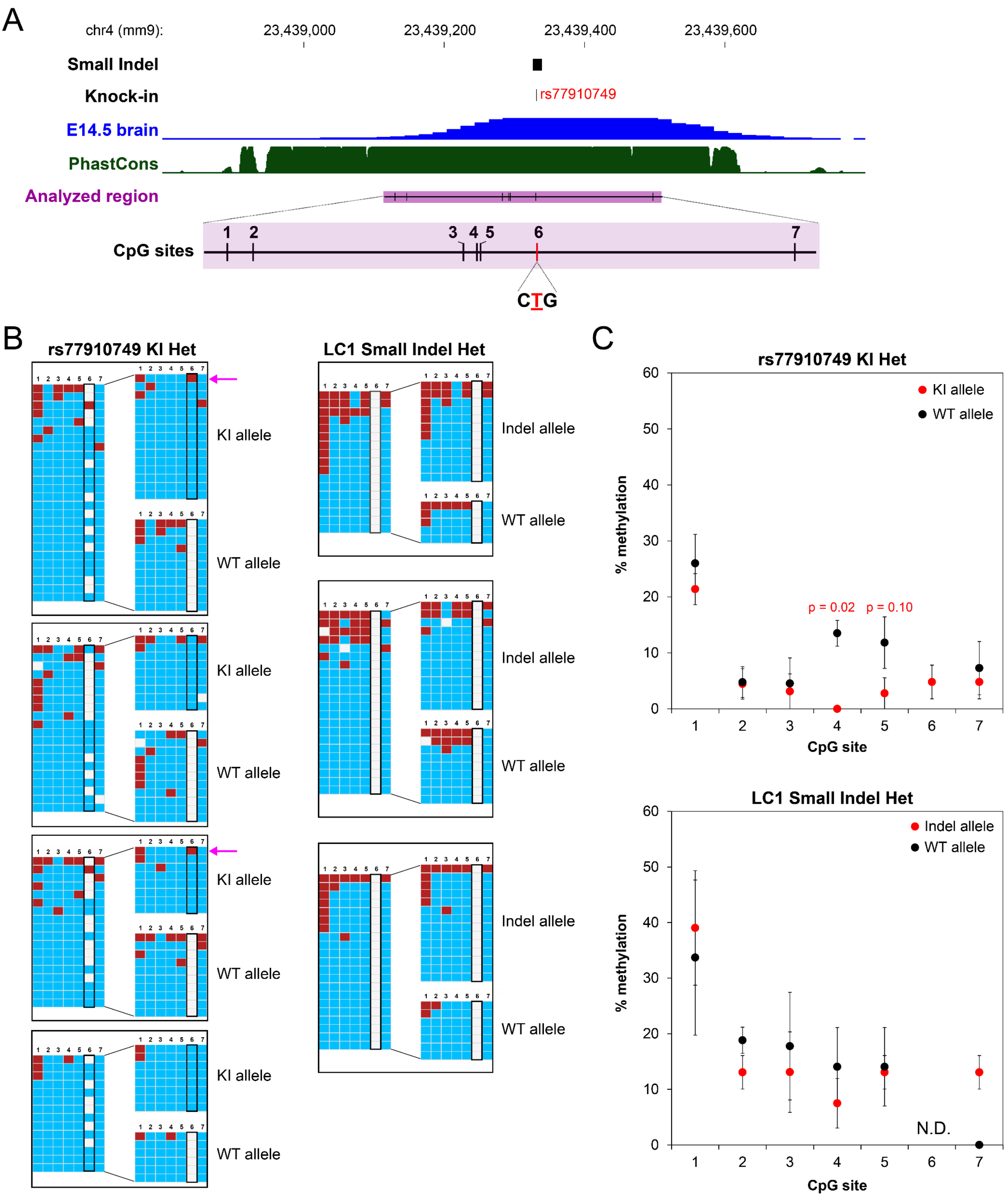

Supplemental Figure S9

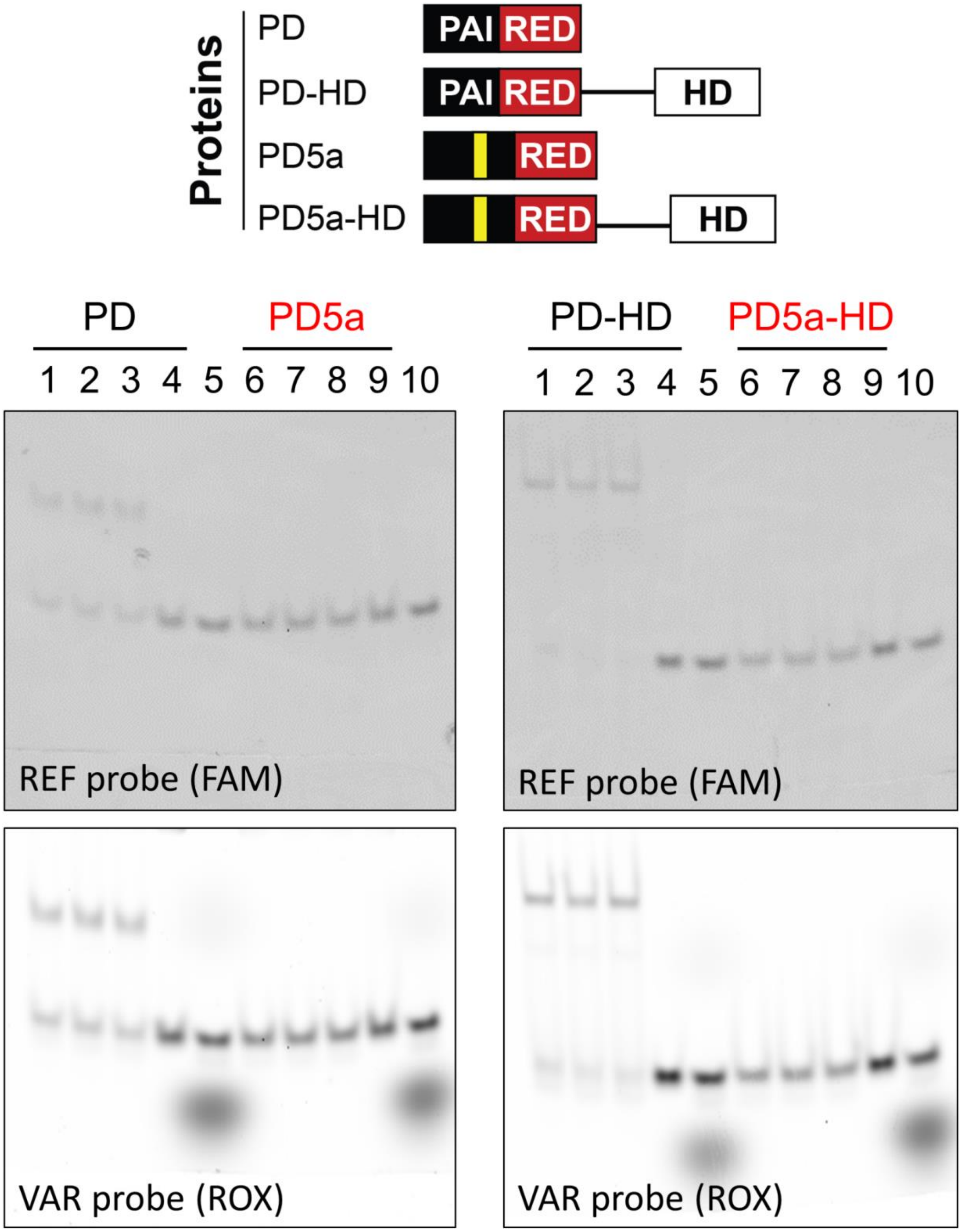

Supplemental Figure S10

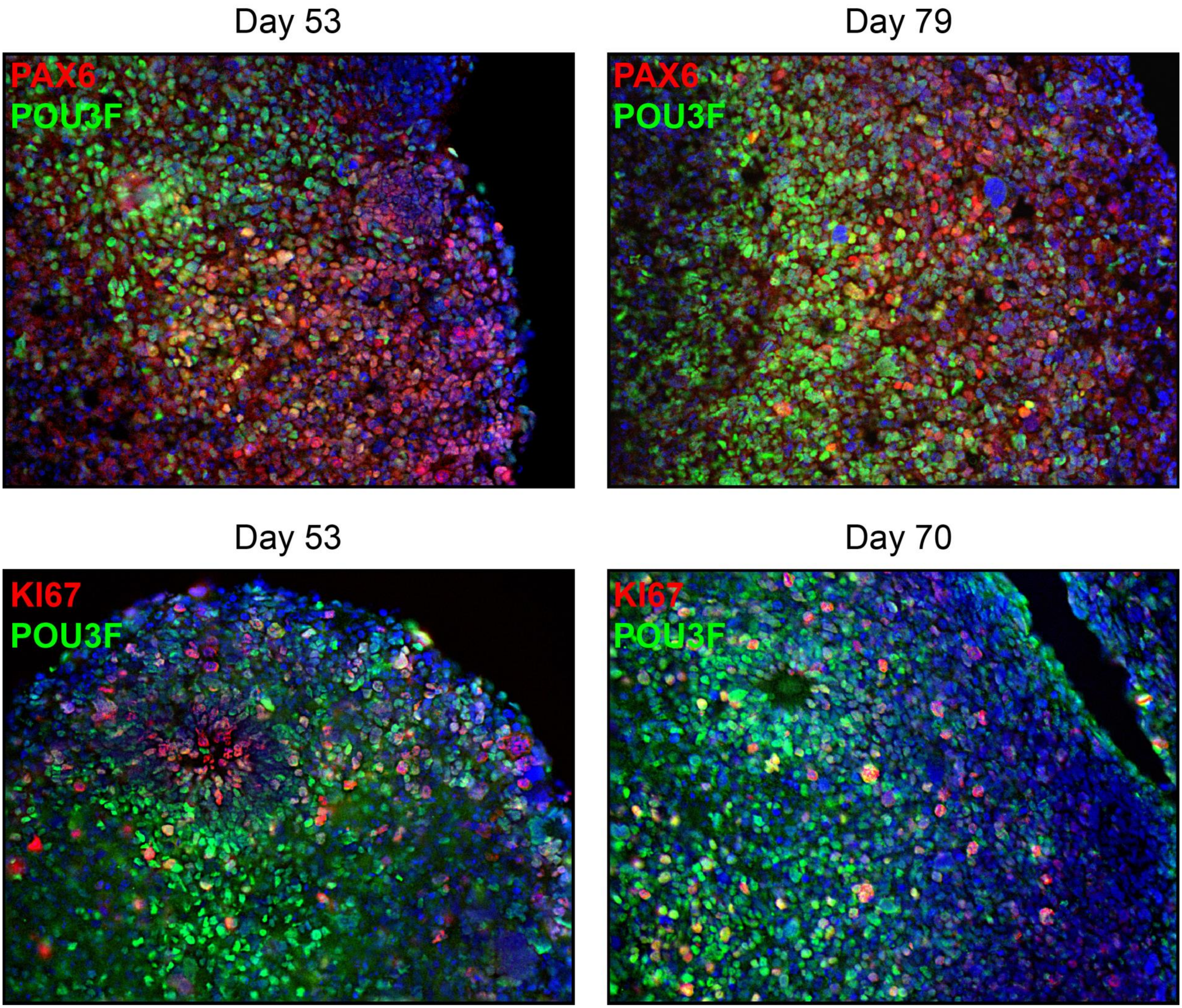
