## Supplemental File S1 for "A candidate causal variant underlying both enhanced cognitive performance and increased risk of bipolar disorder"

### 1-HR LOCOMOTOR ACTIVITY

##### AMBULATORY ACTIVITY

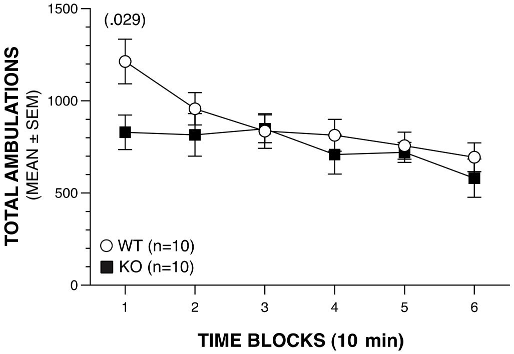


**No significant effects involving Genotype or Sex.**

##### VERTICAL REARING FREQUENCY

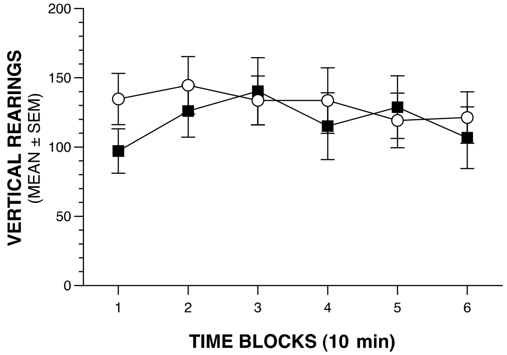


**No significant effects involving Genotype or Sex and no significant comparisons.**

##### DISTANCE IN PERIPHERY

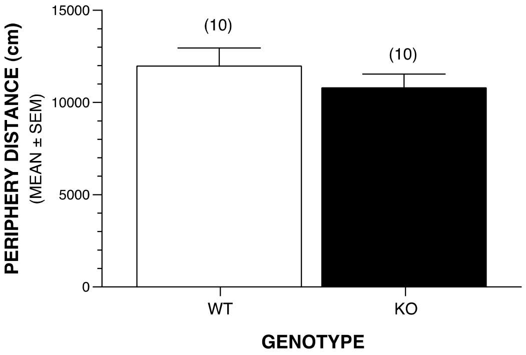


**No significant effects involving Genotype or Sex.**

##### ENTRIES INTO CENTER

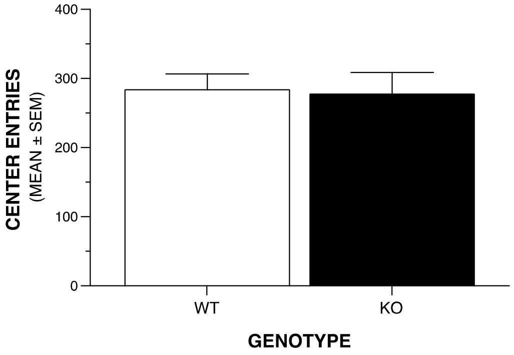


**No significant effects involving Genotype or Sex.**

##### TIME IN CENTER

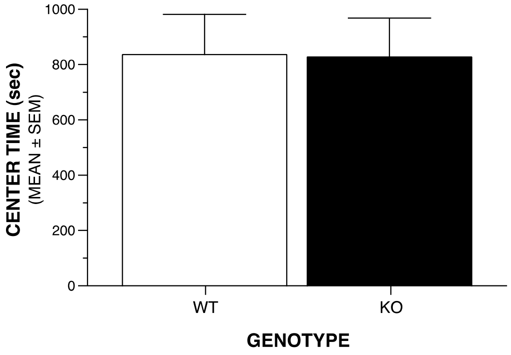


**No significant effects involving Genotype or Sex.**

##### DISTANCE IN CENTER

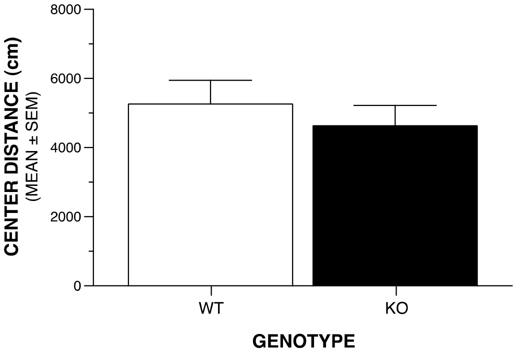


**No significant effects involving Genotype or Sex.**

### 1-HR LOCOMOTOR ACTIVITY: FIRST 5 MIN.

#### DISTANCE IN PERIPHERY

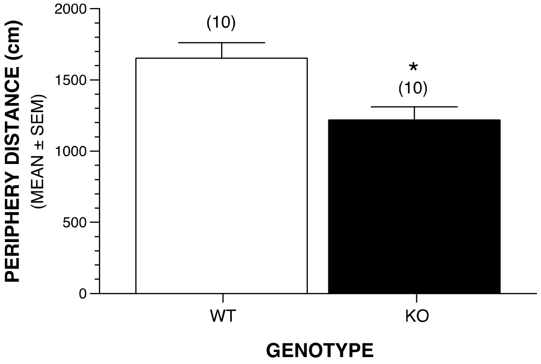


Significant Genotype effect: *p=010.

#### DISTANCE IN CENTER

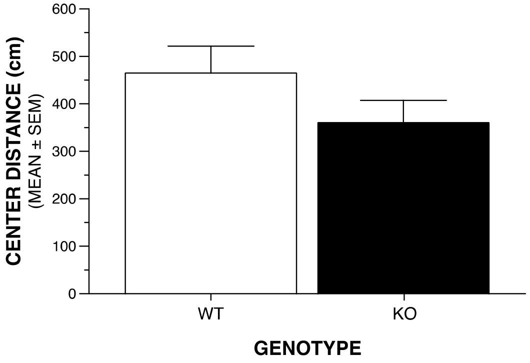


No significant effects involving Genotype or Sex.

#### TIME IN CENTER

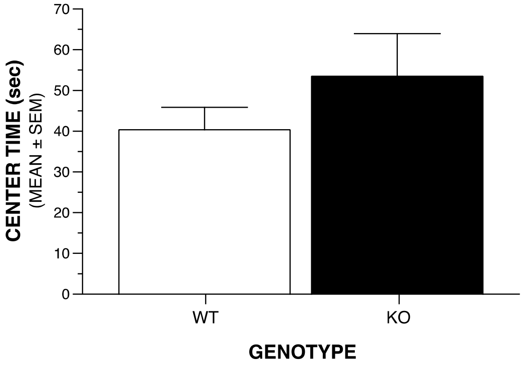


No significant effects involving Genotype or Sex.

#### ENTRIES INTO CENTER

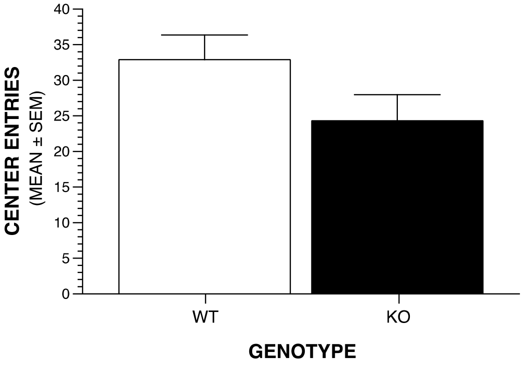


No significant effects involving Genotype or Sex.

### SENSORIMOTOR BATTERY

#### WALKING INITIATION

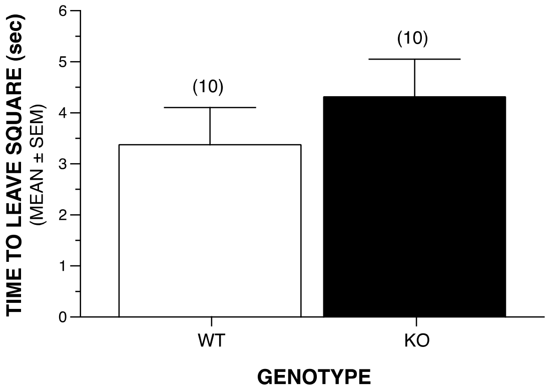


**No significant effects involving Genotype or Sex.**

#### LEDGE

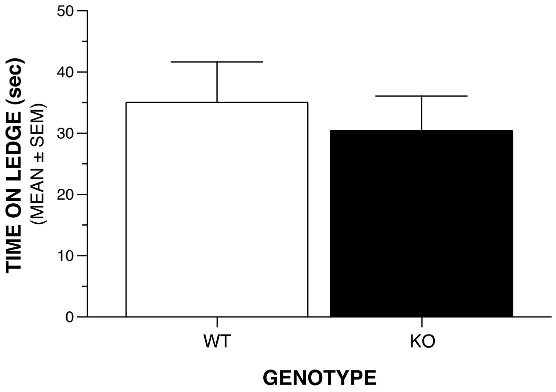


**No significant effects involving Genotype or Sex.**

#### PLATFORM

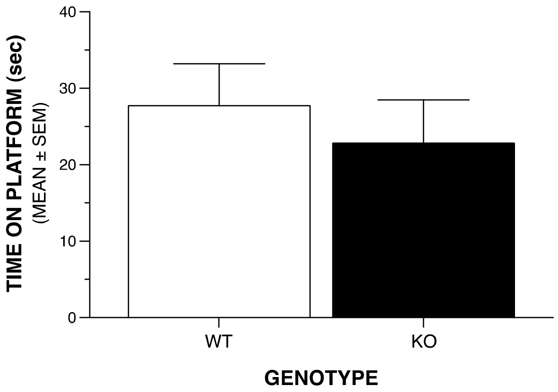


**No significant effects involving Genotype or Sex.**

#### POLE

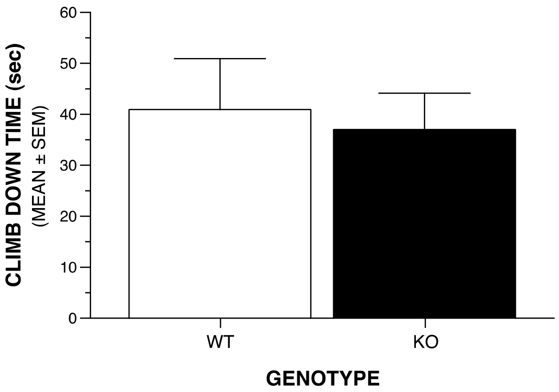


**No significant effects involving Genotype or Sex.**

### SENSORIMOTOR BATTERY: SCREEN TESTS

#### 60° INCLINED SCREEN

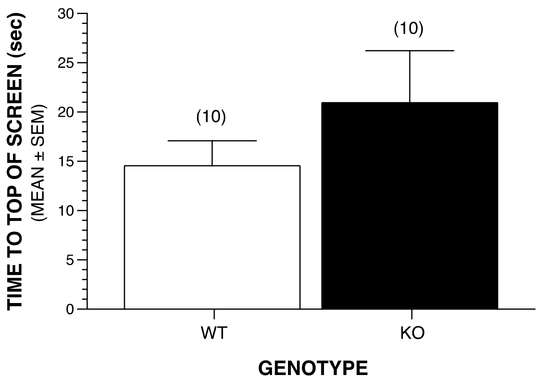


**No significant effects involving Genotype or Sex.**

#### 90° INCLINED SCREEN

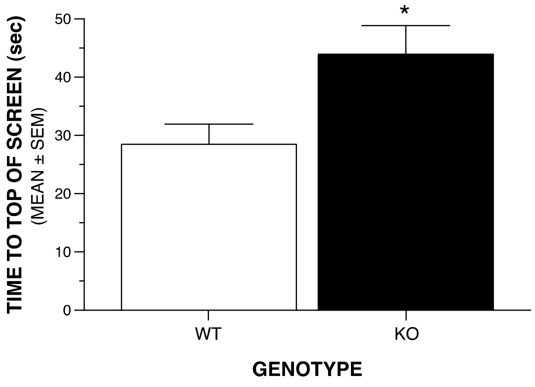


**Significant Genotype effect (*p=.015).**
**Sex effect: p=.066.**

#### INVERTED SCREEN

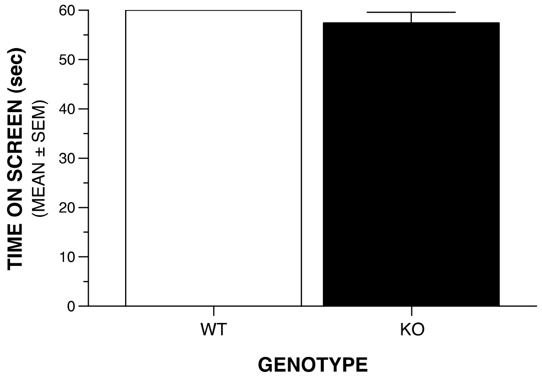


**No significant effects involving Genotype or Sex.**

### MORRIS WATER MAZE: CUED & PLACE TRIALS

#### CUED TRIALS: PATH LENGTH

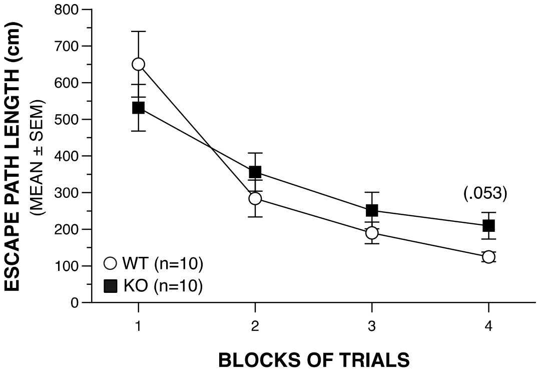


**No significant effects involving Genotype or Sex and no significant comparisons.**

#### CUED TRIALS: LATENCY

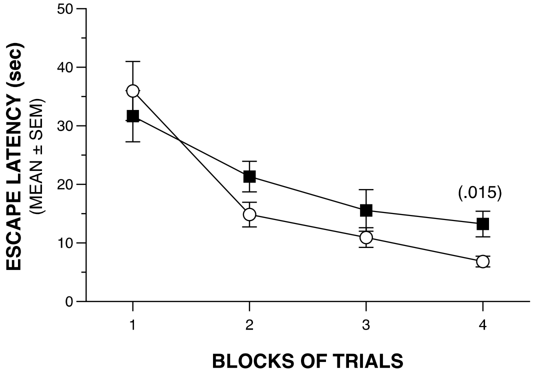


**No significant effects involving Genotype, but a significant Sex effect (p=.041).**

#### CUED TRIALS: SWIMMING SPEEDS

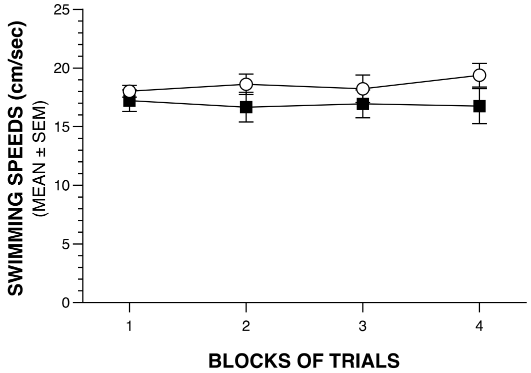


**Significant Sex effect (p=.0003), and Genotype x Sex interaction (p=.036).**

#### PLACE TRIALS: PATH LENGTH

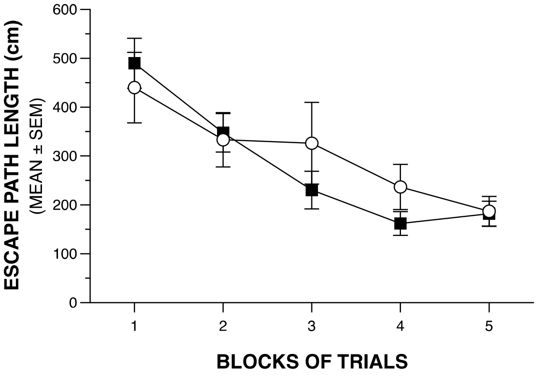


**No significant effects involving Genotype or Sex and no significant comparisons.**

#### PLACE TRIALS: LATENCY

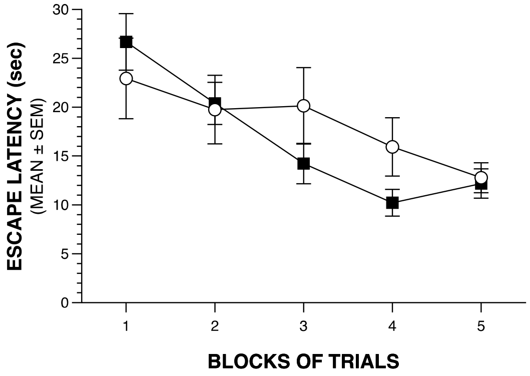


**No significant effects involving Genotype or Sex and no significant comparisons.**

#### PLACE TRIALS: SWIMMING SPEEDS

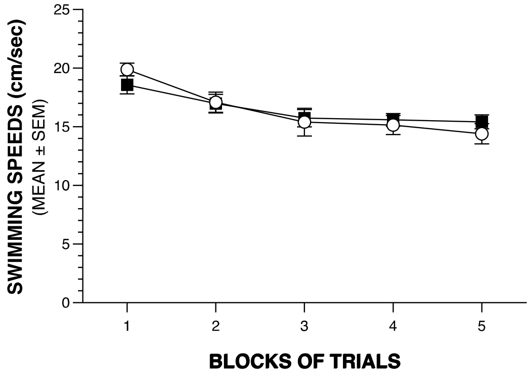


**No significant effects involving Genotype or Sex and no significant comparisons.**

### MORRIS WATER NAVIGATION: PROBE TRIAL

#### PLATFORM CROSSINGS

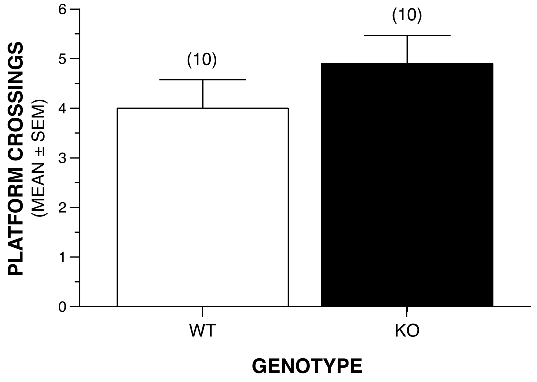


No significant effects involving Genotype or Sex.

#### TIME IN TARGET QUADRANT

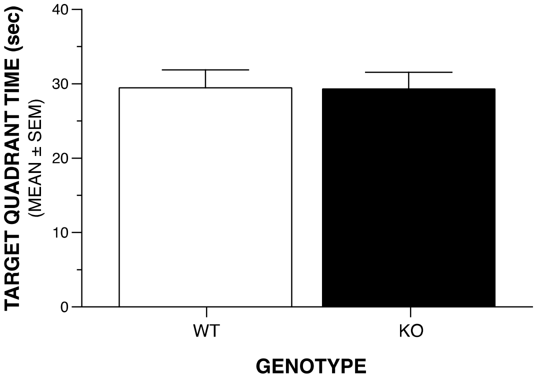


No significant effects involing Genotype or Sex.

#### SPATIAL BIAS

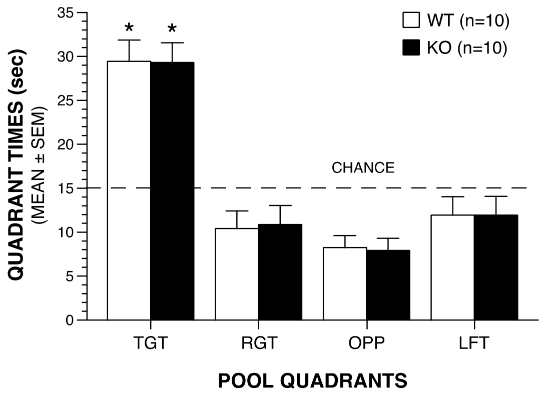


The WT and KO groups each showed spatial bias by spending significantly more time in the target quadrant vs each of the other quadrants (*p<.00005).

#### CONDITIONED FEAR

##### DAY 1: BASELINE - T/S TRAINING

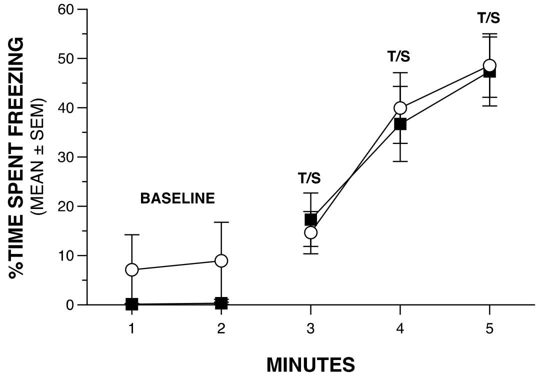


**BASELINE: no significant effects involving Genotype or Sex and no signficant comparisons.**

**T/S TRAINING: no significant effects involving Genotype but a significant Sex effect (p=.012). No significant between-groups comparisons.**

##### DAY 2: CONTEXTUAL FEAR

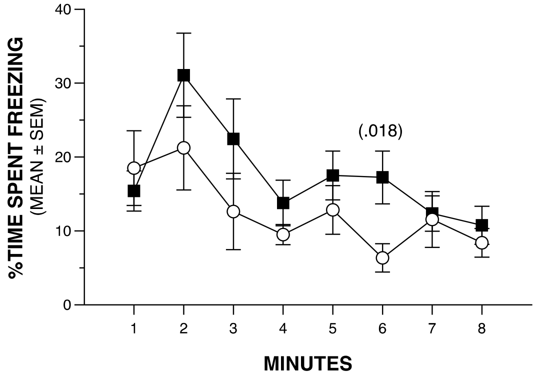


**No significant effects involving Genotype or Sex.**

##### DAY 3: AUDITORY CUE

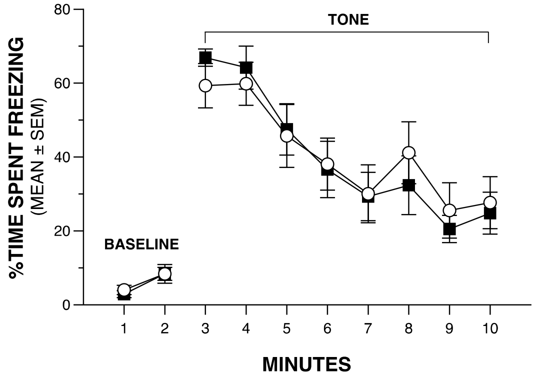


**BASELINE: no significant effects involving Genotype or Sex and no significant comparisons.**

**AUDITORY CUE: no significant effects involving Genotype, but a significant Sex x Minutes interaction (p=.003). No significant between-groups comparisons.**

### ACOUSTIC STARTLE/PPI & VOS

#### ACOUSTIC STARTLE

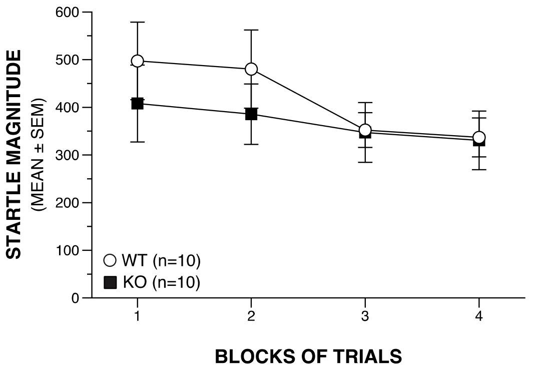


No significant effects involving Genotype or Sex and no significant comparsions.

#### % OF MICE SIGNIFICANTLY STARTLED

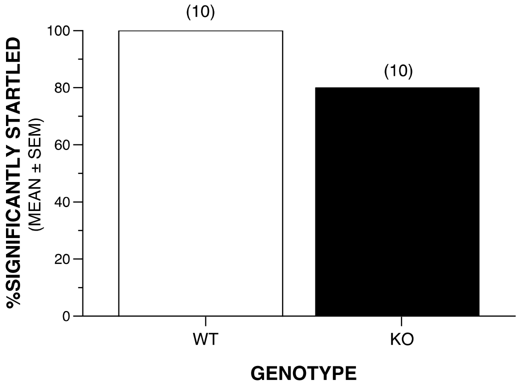


10/10 WT and 8/10 KO mice showed a significant startle response compared to levels during no stimulus trials.

#### UNTRANSFORMED PPI DATA

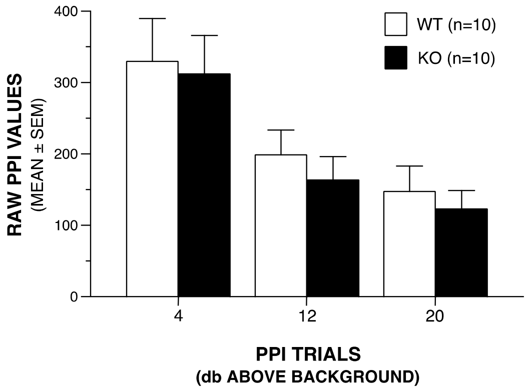


#### TOTAL MEAN %PPI

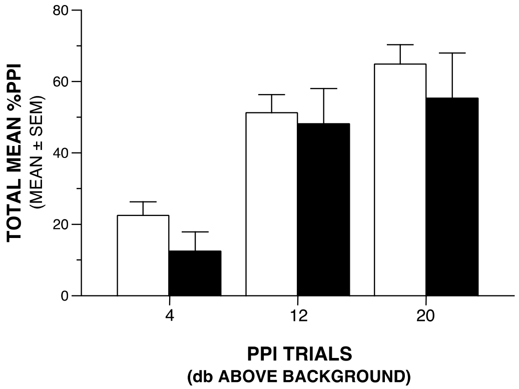


No significant effects involving Genotype and no significant comparisons.

#### BODY WEIGHTS AT TESTING

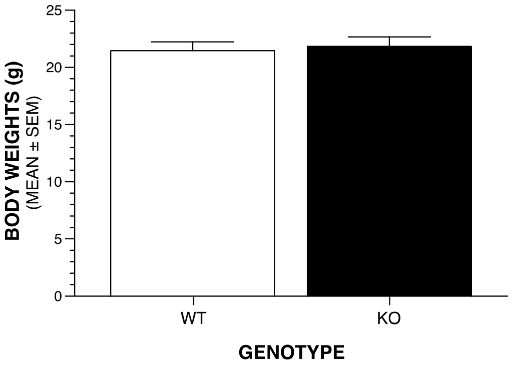


No significant effects involving Genotype but a significant Sex effect (p=.001).

### ELEVATED PLUS MAZE: OPEN ARM VARIABLES

#### ENTRIES INTO OPEN ARMS

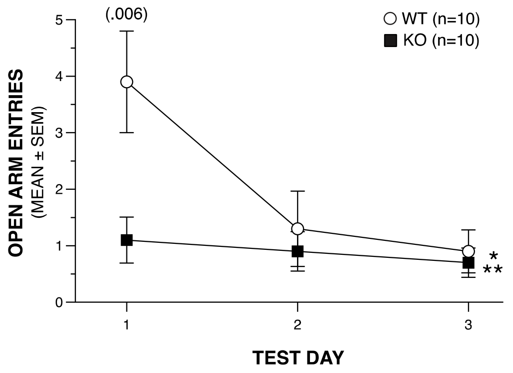


**Significant Genotype effect (*p=.049), and Genotype x Test Day (**p=.011) and Genotype x Sex x Test Day (p=.017) interactions.**

#### TIME IN OPEN ARMS

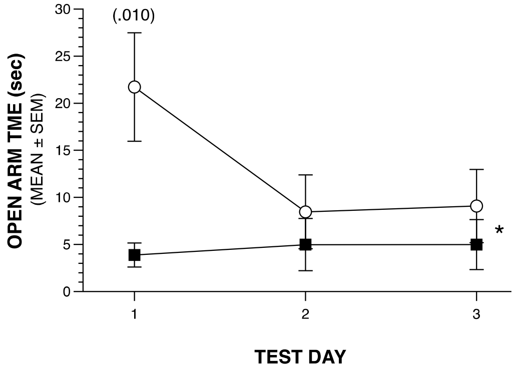


**Significant Genotype x Test Day interaction (*p=.038), but no significant effects involving Sex. Genotype effect: p=.051.**

#### DISTANCE TRAVELED IN OPEN ARMS

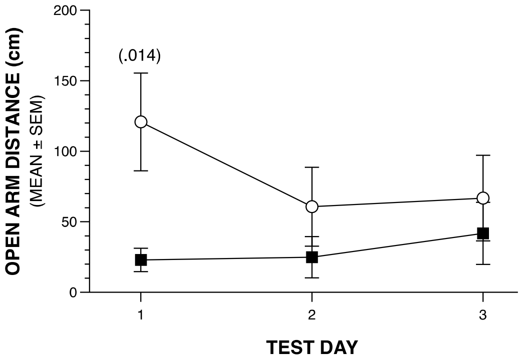


**No significant effects involving Genotype or Sex. Genotype effect: p=.062.**

#### % OF TOTAL ARM ENTRIES MADE INTO OPEN ARMS

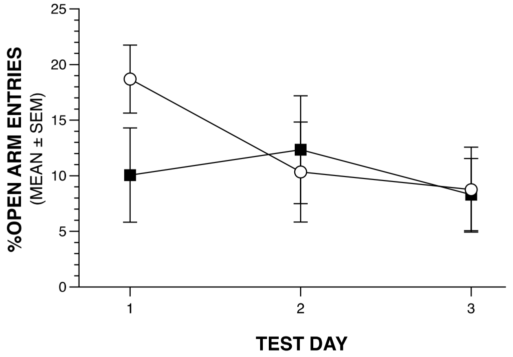


**No significant effects involving Genotype or Sex and no significant between-group comarisons.**

#### % OF TOTAL ARM TIME SPENT IN OPEN ARMS

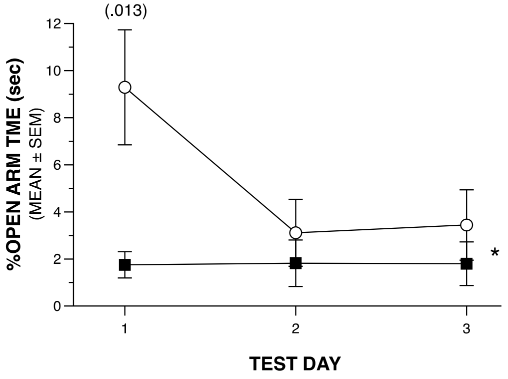


**Significant Genotype x Test Day interaction (*p=.031), but no significant effects involving Sex. Genotype effect: p=.052.**

#### % OF TOTAL ARM DISTANCE TRAVELED IN OPEN ARMS

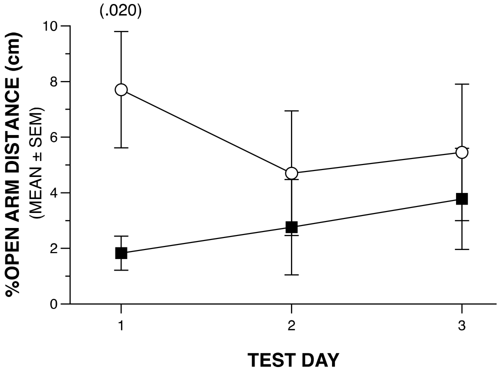


**No significant effects involving Genotype or Sex.**

#### ELEVATED PLUS MAZE

##### DISTANCE VARIABLES

###### TOTAL DISTANCE TRAVELED

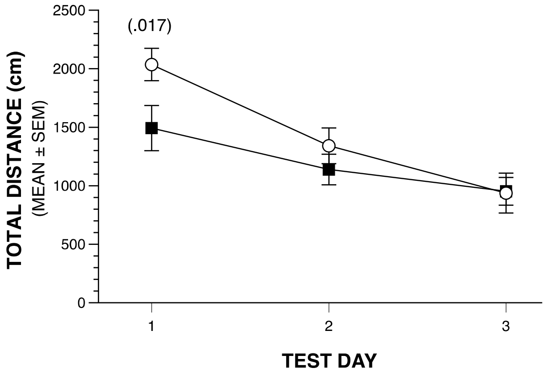


**No significant effects involving Genotype or Sex.**

###### DISTANCE TRAVELED IN CLOSED ARMS

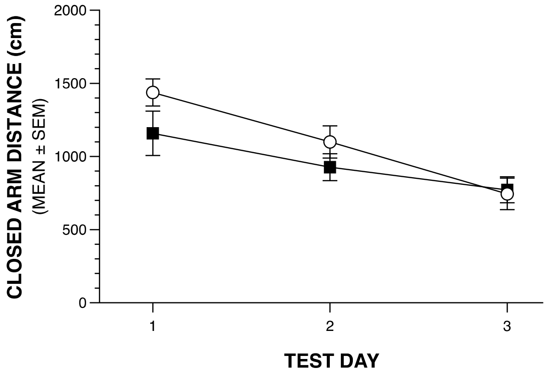


**No significant effects involving Genotype or Sex and no significant between-group comarisons.**

###### DISTANCE TRAVELED IN CENTER

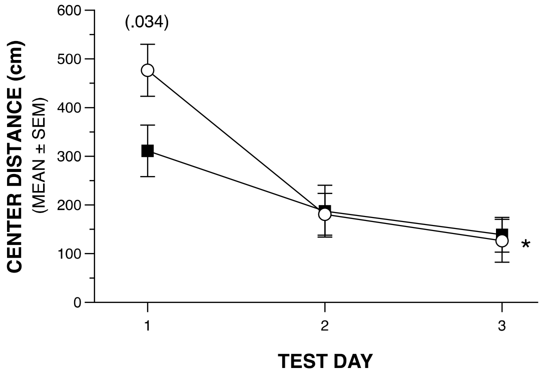


**Significant Genotype x Test Day (*p=.009), and Sex x Test Day (p=.045) interactions.**

###### DISTANCE TRAVELED IN OPEN ARMS

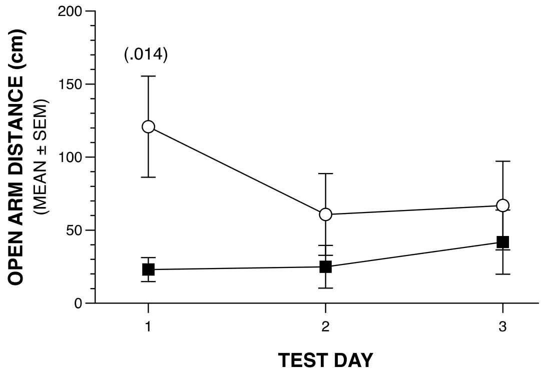


**No significant effects involving Genotype or Sex. Genotype effect: p=.062.**

### 1-HR OPEN-FIELD ACTIVITY/EXPLORATORY BEHAVIOR

#### ZONE ANALYSES

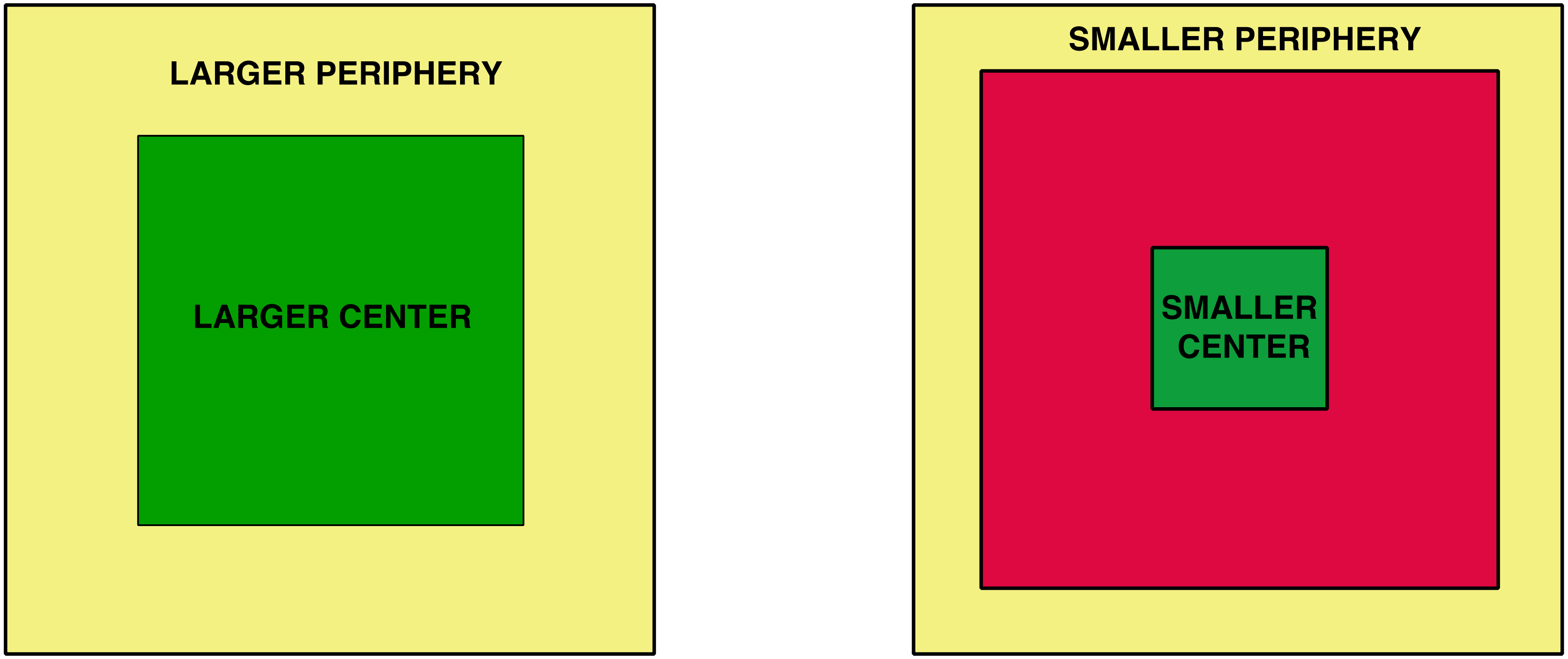

#### 1-HR OPEN-FIELD ACTIVITY/EXPLORATORY BEHAVIOR

##### DISTANCE IN “LARGER” PERIPHERY

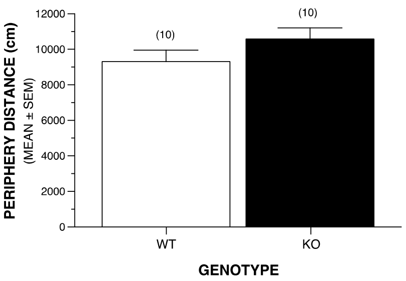


No significant effects involving Group or Sex.

##### DISTANCE IN “LARGER” CENTER

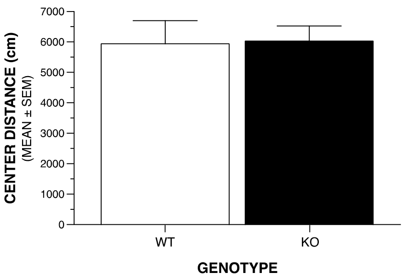


No significant effects involving Group or Sex.

##### TIME IN “LARGER” CENTER

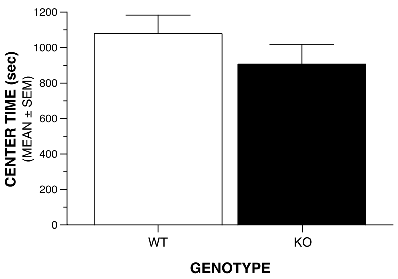


No significant effects involving Group or Sex.

##### ENTRIES INTO “LARGER” CENTER

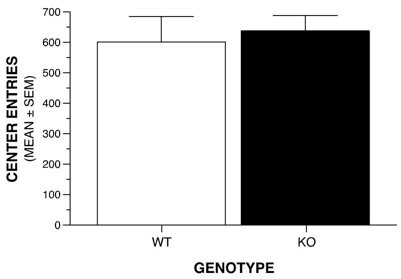


No significant effects involving Group or Sex.

##### DISTANCE IN “SMALLER” PERIPHERY

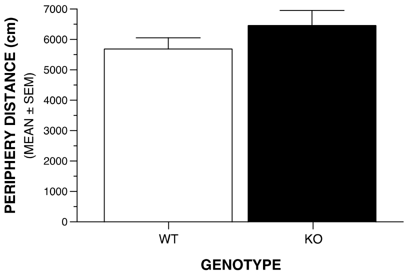


No significant effects involving Group or Sex.

##### DISTANCE IN “SMALLER” CENTER

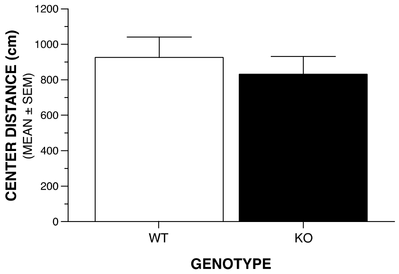


No significant effects involving Group or Sex.

##### TIME IN “SMALLER” CENTER

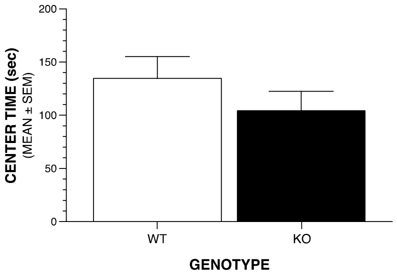


No significant effects involving Group or Sex.

##### ENTRIES INTO “SMALLER” CENTER

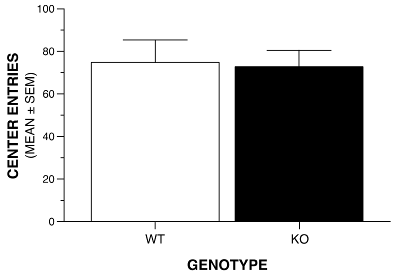


No significant effects involving Group or Sex.

#### 1-HR OPEN-FIELD ACTIVITY/EXPLORATORY BEHAVIOR

##### AMBULATORY ACTIVITY

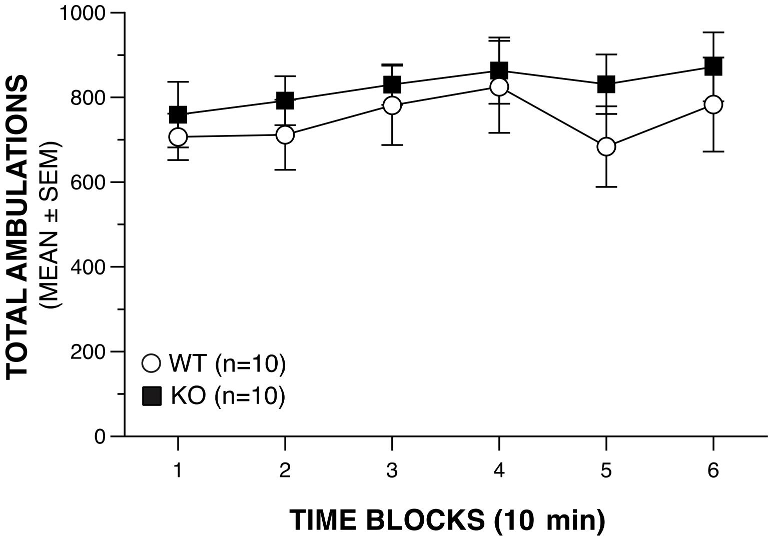


**No significant effects involving Genotype or Sex and no significant comparisons.**

##### VERTICAL REARING FREQUENCY

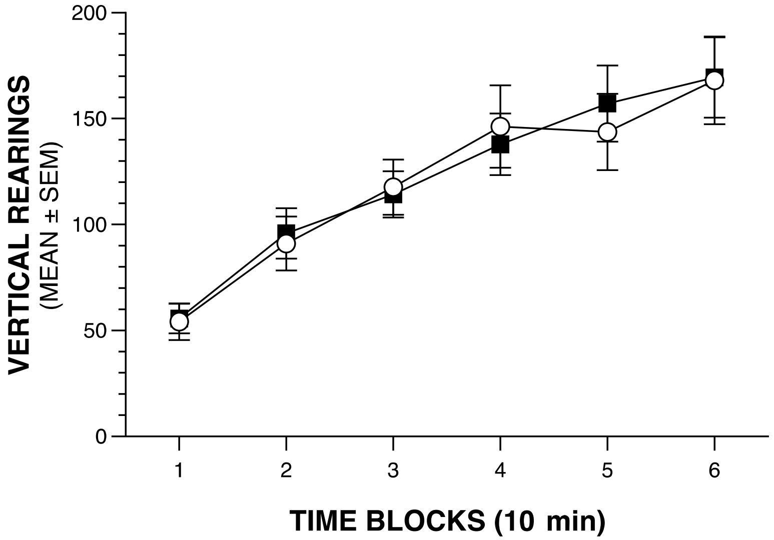


**No significant effects involving Genotype or Sex and no significant comparisons.**
