## Supplemental Files S2 for "A candidate causal variant underlying both enhanced cognitive performance and increased risk of bipolar disorder"

### 1-HR LOCOMOTOR ACTIVITY

#### AMBULATORY ACTIVITY

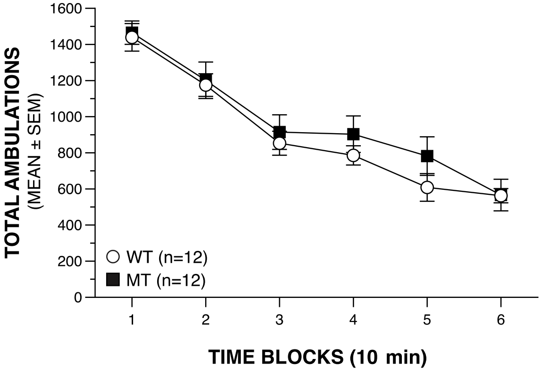


No significant effects involving Genotype or Sex and no significant between-groups comparisons.

#### REARING FREQUENCY

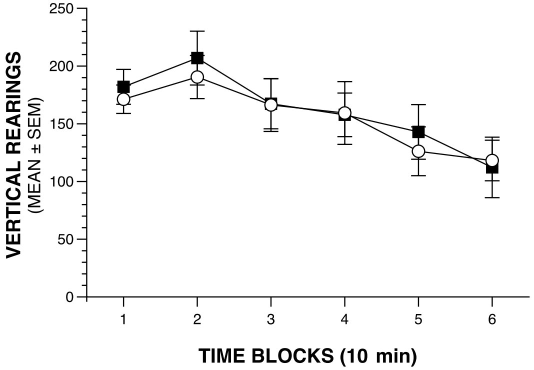


No significant effects involving Genotype, but a significant Sex x Time interaction ($p=.018$). No significant between-groups comparisons.

#### DISTANCE IN PERIPHERY

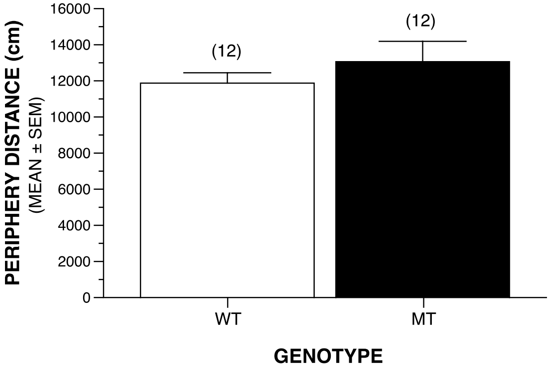


No significant effects involving Genotype or Sex.

#### ENTRIES INTO CENTER

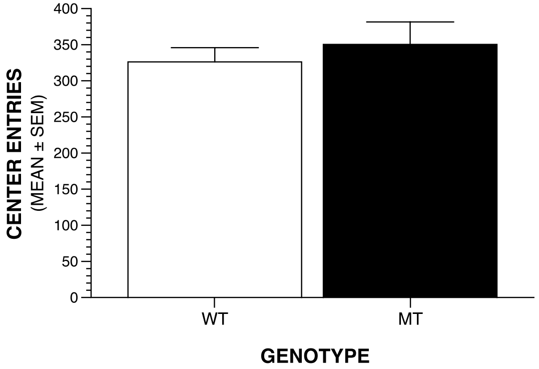


No significant effects involving Genotype or Sex.

#### TIME IN CENTER

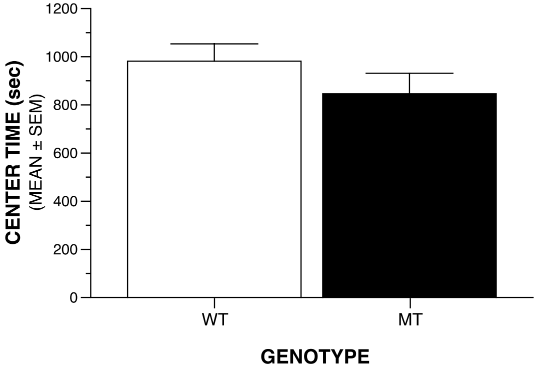


No significant effects involving Genotype or Sex.

#### DISTANCE IN CENTER

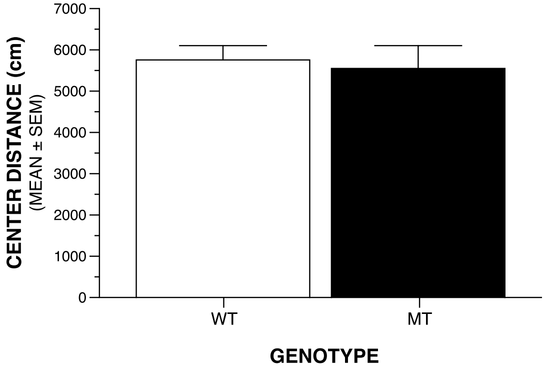


No significant effects involving Genotype or Sex.

#### SENSORIMOTOR BATTERY

##### WALKING INITIATION

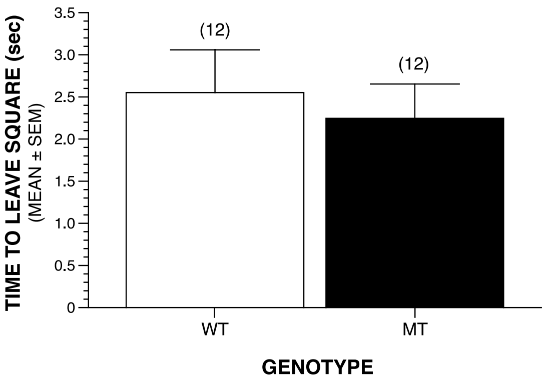


**No significant effects involving Genotype or Sex.**

##### LEDGE

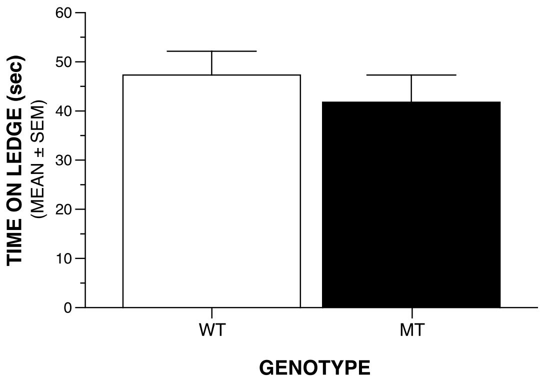


**No significant effects involving Genotype or Sex.**

##### PLATFORM

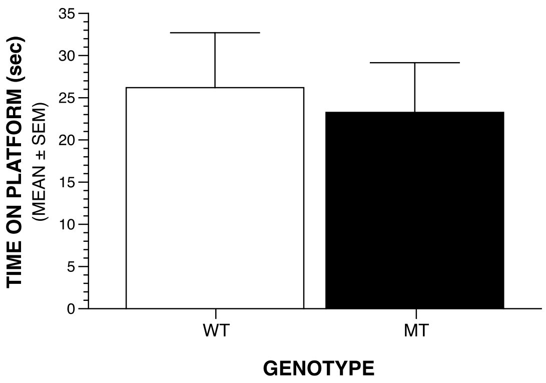


**No significant effects involving Genotype or Sex.**

##### POLE

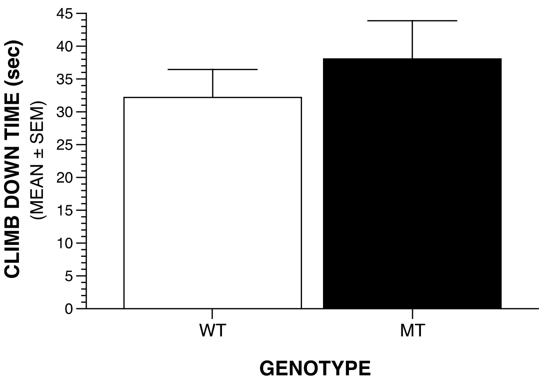


**No significant effects involving Genotype or Sex.**

#### SENSORIMOTOR BATTERY: SCREEN TESTS

##### 60º INCLINED SCREEN

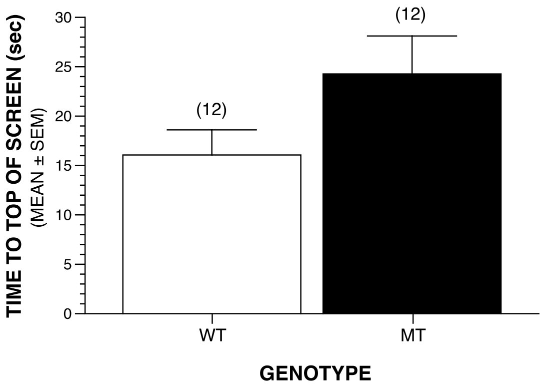


**No significant effects involving Genotype or Sex. Genotype effect: p=.071.**

##### 90º INCLINED SCREEN

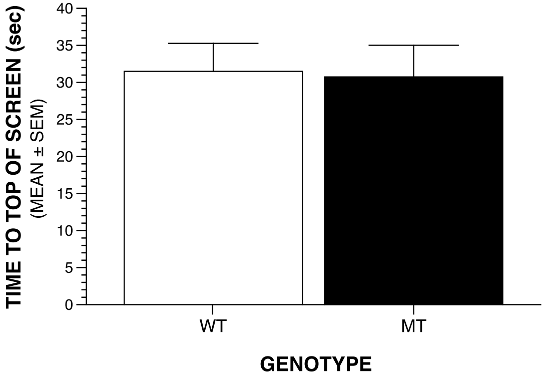


**No significant effects involving Genotype or Sex.**

##### INVERTED SCREEN

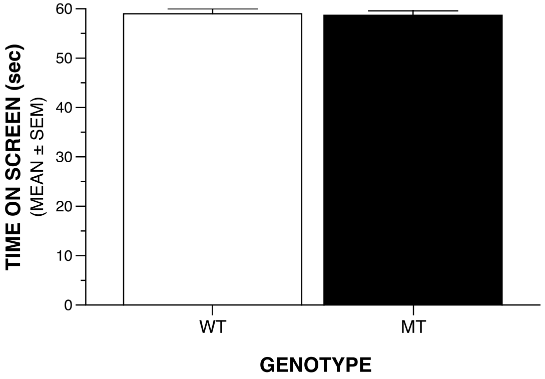


**No significant effects involving Genotype or Sex.**

### MORRIS WATER MAZE: CUED & PLACE TRIALS

##### CUED TRIALS: PATH LENGTH

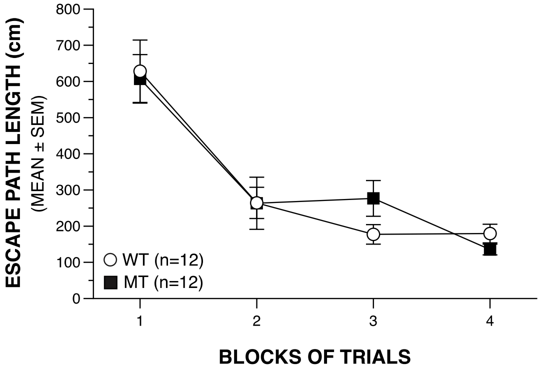


**No significant effects involving Genotype or Sex and no significant comparisons.**

##### CUED TRIALS: LATENCY

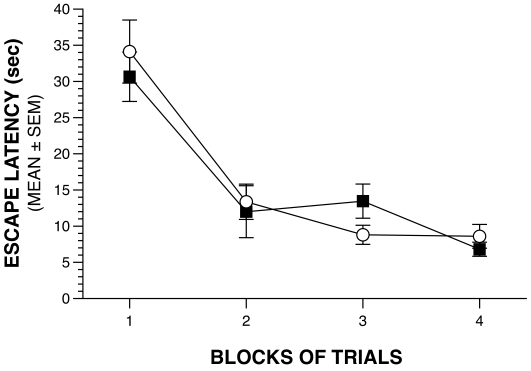


**No significant effects involving Group or Sex and no significant comparisons.**

##### CUED TRIALS: SWIMMING SPEEDS

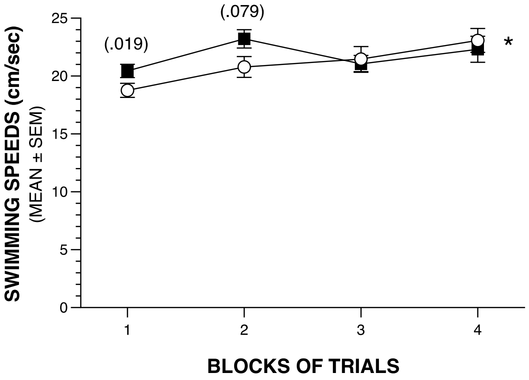


**Significant Sex effect (p=.018), and Group x Blocks of Trials interaction (*p=.035).**

##### PLACE TRIALS: PATH LENGTH

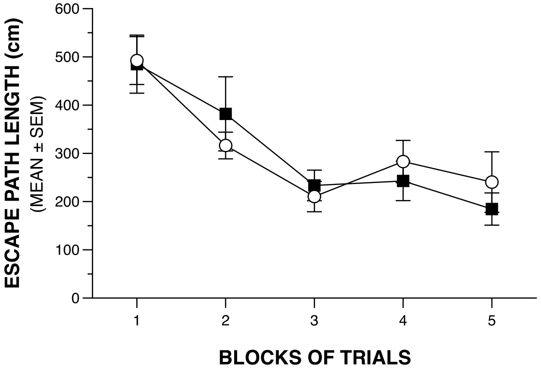


**No significant effects involving Genotype or Sex and no significant comparisons.**

##### PLACE TRIALS: LATENCY

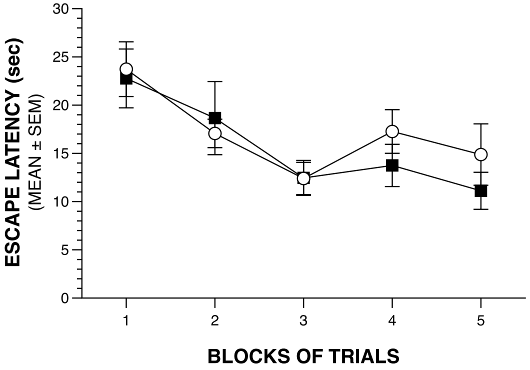


**No significant effects involving Genotype or Sex and no significant comparisons.**

##### PLACE TRIALS: SWIMMING SPEEDS

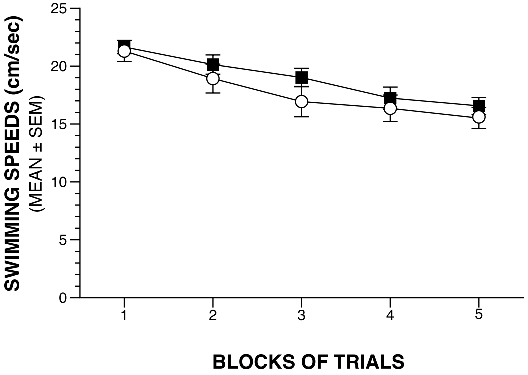


**No significant effects involving Genotype, but a significant Sex effect (p=.018).**

### MORRIS WATER NAVIGATION: PROBE TRIAL

##### PLATFORM CROSSINGS

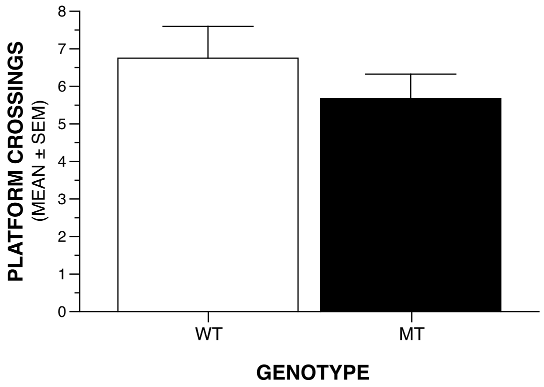


**No significant effects involving Genotype or Sex.**

##### TIME IN TARGET QUADRANT

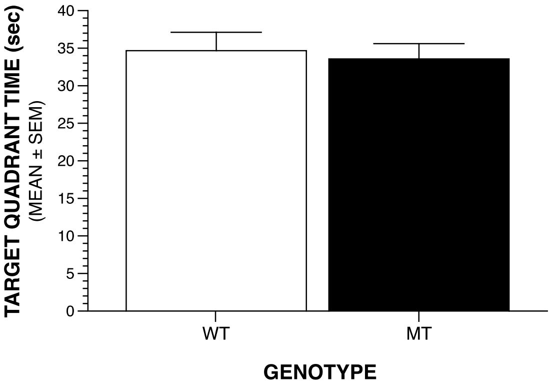


**No significant effects involving Genotype or Sex.**

##### SPATIAL BIAS

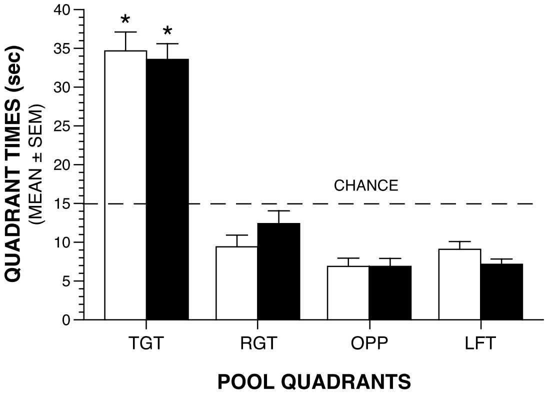


**The WT and MT groups each showed spatial bias for the target quadrant by spending significantly (*p<.00005) more time in the target quadrant vs each of the other quadrants.**

### ELEVATED PLUS MAZE: OPEN ARM VARIABLES

##### TIME IN OPEN ARMS

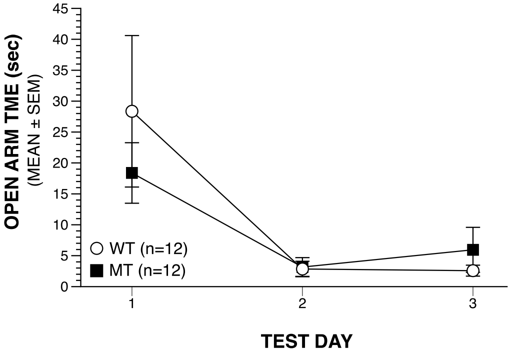


**No significant effects involving Genotype or Sex and no significant comparisons.**

##### ENTRIES INTO OPEN ARMS

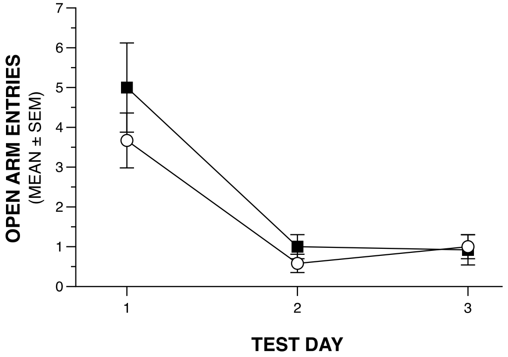


**No significant effects involving Genotype or Sex and no significant comparisons.**

##### DISTANCE TRAVELED IN OPEN ARMS

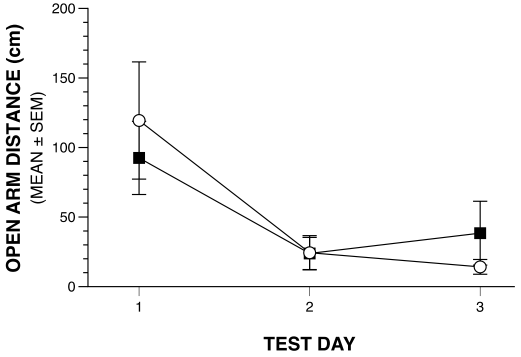


**No significant effects involving Genotype or Sex and no significant comparisons.**

##### % OF TOTAL ARM TIME SPENT IN OPEN ARMS

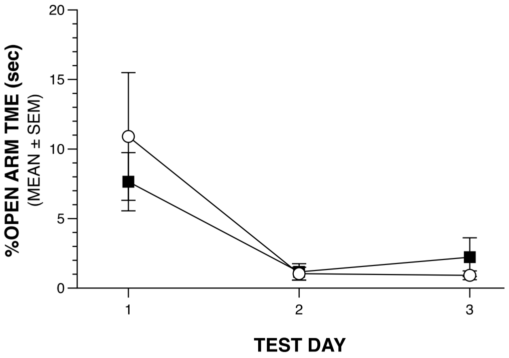


**No significant effects involving Genotype or Sex and no significant comparisons.**

##### % OF TOTAL ARM ENTRIES MADE INTO OPEN ARMS

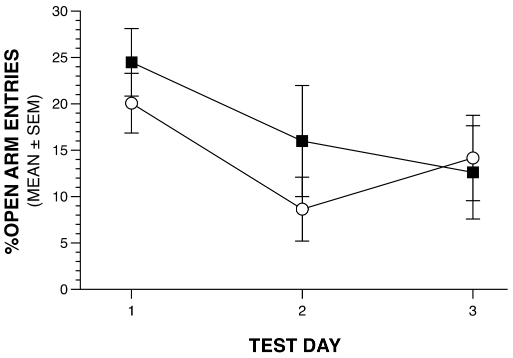


**No significant effects involving Genotype or Sex and no significant comparisons.**

##### % OF TOTAL ARM DISTANCE TRAVELED IN OPEN ARMS

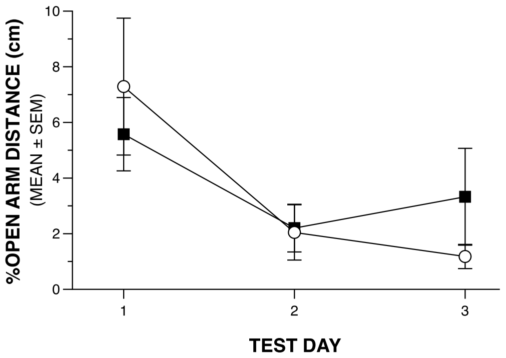


**No significant effects involving Genotype or Sex and no significant comparisons.**

### ELEVATED PLUS MAZE

#### DISTANCE VARIABLES

##### TOTAL DISTANCE TRAVELED

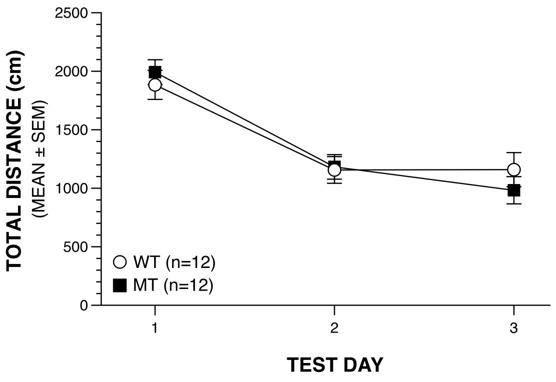


**No significant effects involving Genotype or Sex and no significant comparisons.**

##### DISTANCE TRAVELED IN CLOSED ARMS

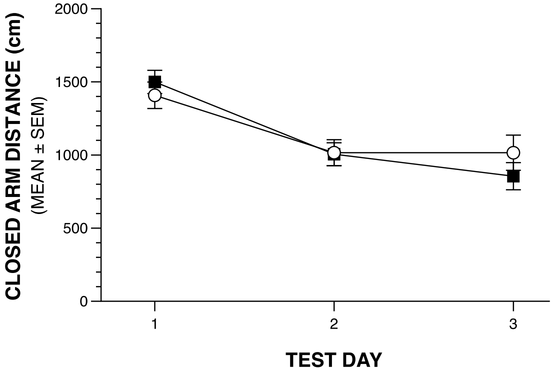


**Significant Genotype x Sex x Test Day interaction: p=.031.**

##### DISTANCE TRAVELED IN CENTER

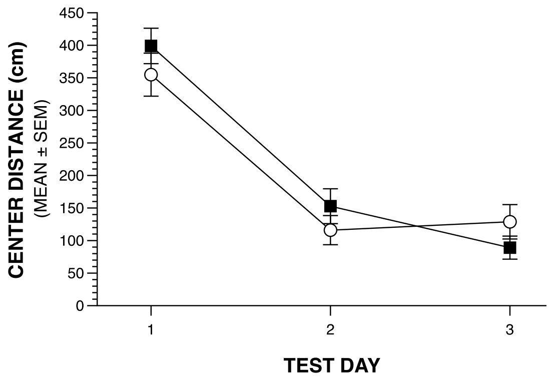


**No significant effects involving Genotype or Sex and no signficant comparisons.**

##### DISTANCE TRAVELED IN OPEN ARMS

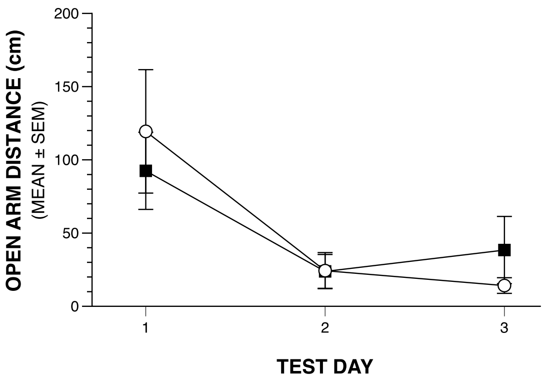


**No significant effects involving Genotype or Sex and no signficant comparisons.**

### CONDITIONED FEAR

#### DAY 1: BASELINE - T/S TRAINING
##### BOTH SEXES COMBINED

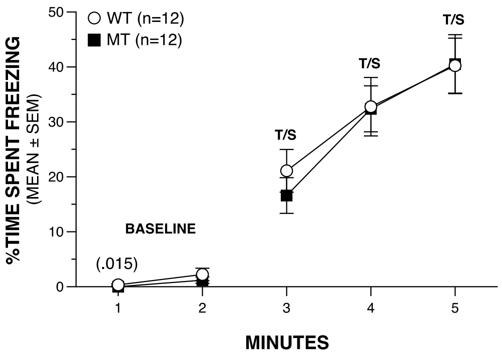


**BASELINE: no significant effects involving Genotype but a significant Sex effect (p=.049).**

**T/S TRAINING: no significant effects involving Genotype or Sex and no significant comparisons.**

#### DAY 2: CONTEXTUAL FEAR
##### BOTH SEXES COMBINED

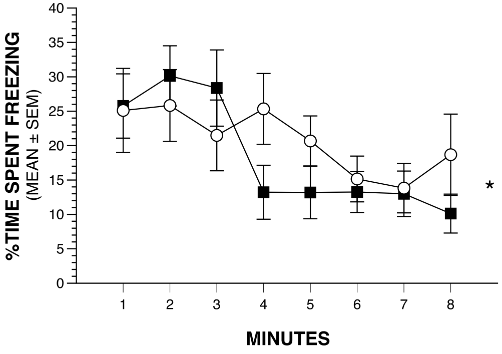


**Significant Genotype x Minutes interaction (*p=.029), and Genotype x Sex x Minutes interaction (p=.017).**

#### DAY 3: AUDITORY CUE
##### BOTH SEXES COMBINED

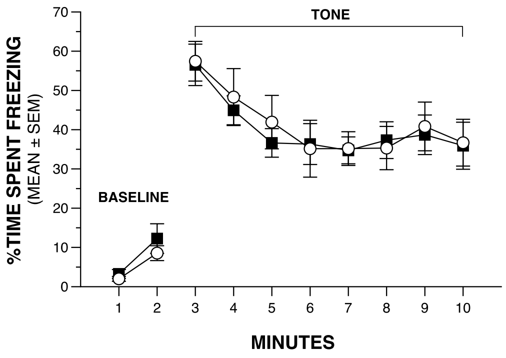


**BASELINE: no significant effects involving Genotype or Sex and no significant comparisons.**

**AUDITORY CUE: no significant effects involving Genotype or Sex and no significant comparisons.**

#### DAY 2: CONTEXTUAL FEAR
##### SEX BY GENOTYPE

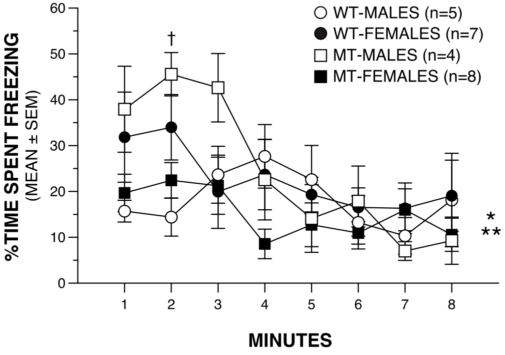


**Significant Genotype x Minutes interaction (*p=.029), and Genotype x Minutes x Sex interaction (**p=.017). WT-Males vs MT-Males: ns; Minute 2 (†p=.002)**

### ACOUSTIC STARTLE/PPI

#### ACOUSTIC STARTLE

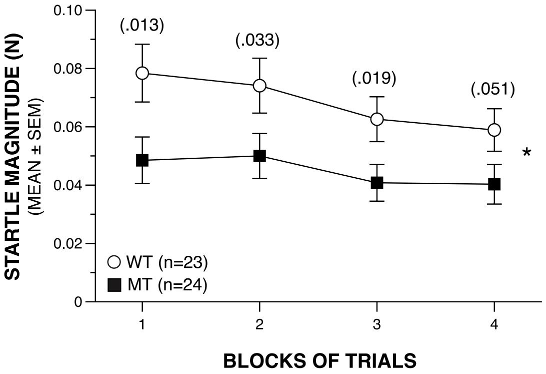


Significant Genotype (*p=.018) and
Sex (p=.004) effects.

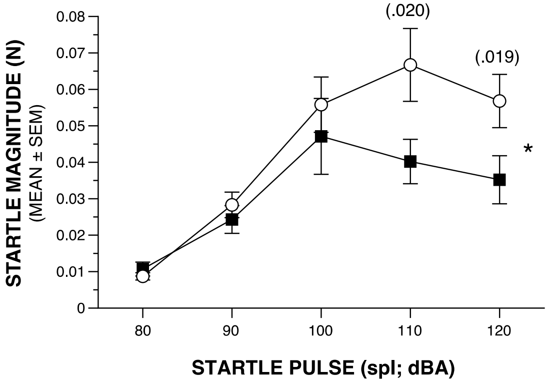


Significant Sex effect (p=.006), and Genotype x
Pulse (*p=.018), and Sex x Pulse (p=.004) interactions.

#### % OF MICE SIGNIFICANTLY STARTLED

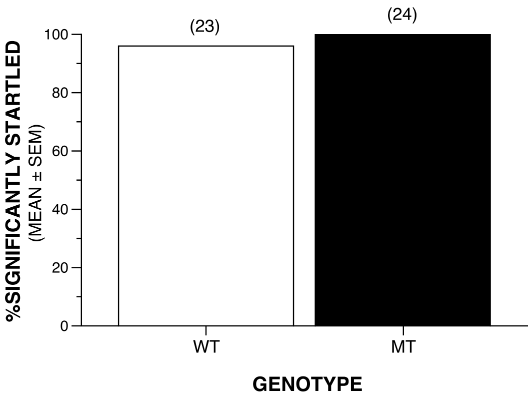


24/24 MT mice and 23/24 WT mice exhibited
a significantly increased startle magnitude
compared to those during no stimulation trials.

#### TOTAL MEAN %PPI

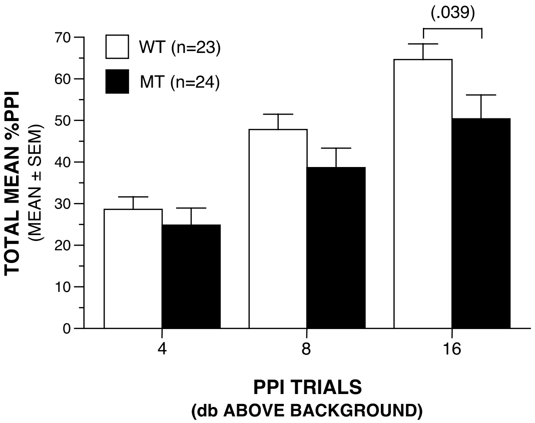


Significant Genotype x PPI trials
interaction: p=.029. Genotype effect: p=.10.

#### SINGLE BLOCK %PPI

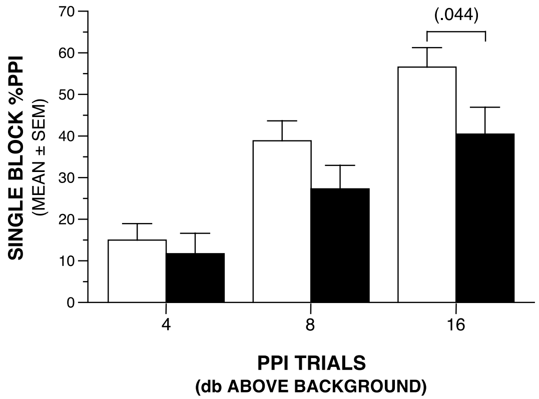


Significant Genotype x PPI trials
interaction: p=.048. Sex effect: p=.051.
Genotype effect: p=.11.

#### BODY WEIGHTS AT TESTING

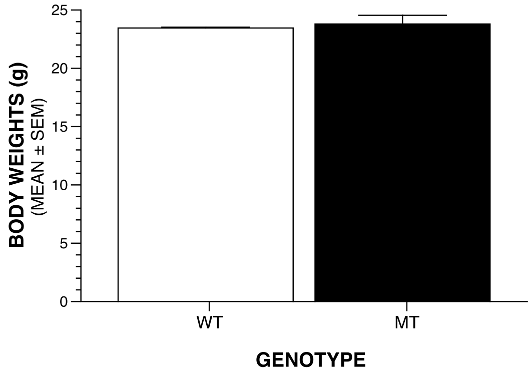


No Significant effect involving Genotype,
but a significant Sex effect (p<.00005) .

### ACOUSTIC STARTLE/PPI

##### ACOUSTIC STARTLE

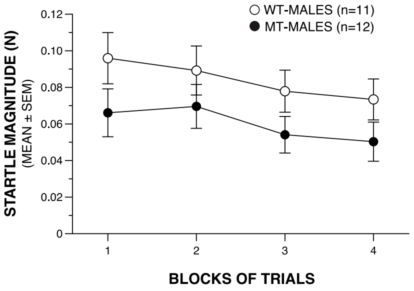


**Significant Genotype (*p=.018) and Sex (p=.004) effects. WT-MALES vs MT-MALES: ns; and no significant between-groups comparisons. Genotype effect: p=.096.**

##### ACOUSTIC STARTLE: INCREASING SPLs

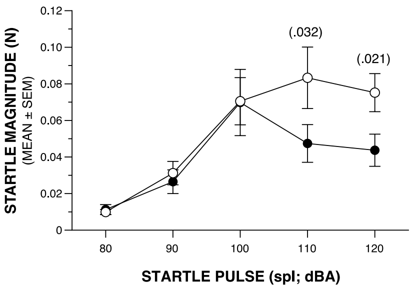


**Significant Sex effect (p=.006), and Genotype x Pulse (*p=.018), and Sex x Pulse (p=.004) interactions. WT-MALES vs MT-MALES: ns.**

##### TOTAL MEAN %PPI

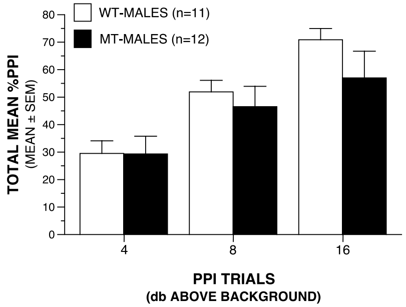


**Significant Genotype x PPI trials interaction: p=.029. WT-MALES vs MT-MALES: ns; and no significant between-groups comparisons.**

##### SINGLE BLOCK %PPI

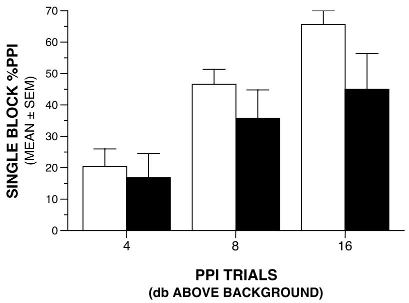


**Significant Genotype x PPI trials interaction: p=.048. Sex effect: p=.051. WT-MALES vs MT-MALES: ns; and no significant between-groups comparisons.**

##### ACOUSTIC STARTLE

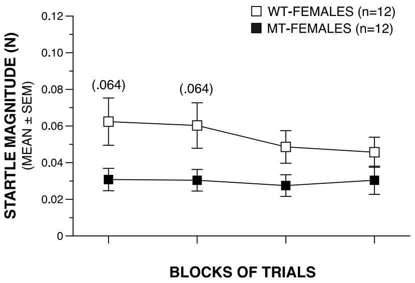


**Significant Genotype (*p=.018) and Sex (p=.004) effects. WT-FEMALES vs MT-FEMALES: ns; and no significant between-groups comparisons. Genotype effect: p=.084.**

##### ACOUSTIC STARTLE: INCREASING SPLs

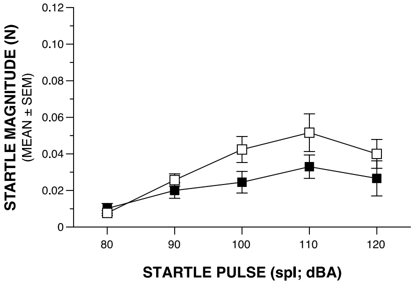


**Significant Sex effect (p=.006), and Genotype x Pulse (*p=.018), and Sex x Pulse (p=.004) interactions. WT-FEMALES vs MT-FEMALES: ns; and no significant between-groups comparisons.**

##### TOTAL MEAN %PPI

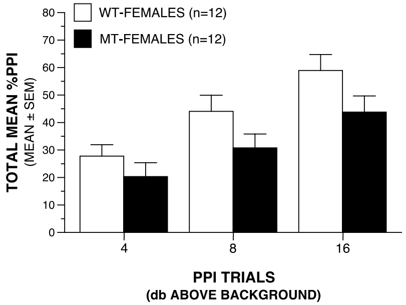


**Significant Genotype x PPI trials interaction: p=.029. WT-FEMALES vs MT-FEMALES: ns; and no significant between-groups comparisons.**

##### SINGLE BLOCK %PPI

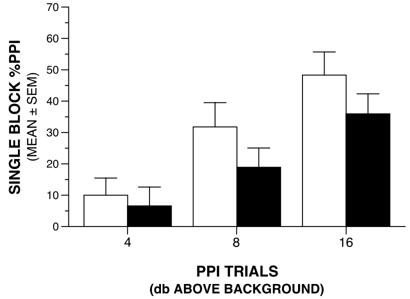


**Significant Genotype x PPI trials interaction: p=.048. Sex effect: p=.051. WT-FEMALES vs MT-FEMALES: ns; and no significant between-groups comparisons.**

#### SOCIAL APPROACH TEST TRIAL - ZONE VARIABLES

##### TIME IN INVESTIGATION ZONE

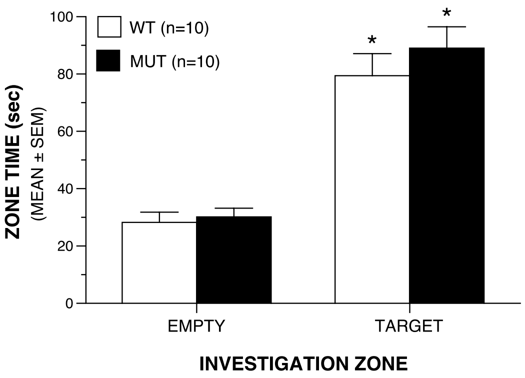


**Significant Genotype x Sex x Zone interaction (p=.044), but no significant between-groups comparisons.**
**WT: Empty vs Target: *p<.00005.**
**MUT: Empty vs Target: *p<.00005.**

##### INVESTIGATION ZONE ENTRIES

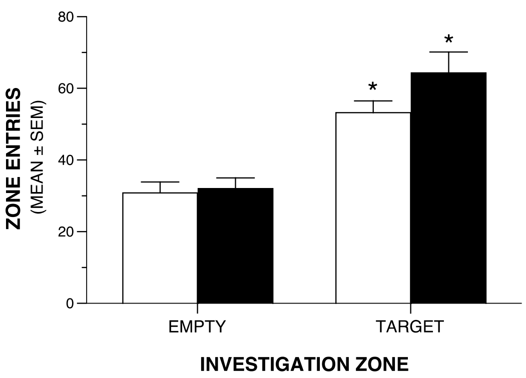


**No significant effects involving Genotype or Sex, and no significant between-groups comparisons.**
**WT: Empty vs Target: *p=.0009.**
**MUT: Empty vs Target: *p<.00005.**

##### LATENCY INTO INVESTIGATION ZONE

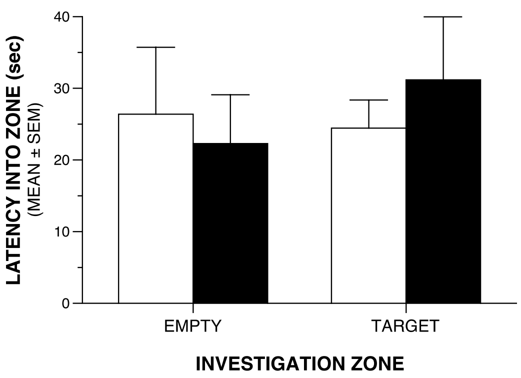


**No significant effects involving Genotype or Sex, and no significant comparisons.**

##### TIME IN INVESTIGATION ZONE

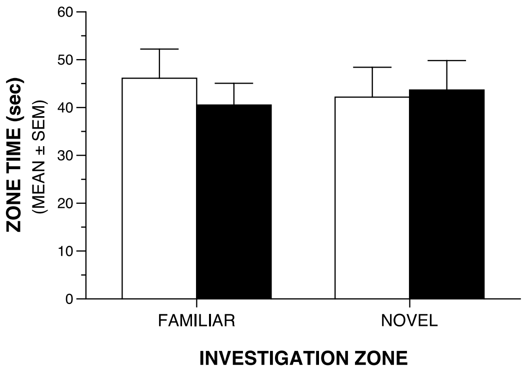


**Significant Genotype x Sex x Zone interaction (p=.0003), but no significant comparisons.**

##### INVESTIGATION ZONE ENTRIES

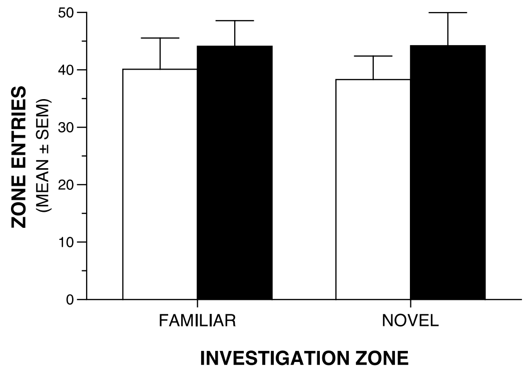


**Significant Genotype x Sex x Zone interaction (p=.0002), but no significant comparisons.**

##### LATENCY INTO INVESTIGATION ZONE

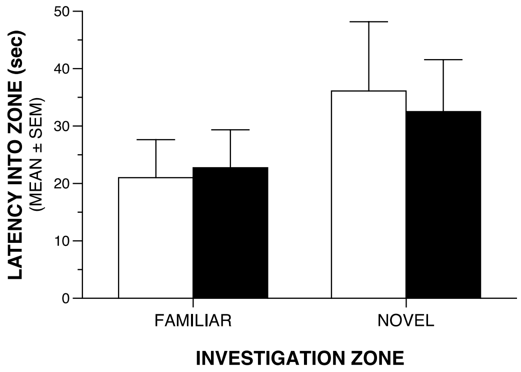


**No significant effects involving Genotype or Sex, and no significant comparisons.**

### SOCIAL APPROACH TEST TRIAL - CHAMBER VARIABLES

#### TIME IN CHAMBERS

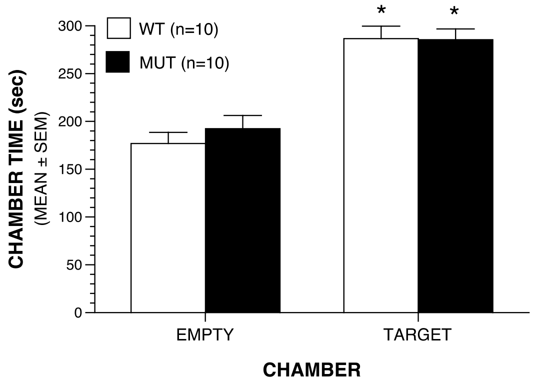


**No significant effects involving Genotype or Sex, and no significant between-groups comparisons.**
**WT: Empty vs Target: *p=.0004.**
**MUT: Empty vs Target: *p=.002.**

#### ENTRIES INTO CHAMBERS

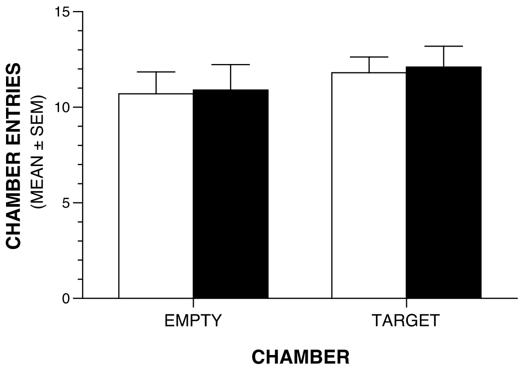


**No significant effects involving Genotype or Sex, and no significant comparisons.**

#### LATENCY INTO CHAMBERS

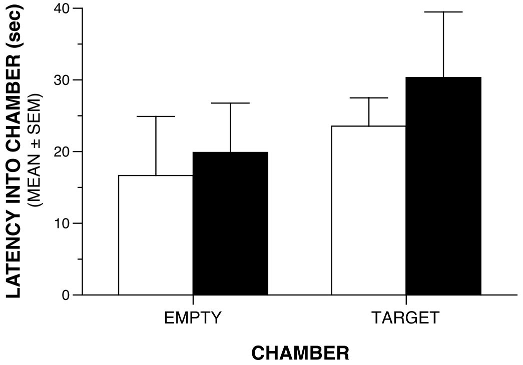


**No significant effects involving Genotype or Sex, and no significant comparisons.**

#### TIME IN CHAMBERS

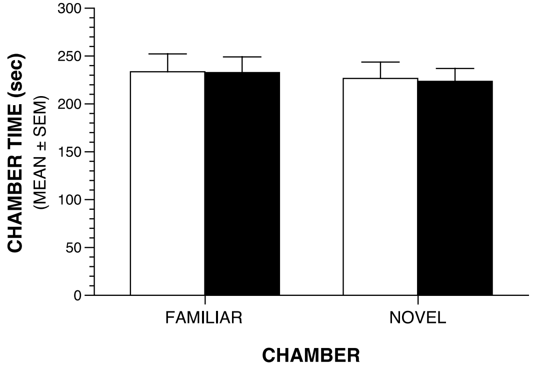


**No significant effects involving Genotype, but a significant Sex x Chamber interaction (p=.009). No significant comparisons.**

#### ENTRIES INTO CHAMBERS

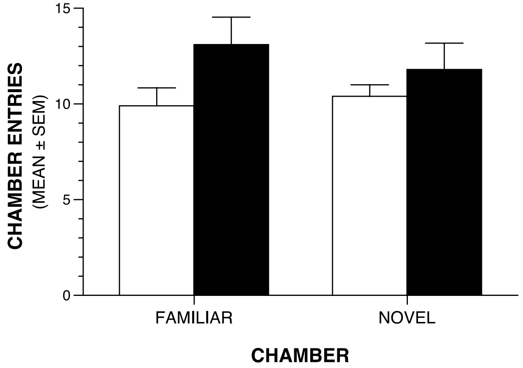


**No significant effects involving Genotype or Sex, and no significant comparisons.**

#### LATENCY INTO CHAMBERS

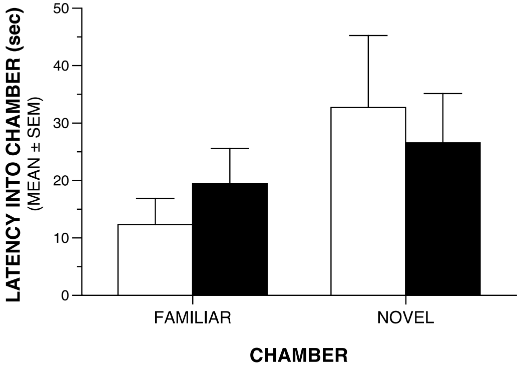


**No significant effects involving Genotype or Sex, and no significant comparisons.**

### TAIL-SUSPENSION TEST

#### TOTAL IMMOBILITY TIME

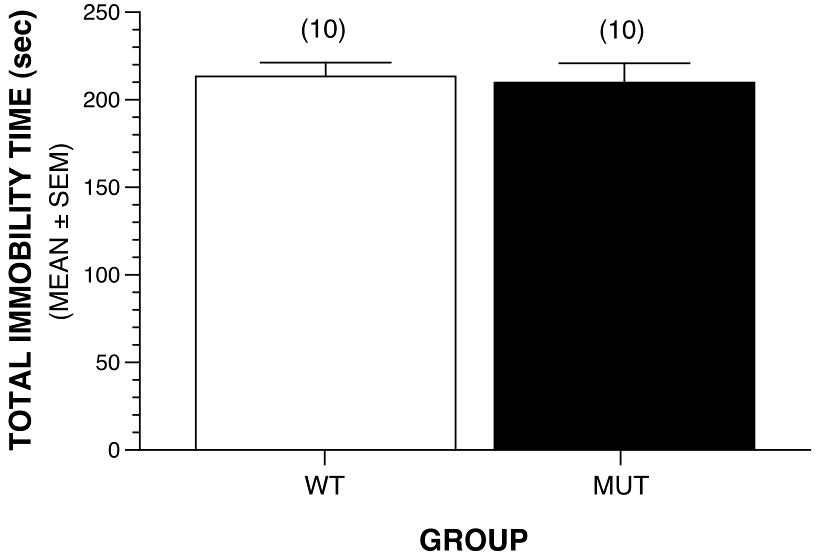


**No significant effects involving Genotype or Sex.**

#### IMMOBILITY ACROSS TIME BLOCKS

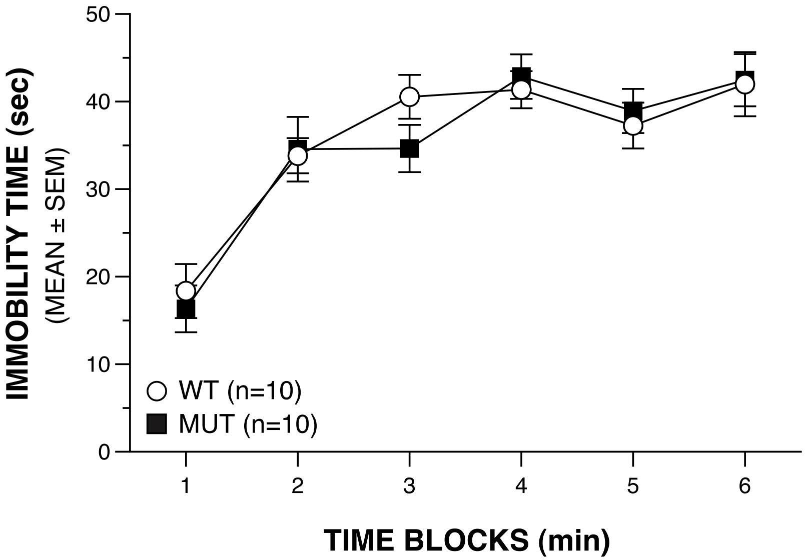


**No significant effects involving Genotype or Sex, and no significant between-groups comparisons.**
